# Technology solutions for immune loci: integrated high-resolution KIR genotyping reveals allelic, structural, and KIR–HLA diversity across 24 human populations

**DOI:** 10.64898/2026.10.22.761185

**Authors:** Chul-Woo Pyo, Ruihan Wang, Alexander Thomas, Yue Song, Cory Friedman, Yunsik Ohm, Akiko Ishitani, Wyatt C Nelson, Daniel E Geraghty

## Abstract

High-resolution genotyping of killer cell immunoglobulin-like receptor (KIR) genes remains challenging because of extensive sequence homology, copy-number variation, and complex haplotype structures. Although KIR allelic variation influences receptor expression and function, routine typing has largely been limited to gene presence or absence. Here, we evaluated an integrated short-read sequencing system combining a four-reaction, two-stage multiplex PCR assay (ScisGo-KIR-v3), centralized cloud-based copy-aware allele analysis, and interactive genotype review. The system was applied to 2,298 individuals representing 24 populations from the 1000 Genomes Project, with matched high-resolution HLA class I genotypes. Independent validation against complete, phased PacBio KIR sequences from 48 samples demonstrated 98.6% automated copy-number concordance and 92.8% combined copy-number and allele concordance across 768 gene–sample comparisons. Following desktop review, all evaluated assignments were concordant with the reference sequences. In a separate comparison of four computational genotyping methods involving 311 shared samples, the automated Scisco pipeline showed the lowest overall mismatch with the four-method consensus (1.62%). Population analyses identified substantial allele diversity and distinct centromeric and telomeric haplotype-motif distributions. Although exact haplotype pairs were frequently ambiguous, structural assignments became increasingly resolvable at the motif level, and motif backgrounds exhibited characteristic allele compositions. Paired KIR and HLA analysis revealed inherited receptor-ligand configurations concealed by gene-content typing. Among 1,401 KIR3DL1/HLA-B Bw4 co-carriers, 57 lacked an allele annotated for ordinary surface expression, while 457 of 1,175 KIR2DS4 co-carriers of six specified HLA-C allotypes carried only truncated receptor forms. Allele-specific KIR2DL1 and KIR2DS5 classifications further distinguished receptor properties and published ligand-binding annotations that varied among populations. Although potential receptor-ligand repertoire composition differed across populations, analyses did not establish preferential KIR-HLA genetic association after accounting for population structure and multiple testing. Together, these findings demonstrate that integrated short-read KIR genotyping can provide scalable, high-resolution allele and copy-number information while explicitly retaining uncertainty in inferred haplotype structures. Compatibility with established HLA laboratory workflows provides a practical foundation for incorporating allele-resolved KIR analysis into immunogenetic research and future clinical investigations.

## 1. Introduction

Human leukocyte antigen (HLA) class I molecules and the killer cell immunoglobulin-like receptors (KIR) together form a key interface between innate and adaptive immunity (1, 2). Besides presenting peptides to CD8⁺ T cells, HLA class I allotypes carrying defined epitopes (C1, C2, Bw4 and certain HLA-A allotypes) are ligands for KIR expressed on natural killer (NK) cells and subsets of T cells. Their interactions set the balance between NK cell inhibition and activation (3, 4). The KIR locus varies in two dimensions. First, haplotypes differ in gene content and copy number: up to 15 genes and 2 pseudogenes are arranged in A and B haplotypes. These haplotypes are built from distinct centromeric and telomeric segments, and recombination between them has expanded and contracted the locus (5–9). Second, most KIR genes are highly polymorphic at the allele level (10–12). Large sequencing studies have shown that this allelic diversity is broad but unevenly distributed across loci and populations (13, 14). Taken together, the population diversity of the region rivals or exceeds that of HLA. Because of this structural variability and high sequence identity between paralogous genes, KIR is among the most difficult regions of the genome to genotype accurately.

Since KIR A and B haplotypes were first described nearly 30 years ago (15, 16), KIR variation and KIR–HLA combinations have been associated with infectious disease, autoimmunity, pregnancy disorders, cancer and transplantation outcomes (reviewed in 2, 17–19). Almost all of these studies have defined KIR by gene presence or absence, or by haplotype content. Allelic variation has largely been ignored because practical methods to resolve it were lacking.

This omission matters because KIR alleles are not functionally equivalent. KIR3DL1 allotypes range from high to low surface expression, and some are retained inside the cell (20). Specific combinations of KIR3DL1 allotypes and HLA-B Bw4 ligands are associated with differences in HIV disease progression that gene-level data do not capture (21). KIR2DL1 allotypes differ in inhibitory strength, and donor KIR2DL1 alleles have been associated with outcomes after allogeneic transplantation (22). Likewise, KIR3DL1 and HLA-B subtype combinations have been associated with acute myeloid leukemia relapse (23). The most common KIR2DS4 alleles carry a deletion that removes the receptor from the cell surface (10). Most recently, adding high-resolution KIR genotyping improved the prediction of outcomes after hematopoietic stem cell transplantation (HSCT) (24). In each of these examples, gene-level typing groups together alleles with different functions, which can weaken or hide real associations.

Several approaches to allele-level KIR genotyping have been described (reviewed in 25). Short-read laboratory methods include target capture with *HLA-A*, *-B* and *-C* (13), long-range PCR with Illumina sequencing (26, 27), and exon-targeted, copy-aware amplicon typing, which has been applied to more than a million registry donors (28). Long-read methods phase complete haplotypes (29–32) but make greater demands on infrastructure and throughput. Computational tools recover KIR genotypes from existing data by imputation from SNP arrays (33), from short-read whole-genome or exome sequence with PING (34), T1K (35) and kir-mapper (36), or by long-read assembly (37). Because paralogy and exon sharing vary between KIR genes, the accuracy of all of these approaches is likely to vary by locus.

Allele-level KIR typing is still rarely used in routine clinical laboratories, where gene-content typing by PCR-SSP, SSO or real-time PCR remains standard. The main exception is registry-scale donor typing (28). Two factors probably explain this. First, evidence that KIR information should guide clinical decisions remains mixed. Donor selection based on KIR B-haplotype content has been associated with reduced relapse after HSCT for acute myeloid leukemia (38), whereas other haplotype-based models have failed to predict outcome (39, 40). Gene-level typing may be one reason the results are inconsistent. Second, most allele-level methods require specialized instruments, long-range amplification or local bioinformatic expertise that clinical HLA laboratories rarely have. A method that couples a simple bench process with standardized, centrally executed analysis would remove much of this barrier.

The 1000 Genomes Project panel is an open, ancestrally diverse reference resource and a natural benchmark for KIR typing. Analysis of a subset of KIR genes in this panel has already revealed substantial previously uncharacterized sequence and structural diversity (13). Genotypes from kir-mapper, PING and T1K are available for overlapping samples (36), and complete KIR haplotype sequences exist for a further subset. Together these allow a targeted assay to be compared directly with independent methods on the same individuals. However, no targeted laboratory assay has yet produced allele-level KIR genotypes and haplotype structures for the full panel and paired them with high-resolution HLA typing. Short-read data also cannot fully phase the KIR locus, so a population-scale analysis needs a structural interpretation that states openly where haplotype assignments are ambiguous rather than forcing a single assignment.

Here we describe a commercially available, amplicon-based short-read KIR typing kit (ScisGo-KIR-v3) that resolves both haplotype structure and full-exon allele genotypes at all 17 KIR loci. It combines a four-reaction, two-stage multiplex PCR with centralized cloud-based, copy-aware analysis and interactive review of calls, and uses the same bench workflow as an established clinical HLA typing system. We applied it to 2,298 individuals from 24 populations of the 1000 Genomes panel, for which HLA genotypes at 3-field resolution were determined separately (Pyo et al., in preparation). We first assess accuracy, comparing calls with complete phased KIR haplotype sequences determined independently for 48 samples and benchmarking allele-level concordance against kir-mapper, PING and T1K on shared samples. We then describe KIR allele diversity across the 24 populations and, using ambiguity-aware inference, characterize centromeric and telomeric haplotype structure together with the alleles carried on each structural background. Pairing these calls with HLA data from the same samples, we identify inherited receptor–ligand configurations that gene-level KIR typing conceals, summarize the potential receptor–ligand repertoires they generate, and test for preferential receptor–ligand association both as frequency covariation between populations and as genetic co-carriage within them. Finally, we compare the laboratory and analytical demands of the KIR workflow with those of the established HLA workflow.

## 2. Materials and Methods

### 2.1 KIR allele typing with the ScisGo-KIR-v3 kit

KIR genotyping was performed with a commercially available targeted amplicon kit, ScisGo-KIR-v3 (Scisco Genetics, Inc., Seattle WA), following the manufacturer’s protocol. The kit is a unified KIR typing system that generates gene-detection, copy-number, structural, and allele-level outputs within a single assay-and-analysis framework (Figure 1), combining multiplex PCR, Illumina sequencing, centralized cloud-based analysis, and interactive review of final calls. Four multiplex reactions are run in a two-stage PCR so that exon-level evidence across all KIR genes is generated in parallel rather than through serial locus-specific assays. In stage 1, locus-targeting amplification enriches KIR exon-containing regions across the locus. In stage 2, sequencing-compatible products are generated directly from the stage 1 amplicons without transfer between wells, which reduces sample handling and supports batch processing.

**Figure 1.**
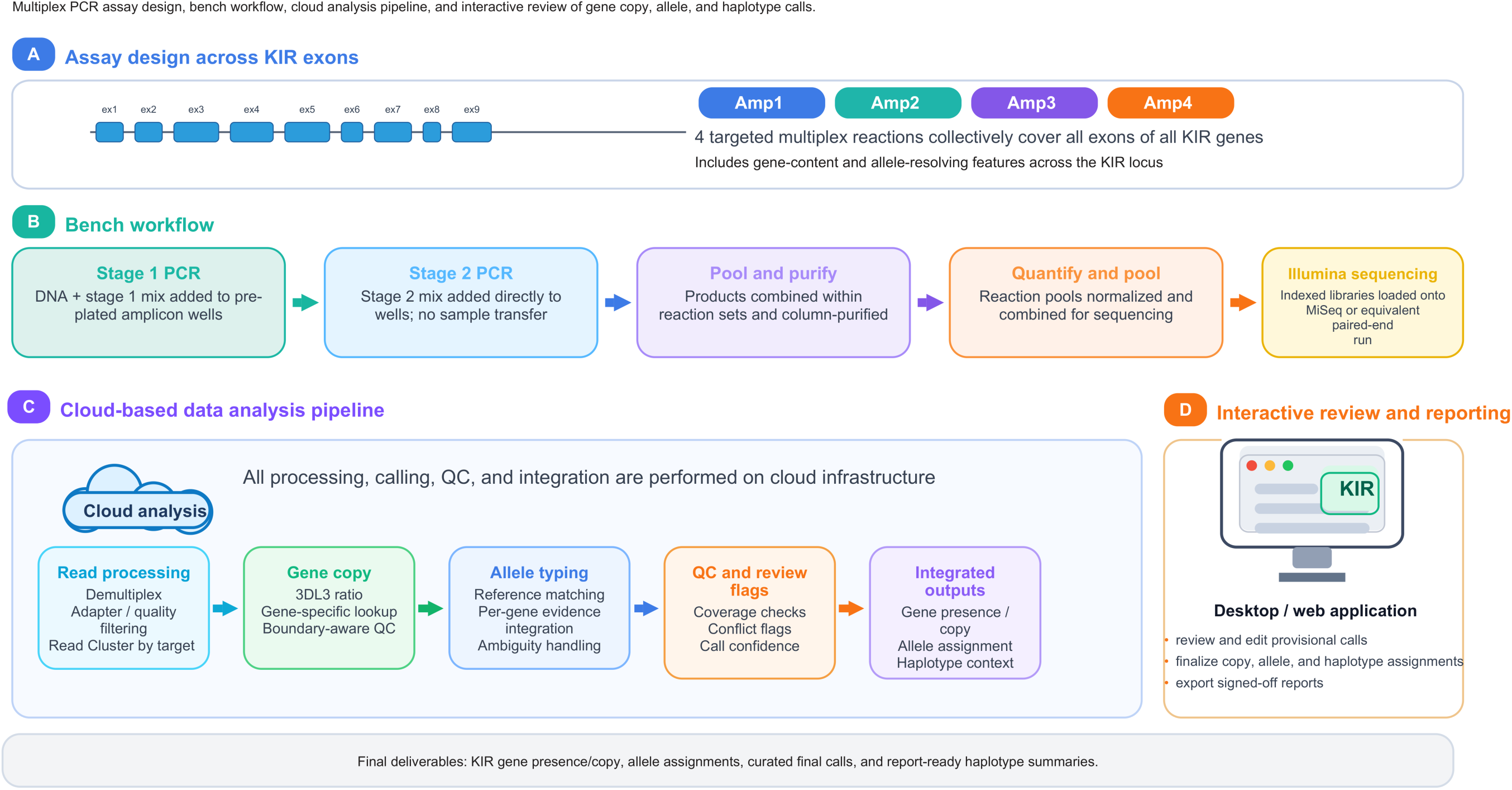
KIR high-resolution allele typing workflow. Schematic overview of the copy-aware KIR typing system used in this study. Panel A shows assay design across KIR exons. Panel B summarizes the two-stage multiplex bench workflow, in which four coordinated reactions are pooled for sequencing with limited sample handling. Panel C shows the cloud-based analysis pipeline, where shared reference resources, quality-control rules, copy-number determination, and copy-aware allele assignment are applied uniformly across batches. Panel D shows the review and reporting environment used for final curation and export of gene, copy, structural, and allele outputs.

Products from the four reactions were combined within reaction sets, column-purified, quantified, normalized, and sequenced on an Illumina NextSeq, MiSeq and MiSeq i100 (Illumina, San Diego CA) with paired-end reads and batch sizes determined by the sequencing platform (2 × 300 bp on NextSeq 2000 P1 Reagents for eight 96-sample plates; 2 × 250 bp on MiSeq Kit v2 on two 96-sample plates and 2 × 300 bp on MiSeq i100 25M Reagent Kit for four 96-sample plates). Sequence data were analyzed with the Scisco cloud-based analysis service (Sciscloud.net, workflow KIR-Allele 0.1.37; Scisco Genetics, Inc., Seattle WA), which applies the same reference resources, quality-control rules and interpretation logic to all runs for raw-data preprocessing, copy-number determination, and copy-aware allele assignment. Final calls were reviewed and finalized in the GeMS-KIR12 desktop application (Scisco Genetics, Inc., Seattle WA), which supports interactive inspection of supporting evidence, targeted editing of provisional assignments, and export of curated report-ready outputs.

The GeMS-KIR12 application provides an interactive environment for evaluating sequence evidence, resolving ambiguous allele assignments, reconciling copy-number and allele calls, and reviewing predicted KIR haplotype structures. The application includes multiple editing and interpretation functions that support expert review of complex KIR genotypes. Detailed descriptions of these functions, including their operation and application to genotype review, are provided in the Scisco Genetics GeMS-KIR12 User Manual (Scisco Genetics, Inc., Seattle, WA). Final allele and copy-number assignments were exported following review.

### 2.2 KIR gene copy-number determination

KIR gene copy determination was implemented within the allele-typing pipeline (Figure S1). KIR3DL3 was assumed to be present at two copies in each sample and served as the internal dosage reference for target-gene copy-number determination. For each informative exon, the target-gene read count was divided by the corresponding KIR3DL3 exon read count. These exon-specific ratios were averaged to obtain a gene-level normalized ratio.

Integer copy number was assigned by comparing the normalized ratio with gene-specific lookup intervals established from 408 control samples with known copy number (Table S10). Gene-specific rather than global thresholds were used because paralogy and structural variation across the KIR region cause amplification and read-mapping efficiency to differ among genes and exons (5, 34). Following filtering of the control ratios, boundaries between adjacent copy-number classes were calculated as the midpoint between the 75th percentile of the lower-copy class and the 25th percentile of the higher-copy class. Final gene presence or absence and integer copy calls were used as constraints in allele assignment and haplotype compatibility matching.

### 2.3 Allele determination and the allele reference set

Following read preprocessing and copy determination, allele assignment was performed against an allele reference set that combined curated KIR allele sequences from IPD-KIR (11) with full-length KIR gene and haplotype sequences generated in this laboratory by long-read sequencing of 1000 Genomes samples (6, 7, 9, 29). A total of 109 KIR haplotypes from 1000 Genomes Project samples were independently sequenced to completion using PacBio long-read sequencing technology. These data provided fully phased sequences of the KIR haplotypes and their component genes, allowing direct determination of gene content, copy number, allele identity, and haplotype structure from the complete sequences. Forty-eight of the samples represented in this long-read sequence dataset were selected for validation of the ScisGo-KIR-v3 short-read typing system, as described in Section 2.6. The complete long-read sequences provided reference genotypes and haplotype structures against which the independently generated short-read results could be evaluated.

### 2.4 Study cohort and dataset

The workflow was applied to 2,298 individuals from the 1000 Genomes Project Phase 3 panel, organized into 24 populations grouped into five continental super-populations: African (AFR, *n* = 709), Admixed American (AMR, *n* = 215), East Asian (EAS, *n* = 416), European (EUR, *n* = 414), and South Asian (SAS, *n* = 544) (Figure S2A–B; Table S1). Analyses were performed across 17 KIR loci: KIR2DL1, KIR2DL2, KIR2DL3, KIR2DL4, KIR2DL5A, KIR2DL5B, KIR2DP1, KIR2DS1, KIR2DS2, KIR2DS3, KIR2DS4, KIR2DS5, KIR3DL1, KIR3DL2, KIR3DL3, KIR3DP1, and KIR3DS1. Allele nomenclature followed the latest IPD-KIR reference nomenclature (11). The dataset contained diploid copy and allele calls for each individual and locus, enabling direct allele-frequency analysis and copy-aware structural summarization. Primarily, we used NextSeq platforms in three runs to generate most of the data. Around 1–2% of samples produced failed/weak data in each NextSeq run. We used MiSeq or MiSeq i100 to recover the data and to validate primer batches. Using the NextSeq platform, batches of eight 96-sample plates (768 samples) were processed from PCR initiation to sequencer loading within three days. Smaller batches of one or two 96-sample plates were processed using the MiSeq or MiSeq i100 platforms within two days.

### 2.5 Allele frequency estimation, filtering, population differentiation, and dominant-allele summaries

For each locus, allele frequencies were estimated within each analysis unit (overall cohort, super-population, and population) from all observed allele calls in all three typing slots. The denominator was defined as the total number of non-missing allele copies within the unit at that locus rather than a fixed 2N diploid count, thereby accommodating loci with copy-number variation. For visualization of allele richness, a minimum-count filter of five observed allele copies per analysis unit was applied to reduce inflation by single-observation artefacts. Dominant alleles were defined as the most frequent alleles within each analysis unit. Between-population allele-distribution differences were summarized by Cramér’s V from population-by-allele count tables (41). The eight most frequent alleles overall were retained separately at each locus and remaining alleles were pooled as Other. V was calculated as the square root of χ^2^/[n × min(r − 1, c − 1)], where n is the total allele count and r and c are the numbers of populated rows and columns. This descriptive statistic was evaluated across 24 populations and, separately, five super-populations; it was not treated as an estimate of FST or as a person-level association test. Figure 2A shows the 24-population values. Super-population allele-frequency heatmaps and dominant-allele summaries were generated for four selected inhibitory or framework loci (Figure 2B–F).

**Figure 2.**
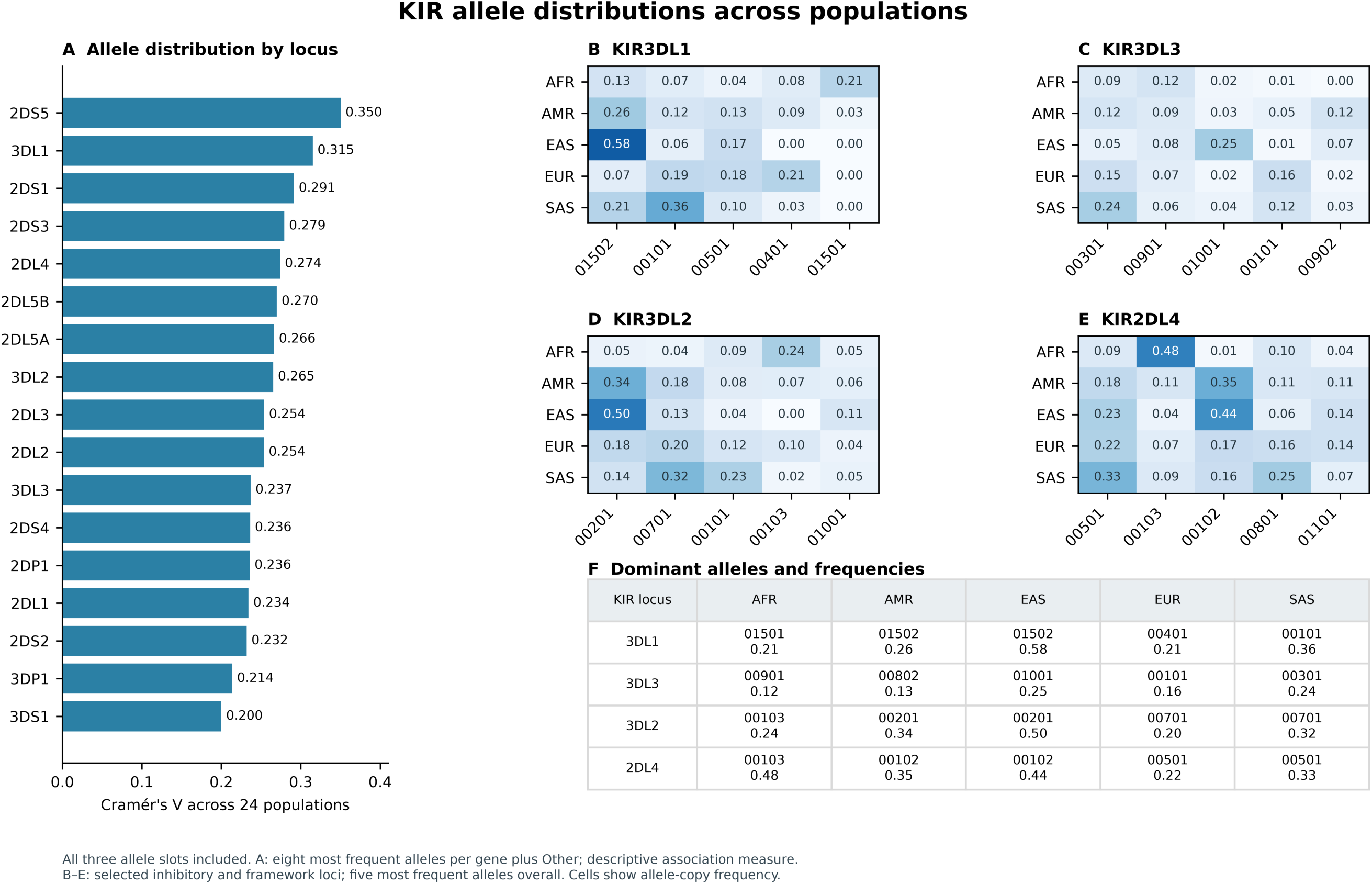
Population differentiation and dominant KIR alleles. Panel A shows descriptive Cramér’s V across 24 populations for 17 KIR loci, using the eight most frequent alleles per locus and a pooled Other category. Panels B–E show allele-copy frequencies across five super-populations for the five most frequent overall alleles at KIR3DL1, KIR3DL3, KIR3DL2 and KIR2DL4, respectively. These four loci were selected to illustrate inhibitory and framework allele profiles. Panel F gives the dominant allele and its frequency in each super-population at the same four loci. All three typing slots were included; denominators are observed allele copies.

### 2.6 Cross-method comparison

The performance of the Scisco KIR-Allele analysis software in determining KIR gene copy number and allele assignments from short-read sequence data was compared with three independently developed KIR genotyping methods: KIR-MAPPER, PING, and T1K. The comparison used 311 overlapping samples with callable results across all four methods, integrating the current workflow with previously reported short-read KIR analyses (36). Scisco calls were taken directly from the automated KIR-Allele pipeline before GeMS-KIR12 desktop review. Per-gene mismatch rates were calculated relative to a four-method consensus (Table 1). The methods employ distinct computational approaches: PING incorporates genotype-aware analysis and copy-number determination (34), T1K performs joint inference across homologous genes using whole-genome, whole-exome, or RNA-seq data (35), and KIR-MAPPER provides allele, copy-number, and variant detection within a reference-genome framework (36). Ambiguous PING calls were counted as matches when the consensus allele pair was included among the reported candidates. Additional comparisons and supporting records are provided in Table S3.

**Table 1.** Comparison of automated KIR genotyping software. Per-gene mismatch rates (%) relative to the four-method consensus across n = 311 shared samples.

| Gene | Current workflow (%) | KIR-MAPPER (%) | PING adjusted (%) | PING ambiguous (%) | T1K (%) |
| --- | --- | --- | --- | --- | --- |
| 2DL1 | 1.68 | 17.51 | 13.47 | 36.70 | 9.76 |
| 2DL2 | 0.00 | 2.41 | 12.05 | 0.00 | 9.64 |
| 2DL3 | 1.09 | 3.65 | 5.47 | 36.13 | 7.66 |
| 2DL4 | 0.66 | 1.98 | 7.59 | 14.19 | 12.87 |
| 2DL5 | 4.29 | 2.45 | 3.07 | 65.03 | 27.61 |
| 2DP1 | 1.01 | 6.40 | 9.76 | 9.09 | 12.12 |
| 2DS1 | 2.59 | 1.72 | 11.21 | 44.83 | 6.03 |
| 2DS2 | 1.29 | 1.94 | 7.74 | 63.87 | 8.39 |
| 2DS3 | 3.75 | 5.00 | 17.50 | 40.00 | 16.25 |
| 2DS4 | 0.34 | 2.41 | 9.66 | 26.55 | 5.86 |
| 2DS5 | 0.00 | 1.77 | 6.19 | 50.44 | 18.58 |
| 3DL1 | 0.00 | 4.08 | 12.59 | 8.84 | 5.78 |
| 3DL2 | 0.00 | 4.19 | 6.45 | 43.87 | 38.06 |
| 3DL3 | 1.97 | 2.62 | 4.92 | 42.95 | 60.33 |
| 3DP1 | 7.26 | 14.52 | 4.29 | 17.16 | 3.63 |
| 3DS1 | 0.00 | 0.00 | 9.57 | 51.06 | 6.38 |
| **Total** | **1.62** | **4.54** | **8.85** | **34.42** | **15.56** |

*Adjusted PING treats ambiguous calls containing the consensus allele as matches.*

Independent reference validation was performed using 48 samples for which complete, phased KIR gene and haplotype sequences had previously been determined by PacBio long-read sequencing. These sequences provided reference assignments of gene content, copy number, allele identity, and haplotype structure. The same samples were genotyped independently using the ScisGo-KIR-v3 short-read assay, with automated calls generated by the Sciscloud KIR-Allele workflow and subsequently reviewed and finalized using GeMS-KIR12. Both provisional and final allele and copy-number assignments were compared with the PacBio-derived references, allowing separate evaluation of automated calling and the complete assay-and-review workflow. The long-read sequences of these 48 samples were excluded from the allele reference set described in Section 2.3 and were unavailable to the reviewer during editing, ensuring that the validation was independent of the reference sequences used for allele assignment and of prior knowledge of the reference genotypes. Concordance was evaluated across 16 KIR gene categories, with KIR2DL5 combined, and included gene absence in the denominator. Copy-number agreement and combined copy-number/allele agreement were assessed separately using the normalization and ambiguity rules described in the supporting comparison workbook (Table S11).

### 2.7 Haplotype structure rule set and ambiguity handling

Haplotype structure was assigned using a second, independent rule set describing gene-content haplotypes. This rule set is distinct from the allele reference set of Section 2.3: it is not curated by IPD-KIR or IMGT, and it governs compatibility of multi-locus gene-content structures, not allele identity. Haplotype entries are reported in centromeric and telomeric motif nomenclature, each carrying a centromeric designation (cA01, cB01, cB02 or cB03) and a telomeric designation (tA01 or tB01), together with any gene-content variant at the 2DS3/2DS5 branch positions, established by the previous studies (6, 7). In the supplied haplotype-list-v3 catalog, an exact label such as cA01|tA01 identifies the cA01 centromeric and tA01 telomeric combination; suffixes distinguish gene-content variants, which were mapped to their regional motif designations before collapse. Each haplotype was represented as a copy-count vector across 15 informative KIR genes: 2DL1, 2DL2, 2DL3, 2DL4, 2DL5, 2DP1, 2DS1, 2DS2, 2DS3, 2DS4, 2DS5, 3DL1, 3DL2, 3DP1, and 3DS1. KIR3DL3 was excluded from compatibility matching because it is a framework gene present in all individuals, and KIR2DL5A and KIR2DL5B were combined as total KIR2DL5 copy number to conform to the structure of the rule set.

For each individual, all unordered pairs of haplotype entries, including homozygous combinations, were enumerated and compared with the observed gene-copy profile. A haplotype pair was considered compatible only when the element-wise sum of the two haplotype vectors exactly matched the observed copy counts across all 15 genes. Deterministic first-match assignment was not used in the primary structural analyses. Instead, all compatible exact haplotype pairs were retained and propagated forward so that structural ambiguity remained explicit throughout downstream summarization and interpretation.

### 2.8 Weighted haplotype-combination and motif frequency estimation

Haplotype frequencies were estimated by retaining all compatible exact haplotype pairs rather than selecting a single solution for ambiguous individuals. Each individual contributed equal total weight, distributed uniformly among their compatible pairs. Exact haplotypes were subsequently collapsed into centromeric (cA01, cB01, cB02 or cB03), telomeric (tA01 or tB01), and combined centromeric–telomeric motif categories before frequency estimation. The cA01|tA01 combination corresponds to the conventional A haplotype; all other combinations were classified as B-associated (5–8).

For descriptive summaries of exact haplotype architectures (Figure S4; Table S5), compatible preferred solutions were selected using the supplied haplotype preference catalog, with equal weighting among retained solutions. This preference procedure was not applied to the primary collapsed-motif frequencies or motif-associated allele analyses. Detailed selection rules are provided in the Supplementary Methods.

### 2.9 Motif-associated allele analysis

Motif-associated allele frequencies were estimated from individuals whose compatible haplotype solutions consistently supported the same homozygous centromeric or telomeric motif pair. Allele copies were counted separately for each gene within its predefined motif-specific gene set (Table S7), with gene-specific denominators reflecting the number of contributing carriers. KIR2DS3 and KIR2DS5 were counted individually at their variable branch positions without assigning these genes to a specific chromosome. The complete motif-specific gene definitions and allele distributions are provided in Table S7. The fixed top-two allele definitions and super-population plotting data for Figure S6 are provided in Table S8.

### 2.10 KIR and HLA allele resolution and functional annotation

KIR and HLA genotypes were matched by sample identifier across 2,298 individuals. HLA-A, -B and -C genotypes were generated separately using the established HLA workflow (Pyo et al., in preparation). Core paired analyses required complete and interpretable HLA-A, -B and -C epitope assignments. One individual (NA20861) lacked complete HLA class I typing and was excluded, leaving 2,297 individuals in the core paired cohort. Availability at finer allele resolutions was assessed separately for each comparison (Table S9I–L).

KIR genotypes were analyzed at gene-presence, copy-number, protein-group and full numeric allele resolution, and HLA genotypes at first-, second- and highest available field resolution. HLA-C1/C2, HLA-B Bw4/Bw6 (including Bw4-80I/80T), and HLA-A Bw4 were assigned from reference sequence residues. Published functional annotations included KIR3DL1 surface-expression classes (42, 43), KIR2DL1 residue-245 variants (44), KIR2DS4 full-length and truncated forms (45), and KIR2DS5 allotypes with reported C2 binding (46). Allele-specific, sequence-inferred and unresolved annotations were distinguished.

Functional categories were summarized among individuals carrying both the specified KIR gene and HLA ligand, retaining uncertain annotations in the denominator. Copy number and compatible structural motifs were included as genomic context. Pointwise Wilson intervals were calculated for descriptive proportions (47).

Potential receptor–ligand repertoires were estimated by counting distinct KIR-protein/HLA-protein combinations using published specificity rules (48–50). Synonymous alleles and repeated copies encoding identical proteins were counted once; null or unavailable receptors were excluded. Unresolved assignments contributed only to an upper annotation scenario. These sequence-derived repertoires represent potential interactions rather than measured receptor activity. Detailed rules are provided in Supplementary Methods S17 and S20.

### 2.11 Between-population correlations of KIR and HLA frequencies

Between-population covariation was evaluated across 24 equally weighted populations using 17 KIR gene-level and six allele-class receptor–ligand comparisons, following the approach of Single et al. (51). KIR copy intensity and HLA ligand frequencies were calculated as copy counts divided by 2N. Raw correlations were evaluated alongside genomic calibration using 2,164 individuals matched to the high-coverage 1000 Genomes dataset (52). Of the 2,297 core individuals, 133 had no matched genotypes in this public panel, leaving 2,164 individuals eligible for genomic calibration before comparison-specific annotation filters. The 133 individuals remained in the full-core analyses; missing public genotypes were not imputed (Table S9AO).

Population covariance was estimated using 3,436 intergenic SNPs from 20 autosomes excluding chromosomes 6 and 19. Frequency contrasts were covariance-adjusted and evaluated against chromosome-separated, frequency-matched genomic reference distributions. Empirical p values were derived from the more conservative of two calibration tails and adjusted across the 23 comparisons using the Holm procedure. Additional calibration, pedigree and sensitivity analyses are described in Supplementary Methods S18–S19.

### 2.12 KIR and HLA co-carriage within populations

Within-population genetic co-carriage was evaluated separately from functional annotation and between-population frequency covariation. Six targeted tests included four gene-level receptor–ligand combinations (KIR2DL1/C2, KIR2DL2/C1, KIR2DL3/C1 and KIR3DL1/HLA-B Bw4) and two allele-function comparisons involving KIR3DL1 expression and KIR2DL1 residue 245, conditional on receptor copy number (43, 53).

Analyses were stratified by the 24 constituent populations. Gene-level associations were estimated using Mantel–Haenszel odds ratios, with 5,000 within-population bootstrap replicates for confidence intervals and 100,000 randomizations for targeted tests. The six targeted comparisons were adjusted together using the Holm procedure (54).

Exploratory analyses examined increasing KIR and HLA allele resolution and regional KIR gene combinations while preserving linked KIR genotypes and randomizing complete HLA-A/B/C genotypes within populations. Conditional analyses also accounted for regional motif and copy-number or protein-genotype backgrounds. The 72 regional tests were evaluated using 99,999 randomizations and Holm correction; additional exploratory testing families used Benjamini– Hochberg (55) or Benjamini–Yekutieli (56) adjustment as specified. Detailed test definitions, stratification procedures and sensitivity analyses are provided in the Supplementary Methods and Table S9.

### 2.13 Software, code and data availability

Genotype calling and review were performed using the Scisco software described in Section 2.1. Subsequent analyses and figure generation used custom Python scripts, with JavaScript used to assemble supplementary workbooks. Analysis code, dependencies and instructions are provided in Supplementary Code 1. Sample-level KIR and HLA genotypes, population annotations, haplotype catalogs and supporting data are provided in Supplementary Data 1.

## 3. Results

Results are presented in four parts. Allele and copy-number calls were first compared with complete phased haplotype sequences in a set of reference samples, and then benchmarked against three independent short-read methods on a shared set of samples. KIR allele diversity was then described across the 24 populations. Haplotype structure was resolved as far as short-read data allow — to centromeric and telomeric motif backgrounds rather than to exact haplotypes — and those backgrounds were used to link structure to allele content. Finally, paired HLA typing was used to identify inherited receptor–ligand configurations that gene-level KIR calls conceal, to test whether receptor and ligand frequencies covary across populations, and to evaluate genetic co-carriage within them.

### 3.1 Calls matched complete phased haplotype sequences in 48 reference samples

Reference comparison comprised 768 gene–sample combinations across 48 samples and 16 KIR gene categories. In the provisional software output, copy number agreed with the reference in 757/768 comparisons (98.6%), and both copy number and the allele combination agreed in 713/768 (92.8%). The lowest provisional concordance occurred at KIR3DP1 (35/48, 72.9%), KIR2DP1 (39/48, 81.3%) and KIR2DL1 (41/48, 85.4%), corresponding to 13, nine and seven discordant calls, respectively. Of the 55 provisional discordances, 11 involved copy number and 44 involved allele assignment despite matching copy number. KIR2DL3, KIR2DS1, KIR2DS3 and KIR3DS1 were concordant in all 48 samples before review. After desktop review, all assessed KIR copy-number and allele calls in the 48 reference samples were concordant with the phased haplotype sequences. Locus-level provisional results and final reviewed concordance are summarized in Table S11.

### 3.2 The workflow showed the lowest allele-level mismatch of four methods

The performance of the Scisco KIR-Allele software in interpreting short-read sequence data was compared with three independently developed KIR genotyping methods: KIR-MAPPER, PING and T1K. Across 311 shared samples, the Scisco workflow showed the lowest overall mismatch rate relative to the four-method consensus (1.62%), followed by KIR-MAPPER (4.54%), ambiguity-adjusted PING (8.85%) and T1K (15.56%) (Table 1). These comparisons used the automated Scisco output before GeMS-KIR12 review, allowing evaluation of computational genotype assignment independently of subsequent manual editing.

Relative performance varied by locus. Scisco mismatch rates were below 2.6% for most genes, with higher rates at KIR3DP1 (7.26%), KIR2DL5 (4.29%) and KIR2DS3 (3.75%). Individual comparator methods showed lower mismatch rates at selected loci, although their overall discordance was greater. KIR-MAPPER showed particularly high mismatch rates at KIR2DL1 (17.51%) and KIR3DP1 (14.52%), while T1K showed substantial discordance at KIR3DL3 (60.33%) and KIR3DL2 (38.06%). PING produced frequent ambiguous allele assignments at several loci, particularly KIR2DL5, KIR2DS2, KIR3DS1 and KIR2DS5. Although ambiguity-aware matching reduced the apparent mismatch rate by counting candidate sets containing the consensus assignment as concordant, these large candidate sets represented reduced resolution of allele assignments.

Together, these findings demonstrate that the automated Scisco KIR-Allele pipeline achieved the highest overall concordance with the four-method consensus among the methods evaluated. This computational comparison complements the independent long-read reference validation described in Section 3.1, supporting the performance of the integrated Scisco system for copy-aware, allele-level KIR genotyping.

### 3.3 Allele richness was greatest at inhibitory and framework loci

After the five-copy filter, retained allele richness ranged from one to 41 alleles per locus within a super-population (Figure S2C; Table S2). Inhibitory and framework loci retained the broadest allele spectra (up to 41 alleles at KIR3DL3, and 24 and 23 at KIR3DL1 and KIR3DL2), whereas activating loci such as KIR2DS1, KIR2DS3 and KIR3DS1 were often represented by only one to three alleles, consistent with previous large studies (13, 14). Because richness depends on sample size, which ranged from 215 (AMR) to 709 (AFR) individuals, differences in richness between super-populations were not interpreted.

### 3.4 Dominant alleles switched between super-populations at four loci

The global allele-frequency landscape revealed marked between-population heterogeneity across the KIR region (Figure 2; Figure S3). The largest descriptive Cramér’s V values were observed for KIR2DS5 (V = 0.350), KIR3DL1 (V = 0.315), KIR2DS1 (V = 0.291) and KIR2DS3 (V = 0.279) (Figure 2A). The selected KIR3DL1, KIR2DL4, KIR3DL2 and KIR3DL3 profiles illustrate shifts in allele-frequency magnitude and changes in the identity of the dominant allele across super-populations (Figure 2B–F). These selected profiles were not the four highest-ranked loci by the 24-population statistic.

This pattern is consistent with the established view that KIR diversity is shaped by both allelic polymorphism and structural heterogeneity, such that different population histories give rise to distinct combinations of frequent alleles and structural backgrounds (5, 6, 15). In 1000 Genomes-derived material, Norman et al. (13) likewise showed that even analysis of a limited subset of KIR genes uncovered substantial previously uncharacterized sequence and structural diversity, supporting the biological plausibility of the differentiated allele landscapes observed here.

This pattern was especially evident for KIR3DL1 and KIR2DL4 (Figure 2B–F). KIR3DL1 showed strong dominant-allele switching across super-populations: KIR3DL1*01501 was the most frequent allele in AFR (0.21), KIR3DL1*01502 predominated in EAS (0.58) and also led in AMR (0.26), KIR3DL1*00401 was most frequent in EUR (0.21), and KIR3DL1*00101 led in SAS (0.36). KIR2DL4 showed a similarly structured pattern, with KIR2DL4*00103 predominating in AFR (0.48), KIR2DL4*00102 leading in both AMR (0.35) and EAS (0.44), and KIR2DL4*00501 most frequent in EUR (0.22) and SAS (0.33).

KIR3DL3 and KIR3DL2 also showed geographically structured, although less uniform, dominant-allele patterns. For KIR3DL3, no single allele predominated across all super-populations: KIR3DL3*00901 led in AFR (0.12), KIR3DL3*00802 in AMR (0.13), KIR3DL3*01001 in EAS (0.25), KIR3DL3*00101 in EUR (0.16), and KIR3DL3*00301 in SAS (0.24). For KIR3DL2, KIR3DL2*00201 predominated in AMR (0.34) and EAS (0.50), whereas KIR3DL2*00103 was most frequent in AFR (0.24) and KIR3DL2*00701 led in EUR (0.20) and SAS (0.32). At these loci, the population signal was not fully captured by the identity of the top allele alone, because multiple moderate-frequency alleles contributed (Figure 2B–E).

Taken together, the principal between-group signal in the allele-centered analysis was locus-specific redistribution among multiple high-frequency alleles, rather than a change in the abundance of a single globally common allele.

### 3.5 Structure resolved at the motif level but not at exact haplotypes

Compatibility matching against the KIR haplotype structure rule set demonstrated substantial ambiguity in exact haplotype assignments (Figure 3; Table S4). Among 2,298 individuals, 197 (8.6%) had a unique exact haplotype-pair solution, 2,009 (87.4%) had multiple compatible solutions, and 92 (4.0%) had no compatible haplotype pair in the structural reference catalog. Resolution improved markedly as exact haplotypes were collapsed into broader structural categories. At the combined centromeric–telomeric motif level, 1,827 individuals (79.5%) were uniquely resolved, while 379 (16.5%) remained ambiguous. At the centromeric motif level, 2,099 individuals (91.3%) were uniquely resolved, with 107 (4.7%) remaining ambiguous. Telomeric motif pairs were uniquely determined for all 2,206 individuals with at least one compatible structural solution. The 92 individuals without compatible solutions remained unresolved at every structural level.

**Figure 3.**
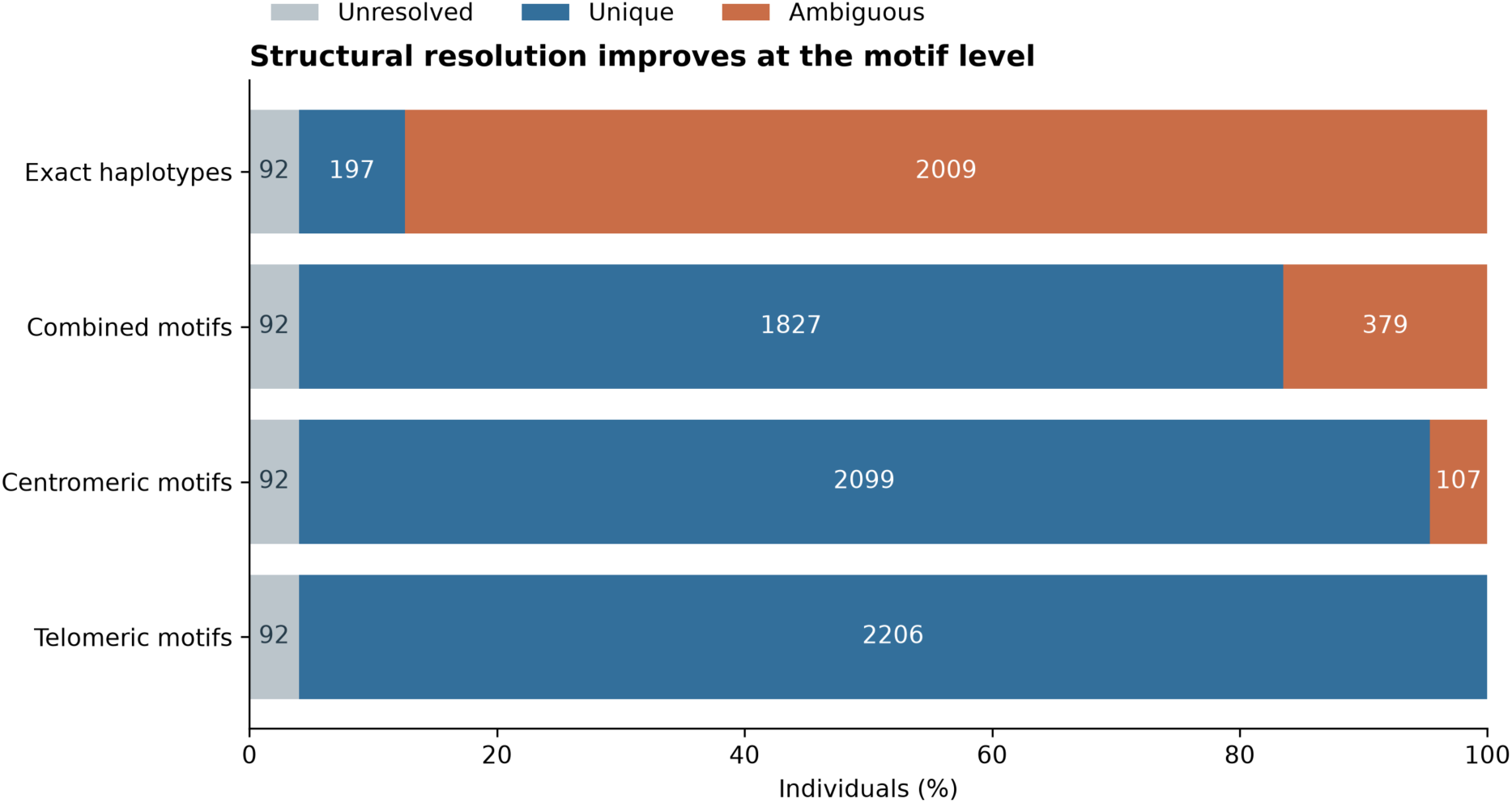
Structural ambiguity and resolution across inference levels. Fractions of individuals (n = 2,298) that are unresolved, uniquely resolved, or ambiguous at the exact-haplotype, collapsed haplotype-combination, centromeric motif, and telomeric motif levels.

The extent of exact-haplotype ambiguity differed among populations, with unique assignments least frequent in EAS (0.72%) and EUR (2.90%) and most frequent in AFR (20.87%) (Table S4). These findings support the use of collapsed haplotype combinations and centromeric and telomeric motifs as the primary structural summaries derived from short-read copy-number profiles, while retaining uncertainty in exact haplotype assignments.

Weighted exact-haplotype architectures, summarized descriptively in Figure S4 and Table S5, showed that the canonical A-haplotype architecture (cA01|tA01) accounted for 56.32% of weighted haplotypes overall, ranging from 41.10% in SAS to 66.57% in EAS. These exact-architecture estimates used the preferred-pair selection procedure described in Section 2.8 and therefore differ from the primary collapsed-motif frequency estimates, which retained all compatible haplotype pairs without preference selection (Section 3.6; Table 2).

**Table 2A.** Weighted centromeric motif frequencies. Ambiguity-aware weighted haplotype frequencies across the overall cohort and five super-populations.

| Centromeric motif | Weighted haplotype count | Overall (%) | AFR (%) | AMR (%) | EAS (%) | EUR (%) | SAS (%) |
| --- | --- | --- | --- | --- | --- | --- | --- |
| cA01 | 3,071.50 | 69.62 | 66.12 | 75.79 | 85.21 | 70.90 | 58.05 |
| cB01 | 691.17 | 15.67 | 19.83 | 9.51 | 4.58 | 13.15 | 23.63 |
| cB02 | 620.83 | 14.07 | 12.32 | 14.07 | 10.13 | 15.85 | 18.27 |
| cB03 | 28.50 | 0.65 | 1.74 | 0.63 | 0.08 | 0.10 | 0.05 |

**Table 2B.** Weighted telomeric motif frequencies. Ambiguity-aware weighted haplotype frequencies across the overall cohort and five super-populations.

| Telomeric motif | Weighted haplotype count | Overall (%) | AFR (%) | AMR (%) | EAS (%) | EUR (%) | SAS (%) |
| --- | --- | --- | --- | --- | --- | --- | --- |
| tA01 | 3,480.00 | 78.88 | 89.72 | 80.90 | 75.98 | 78.75 | 65.60 |
| tB01 | 932.00 | 21.12 | 10.28 | 19.10 | 24.02 | 21.25 | 34.40 |

**Table 2C.** Weighted collapsed haplotype-combination frequencies. Exact haplotype solutions collapsed to centromeric|telomeric structural combinations before ambiguity-aware weighting.

| Collapsed haplotype combination | Weighted haplotype count | Overall (%) | AFR (%) | AMR (%) | EAS (%) | EUR (%) | SAS (%) |
| --- | --- | --- | --- | --- | --- | --- | --- |
| cA01\|tA01 | 2,474.83 | 56.09 | 61.12 | 63.00 | 65.80 | 55.41 | 38.89 |
| cA01\|tB01 | 596.67 | 13.52 | 5.00 | 12.78 | 19.41 | 15.48 | 19.16 |
| cB01\|tA01 | 551.40 | 12.50 | 18.16 | 7.52 | 3.13 | 10.03 | 16.45 |
| cB02\|tA01 | 425.27 | 9.64 | 8.71 | 9.74 | 6.97 | 13.20 | 10.21 |
| cB02\|tB01 | 195.57 | 4.43 | 3.61 | 4.33 | 3.16 | 2.65 | 8.06 |
| cB01\|tB01 | 139.77 | 3.17 | 1.66 | 1.99 | 1.44 | 3.11 | 7.18 |
| cB03\|tA01 | 28.50 | 0.65 | 1.74 | 0.63 | 0.08 | 0.10 | 0.05 |

### 3.6 Motif frequencies differed strongly between super-populations

Among the 2,206 individuals with compatible structural solutions, ambiguity-aware weighting yielded 4,412 haplotype copies for estimating centromeric, telomeric and combined motif frequencies (Figure 4; Table 2).

**Figure 4.**
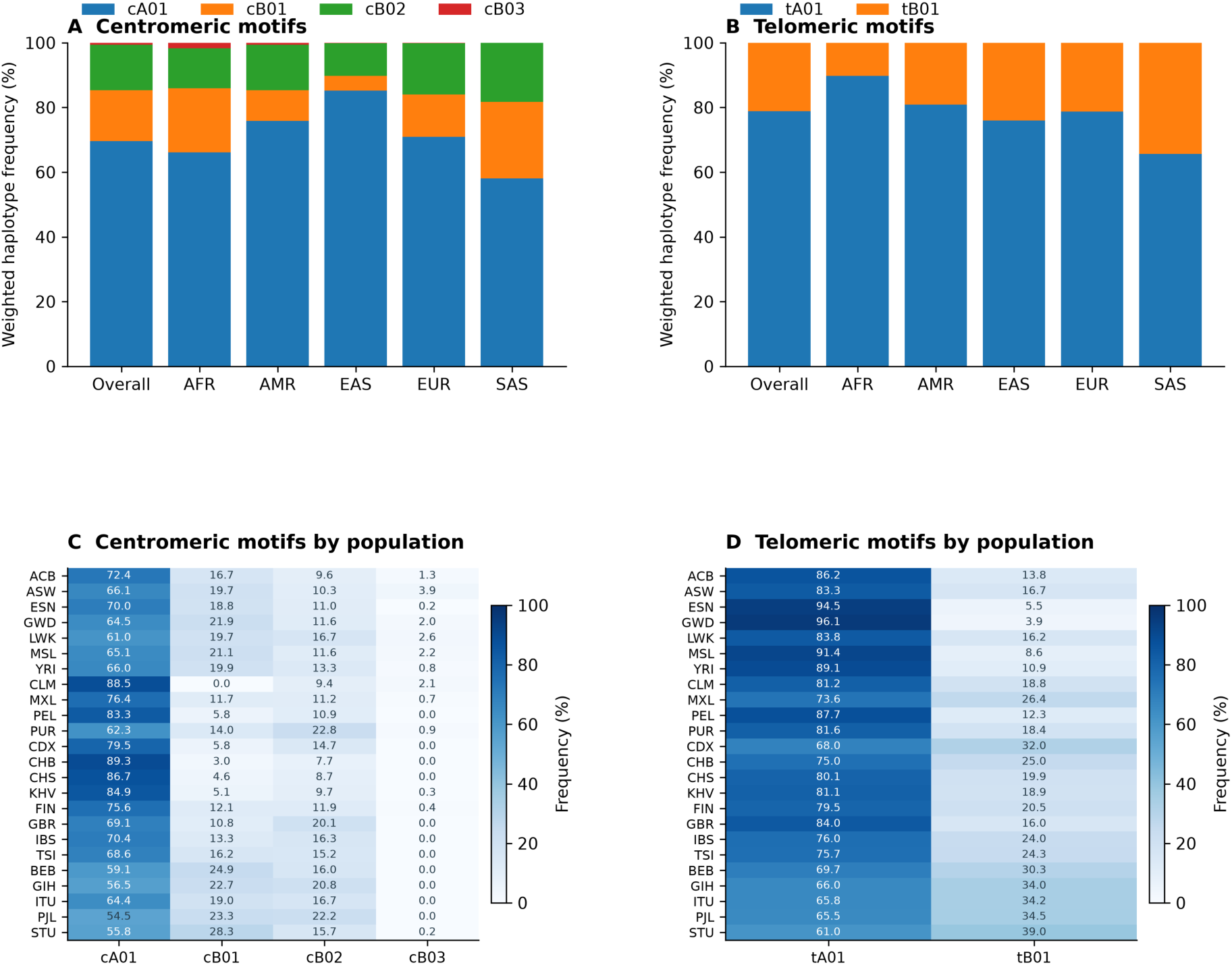
Weighted KIR motif frequencies across the cohort and super-populations. The top row shows stacked weighted frequencies of centromeric motifs and telomeric motifs across the full cohort and the five super-populations. The lower panels show population-level heatmaps for centromeric and telomeric motif frequencies.

After ambiguity-aware weighting of all compatible exact haplotype solutions, the centromeric arm was dominated overall by cA01 (69.62%), followed by cB01 (15.67%), cB02 (14.07%), and a rare cB03 component (0.65%) (Figure 4; Table 2). The telomeric arm was likewise dominated by tA01 (78.88%), whereas tB01 accounted for 21.12% of weighted haplotype copies. Population-level motif frequencies are provided in Table S6.

Super-population differences were pronounced. East Asian samples showed the strongest enrichment of cA01, whereas South Asian samples had the lowest cA01 frequency together with increased representation of both cB01 and cB02. African samples retained the highest tA01 frequency, whereas South Asian samples showed the greatest shift toward tB01. These gradients are directionally consistent with previous reports showing that B-associated KIR structural diversity is unevenly distributed across populations and that the balance between A-like and B-like structural backgrounds differs substantially by ancestry (8, 14).

The weighted profile of collapsed haplotype combinations was likewise readily interpretable (Table 2; Figure S5). Overall, cA01|tA01 was the dominant combination at 56.09%, followed by cA01|tB01 (13.52%), cB01|tA01 (12.50%), cB02|tA01 (9.64%), and cB02|tB01 (4.43%). East Asian samples were most enriched for cA01|tA01 (65.80%), African samples showed the strongest cB01|tA01 component (18.16%), and European samples were relatively enriched for cB02|tA01 (13.20%). South Asian samples showed the broadest redistribution, with cA01|tA01 reduced to 38.89% and several alternative combinations contributing substantial fractions, including cA01|tB01 (19.16%), cB01|tA01 (16.45%), cB02|tA01 (10.21%), cB02|tB01 (8.06%), and cB01|tB01 (7.18%).

### 3.7 Motif backgrounds carried distinct allele compositions

Motif-associated allele composition was evaluated in structurally unambiguous motif-homozygous carriers (Figure 5; Table 3; Table S7). Carrier sets were largest for cA01 (n = 1,082) and tA01 (up to n = 1,388), whereas cB01, cB02 and tB01 were represented by smaller groups. No suitable cB03 homozygous carrier set was available. These analyses describe allele composition within the selected motif-homozygous groups rather than allele frequencies across all carriers of each motif.

**Figure 5.**
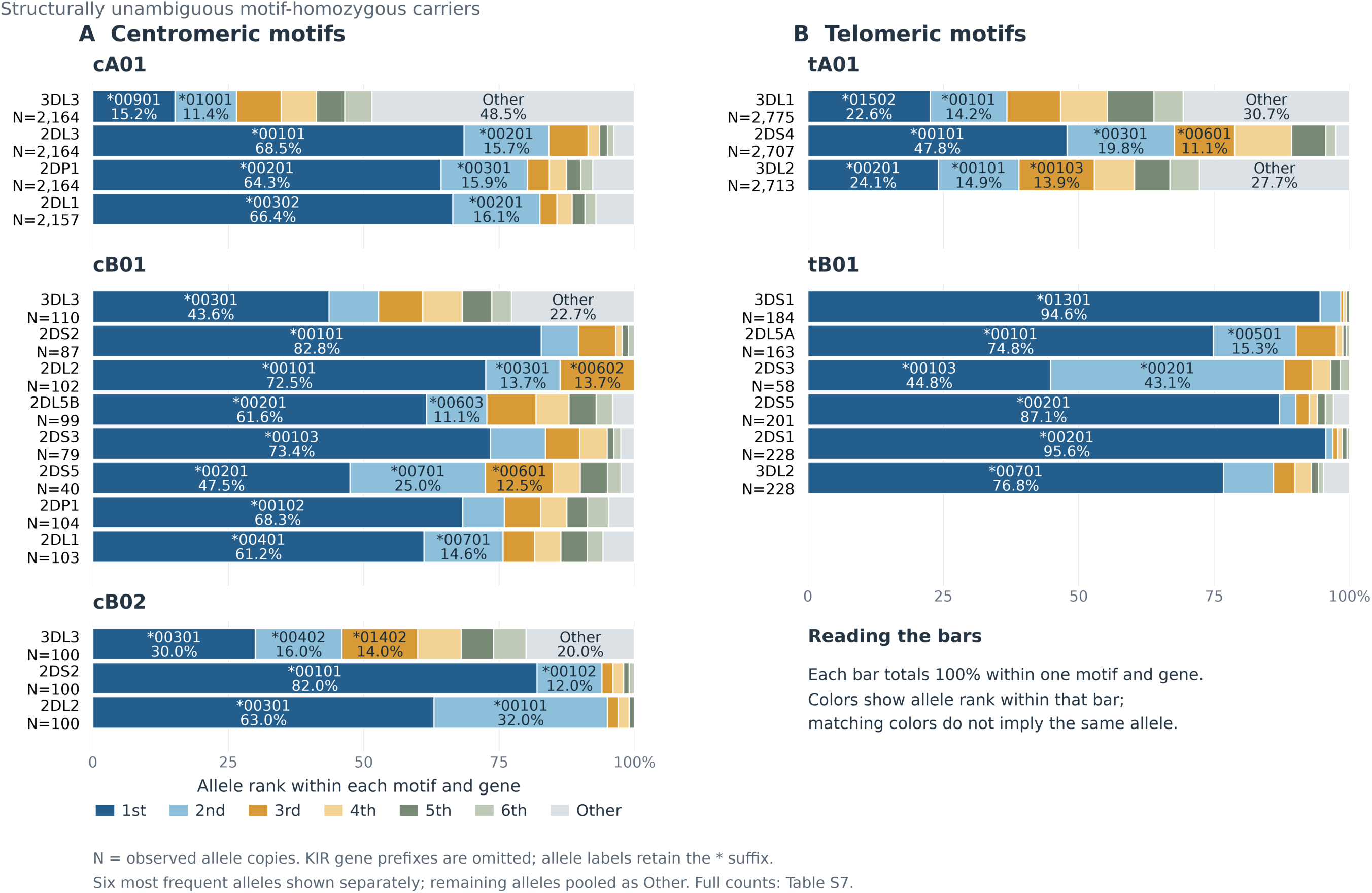
Centromeric and telomeric motif-associated KIR allele composition from structurally unambiguous motif-homozygous carriers. Horizontal stacked bars show allele frequencies among structurally unambiguous motif-homozygous carriers. (A) Centromeric motifs cA01, cB01 and cB02. (B) Telomeric motifs tA01 and tB01. The six most frequent alleles within each motif and gene are shown separately; remaining alleles are pooled as Other. Colors denote frequency rank within each bar, rather than shared allele identity. Labels show gene-specific allele suffixes and frequencies; N denotes observed allele copies. Complete distributions and gene-specific carrier counts are provided in Table S7.

**Table 3A.** Leading centromeric motif-associated alleles. For each motif-specific gene, the two most frequent alleles among structurally unambiguous motif-homozygous carriers are shown; remaining alleles are grouped as Other. Full distributions are provided in Table S7. Support columns are gene-specific because only carriers with the corresponding gene contribute copies.

| Motif | Gene | Leading allele | Leading allele frequency (%) | Second allele | Second allele frequency (%) | Other alleles (%) | Total allele copies | Contributing individuals (gene-specific) |
| --- | --- | --- | --- | --- | --- | --- | --- | --- |
| cA01 | 2DL1 | 2DL1*00302 | 66.43 | 2DL1*00201 | 16.09 | 17.48 | 2,157 | 1,082 |
| cA01 | 2DL3 | 2DL3*00101 | 68.53 | 2DL3*00201 | 15.67 | 15.80 | 2,164 | 1,082 |
| cA01 | 2DP1 | 2DP1*00201 | 64.28 | 2DP1*00301 | 15.94 | 19.78 | 2,164 | 1,082 |
| cA01 | 3DL3 | 3DL3*00901 | 15.16 | 3DL3*01001 | 11.37 | 73.48 | 2,164 | 1,082 |
| cB01 | 2DL1 | 2DL1*00401 | 61.17 | 2DL1*00701 | 14.56 | 24.27 | 103 | 55 |
| cB01 | 2DL2 | 2DL2*00101 | 72.55 | 2DL2*00301 | 13.73 | 13.73 | 102 | 55 |
| cB01 | 2DL5B | 2DL5B*00201 | 61.62 | 2DL5B*00603 | 11.11 | 27.27 | 99 | 54 |
| cB01 | 2DP1 | 2DP1*00102 | 68.27 | 2DP1*010 | 7.69 | 24.04 | 104 | 54 |
| cB01 | 2DS2 | 2DS2*00101 | 82.76 | 2DS2*00102 | 6.90 | 10.34 | 87 | 50 |
| cB01 | 2DS3 | 2DS3*00103 | 73.42 | 2DS3*00201 | 10.13 | 16.46 | 79 | 48 |
| cB01 | 2DS5 | 2DS5*00201 | 47.50 | 2DS5*00701 | 25.00 | 27.50 | 40 | 29 |
| cB01 | 3DL3 | 3DL3*00301 | 43.64 | 3DL3*02801 | 9.09 | 47.27 | 110 | 55 |
| cB02 | 2DL2 | 2DL2*00301 | 63.00 | 2DL2*00101 | 32.00 | 5.00 | 100 | 50 |
| cB02 | 2DS2 | 2DS2*00101 | 82.00 | 2DS2*00102 | 12.00 | 6.00 | 100 | 50 |
| cB02 | 3DL3 | 3DL3*00301 | 30.00 | 3DL3*00402 | 16.00 | 54.00 | 100 | 50 |

**Table 3B.**
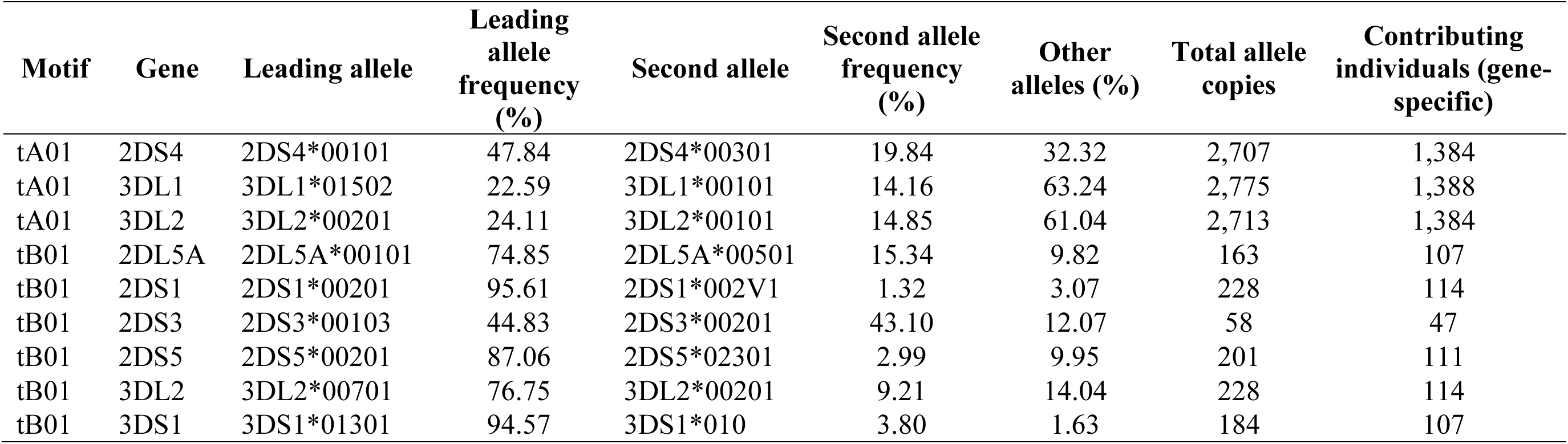
Leading telomeric motif-associated alleles. For each motif-specific gene, the two most frequent alleles among structurally unambiguous motif-homozygous carriers are shown; remaining alleles are grouped as Other. Full distributions are provided in Table S7. Support columns are gene-specific because only carriers with the corresponding gene contribute copies.

| Motif | Gene | Leading allele | Leading allele frequency (%) | Second allele | Second allele frequency (%) | Other alleles (%) | Total allele copies | Contributing individuals (gene-specific) |
| --- | --- | --- | --- | --- | --- | --- | --- | --- |
| tA01 | 2DS4 | 2DS4*00101 | 47.84 | 2DS4*00301 | 19.84 | 32.32 | 2,707 | 1,384 |
| tA01 | 3DL1 | 3DL1*01502 | 22.59 | 3DL1*00101 | 14.16 | 63.24 | 2,775 | 1,388 |
| tA01 | 3DL2 | 3DL2*00201 | 24.11 | 3DL2*00101 | 14.85 | 61.04 | 2,713 | 1,384 |
| tB01 | 2DL5A | 2DL5A*00101 | 74.85 | 2DL5A*00501 | 15.34 | 9.82 | 163 | 107 |
| tB01 | 2DS1 | 2DS1*00201 | 95.61 | 2DS1*002V1 | 1.32 | 3.07 | 228 | 114 |
| tB01 | 2DS3 | 2DS3*00103 | 44.83 | 2DS3*00201 | 43.10 | 12.07 | 58 | 47 |
| tB01 | 2DS5 | 2DS5*00201 | 87.06 | 2DS5*02301 | 2.99 | 9.95 | 201 | 111 |
| tB01 | 3DL2 | 3DL2*00701 | 76.75 | 3DL2*00201 | 9.21 | 14.04 | 228 | 114 |
| tB01 | 3DS1 | 3DS1*01301 | 94.57 | 3DS1*010 | 3.80 | 1.63 | 184 | 107 |

Centromeric motifs showed distinct patterns of allele representation. The cA01 background was dominated by KIR2DL1*00302 (66.43%), KIR2DL3*00101 (68.53%) and KIR2DP1*00201 (64.28%), whereas KIR3DL3 was substantially more diverse, with its leading allele, KIR3DL3*00901, accounting for only 15.16% of copies. In contrast, cB01 showed strong enrichment of KIR2DS2*00101 (82.76%) and KIR2DL2*00101 (72.55%), together with characteristic alleles at KIR2DL1, KIR2DL5B and the KIR2DS3/2DS5 branch positions. The cB02 background was similarly dominated by KIR2DS2*00101 (82.00%) but differed from cB01 in its leading KIR2DL2 allele, KIR2DL2*00301 (63.00%). Thus, centromeric motif backgrounds differed not only in gene content but also in the alleles associated with shared genes.

Telomeric motifs exhibited a particularly clear contrast in allele concentration. The tA01 background retained broad allelic diversity at KIR3DL1 and KIR3DL2, whose leading alleles accounted for only 22.59% and 24.11% of copies, respectively. By comparison, tB01 was dominated by single alleles at several loci, including KIR3DS1*01301 *(94.57%), KIR2DS1*00201* (95.61%), KIR2DS5*00201 *(87.06%) and KIR3DL2*00701* (76.75%). KIR2DS3 was an exception, with *00103 (44.83%) and *00201 (43.10%) occurring at similar frequencies. The same general distinction was evident across super-populations (Figure S6; Table S8), where tB01-associated KIR3DS1 and KIR2DS1 remained strongly concentrated in single alleles, while tA01 genes retained broader allele distributions.

Together, these findings show that structural motif backgrounds are associated with characteristic, but gene-dependent, patterns of allelic diversity. The distributions provide a link between KIR gene-content structure and allele composition without requiring assignment of individual alleles to physically phased chromosomes.

### 3.8 Allele resolution distinguished KIR3DL1 and HLA Bw4 configurations

Pairing KIR with HLA showed that the same gene-level receptor–ligand call could represent different inherited receptor configurations (Figure 6; Table 4). Among 1,401 individuals carrying both KIR3DL1 and HLA-B Bw4, 1,321 carried at least one allele annotated for ordinary high or low surface expression, 57 (4.1%) lacked such an allele, and 23 had uncertain annotation. The 57 included 44 individuals with one KIR3DL1*004 copy, eight with *004/*004, two with *004/*019, two with one *019 copy, and one with a *16801 call mapping to a null reference. Thus, both single-copy and two-copy gene-positive calls could lack an ordinarily surface-expressed allele. This distinction concerns surface annotation; intracellular retention does not establish absence of all biological function (43).

**Figure 6.**
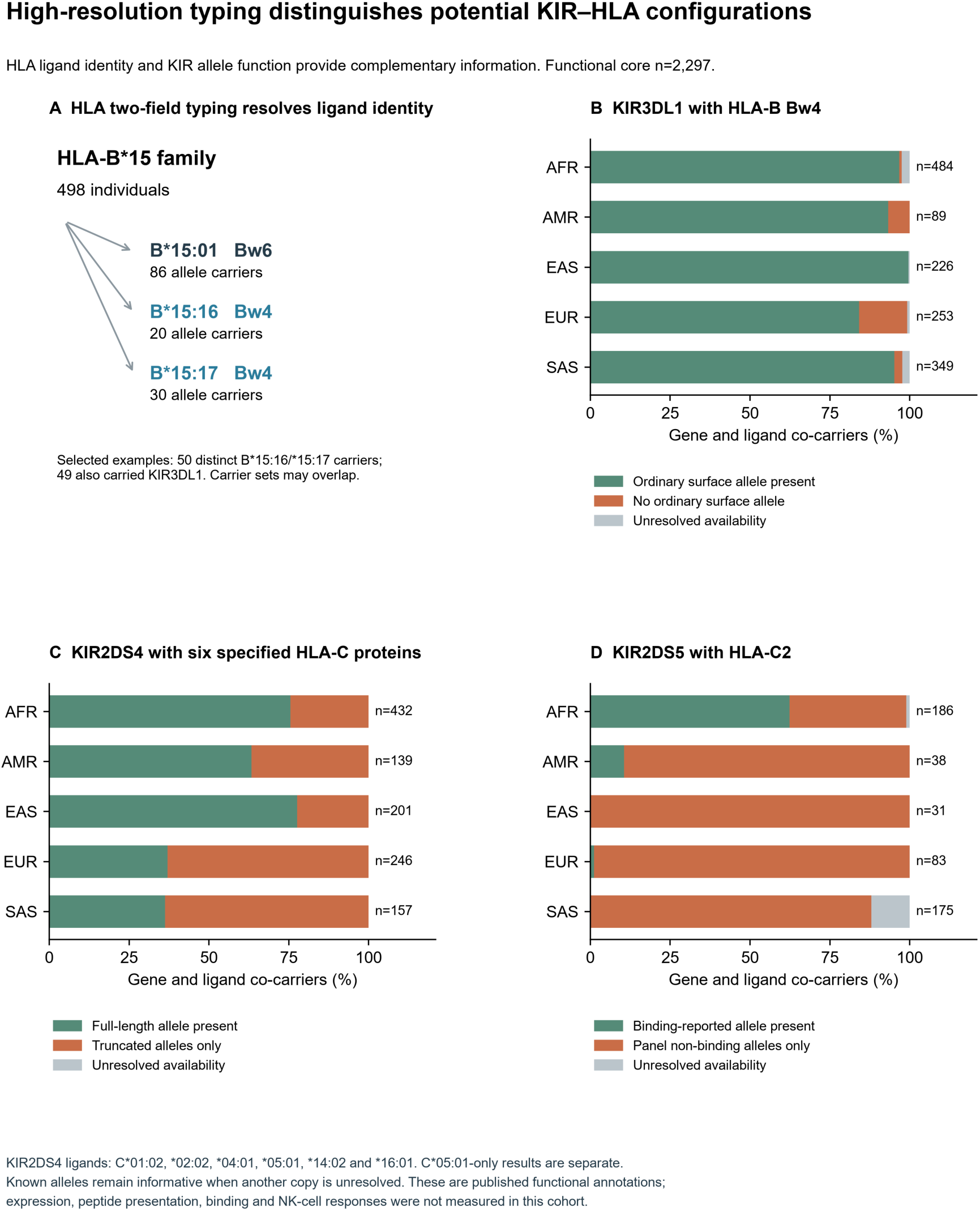
High-resolution typing distinguishes potential KIR and HLA configurations. (A) Selected two-field HLA-B*15 examples distinguish Bw6-bearing B*15:01 from Bw4-bearing B*15:16 and B*15:17 within one first-field family. Counts are allele carriers and can overlap; 50 distinct individuals carried B*15:16 or B*15:17, including 49 KIR3DL1 carriers. (B) Ordinary surface-expression annotation among 1,401 KIR3DL1/HLA-B Bw4 co-carriers. (C) Full-length receptor availability among 1,175 KIR2DS4 co-carriers of C*01:02, C*02:02, C*04:01, C*05:01, C*14:02 or C*16:01. The C*05:01-only analysis is retained separately in Table S9C. (D) Published C2-binding annotation among 513 KIR2DS5/C2 co-carriers. Each bar uses its displayed gene/ligand denominator; an available allele remains informative when another copy is unresolved. These inherited labels are not cohort measurements of expression, binding or NK-cell activity. KIR3DL1 surface retention does not exclude other function; KIR2DS4 recognition is peptide dependent; panel non-binding does not imply universal inactivity.

**Table 4.** Biological information gained from paired high-resolution KIR and HLA typing. Eligible people are gene/ligand co-carriers except for the HLA-B*15 family and whole-core repertoire rows. KIR2DS4 covers six specified HLA-C proteins; its C*05:01-only subset is retained separately. Original observed alleles remain informative beside provisional or missing partners. Repertoire ranges are lower/upper annotation scenarios, not confidence intervals. These inherited labels do not measure receptor expression, peptide presentation, binding or NK-cell activity. Table S9 provides population, allele, copy, motif and inferential support.

| Receptor and ligand or repertoire | Eligible people | Information resolved by high-resolution typing | Allele and copy examples | Population or ligand context |
| --- | --- | --- | --- | --- |
| HLA-B*15 family | 498 | Two-field alleles distinguish Bw4 from Bw6 | B*15:01: Bw6; B*15:16 and B*15:17: Bw4 | 50 distinct B*15:16/*15:17 carriers; 49 also carried KIR3DL1 |
| KIR3DL1 / HLA-B Bw4 | 1,401 | 1,321 with ordinary surface allele; 57 without; 23 unresolved | Single *004, *004/*004 and a surface-expressed partner differ | No ordinary surface allele: EUR 38/253 (15.0%); AFR 4/484 (0.8%) |
| KIR2DL1 / HLA-C2 | 1,384 | 906 R245-only; 111 C245-only; 347 mixed; 20 unresolved | *003/*003 (R/R), *003/*004 (R/C), *004/*004 (C/C) | C245-only: SAS 46/336 (13.7%); EAS 1/119 (0.8%); linked centromeric contexts |
| KIR2DS4 / six HLA-C proteins | 1,175 | 718 with a full-length allele; 457 truncated-only | Full-length *001 versus truncated *003/*006; known alleles retained with unresolved partners | Truncated-only: EUR 155/246 (63.0%); AFR 106/432 (24.5%). C*05:01 subset: 57/112 overall |
| KIR2DS5 / HLA-C2 | 513 | 121 with binding-reported allele; 369 panel non-binding only; 23 unresolved | Binding-reported *003-*008 versus *002/*009/*010/*011 | Binding-reported: AFR 116/186 (62.4%); EUR 1/83 (1.2%) |
| Distinct potential allotype pairs | 2,297 | Mean 6.10-6.20; 144 people with annotation uncertainty | Collapse synonymous alleles and identical repeated copies; retain different KIR and HLA proteins | AFR 6.26-6.40; SAS 6.29-6.45; composition differs |

HLA resolution also changed ligand interpretation (Figure 6A). The B*15 family included Bw6-bearing B*15:01, carried by 86 individuals, and Bw4-bearing B*15:16 and B*15:17, carried by 20 and 30 individuals, respectively. Fifty distinct individuals carried B*15:16 or B*15:17, and 49 also carried KIR3DL1. These two-field distinctions would be lost in a first-field B*15 category. Full numeric labels were retained for traceability, but an additional functional consequence of third-field differences was not assumed. Likewise, KIR3DL1 expression classes were not converted into a universal ranking of HLA binding strength (57).

The frequency of this category differed across populations. It accounted for 38/253 KIR3DL1/Bw4 co-carriers in EUR (15.0%), compared with 4/484 in AFR (0.8%), 6/89 in AMR (6.7%), 0/226 in EAS and 9/349 in SAS (2.6%) (Figure 6B). Within EUR, the corresponding counts were 12/65 in IBS, 8/50 in FIN, 10/75 in TSI and 8/63 in GBR (Figure S7A; Table S9B–C). These proportions retained uncertain calls as a separate category and were not interpreted as measured differences in NK-cell response.

Copy and motif information made the same result more specific (Figure 7A). Among EUR gene/Bw4 co-carriers with a unique telomeric motif pair, 33/90 individuals with tA01/tB01 lacked an ordinary surface allele (36.7%), compared with 5/157 with tA01/tA01 (3.2%). The first group comprised single-copy *004 (n = 30), *019 (n = 2) and the resolved null *16801 call (n = 1). The second comprised *004/*004 (n = 4) and *004/*019 (n = 1). Gene presence, copy count and motif context therefore provided complementary information rather than interchangeable descriptions.

**Figure 7.**
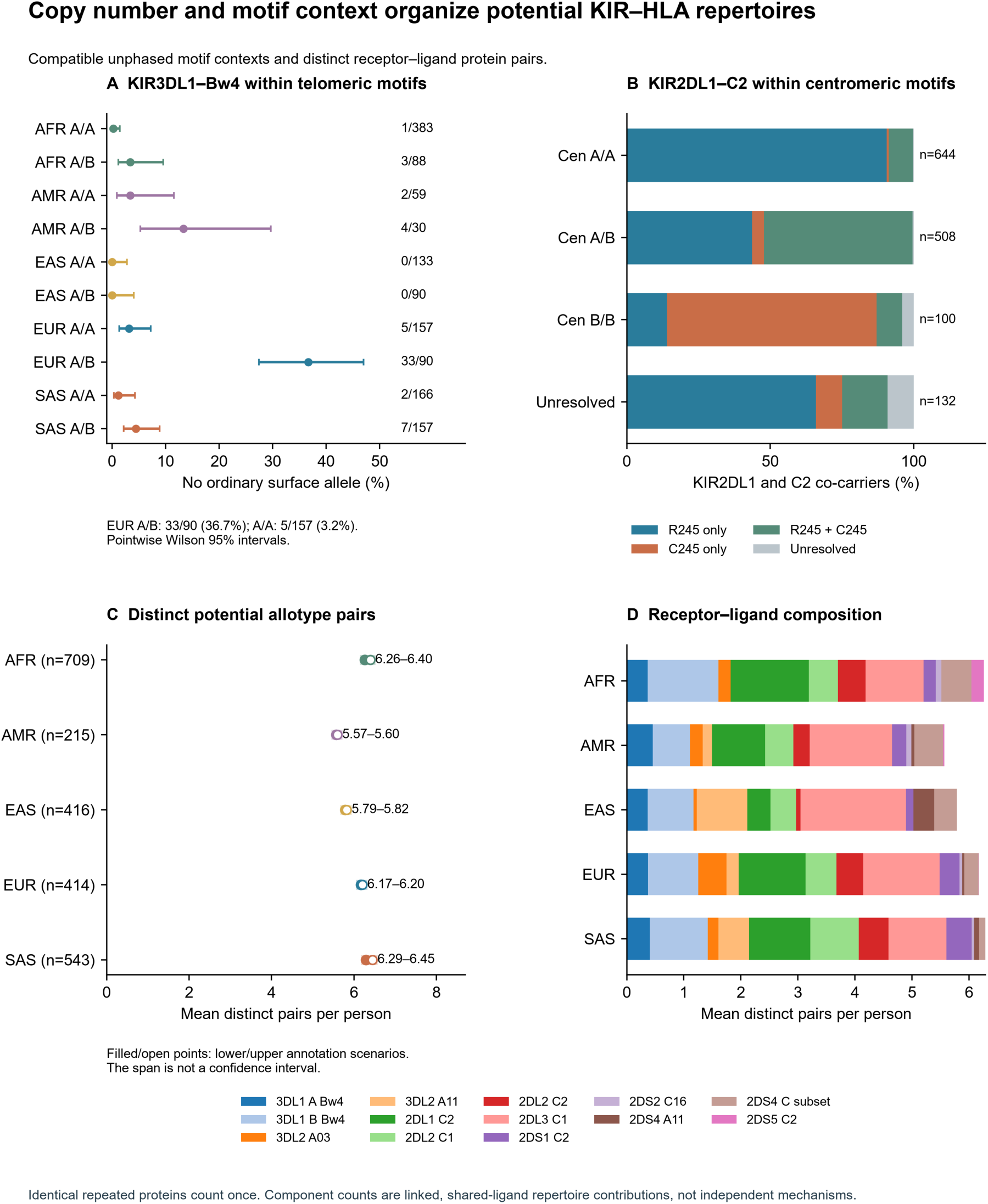
Copy number and motif context organize potential KIR and HLA repertoires. (A) No ordinarily surface-expressed KIR3DL1 allele among HLA-B Bw4 co-carriers within unique tA01/tA01 and tA01/tB01 contexts. Counts and pointwise Wilson 95% intervals are shown. EUR tA01/tB01 includes 30 single-copy *004, two *019 and one resolved null *16801 call. (B) KIR2DL1 residue-245 profiles among C2 co-carriers within centromeric A/A, A/B and B/B contexts; unresolved contexts remain explicit. Motifs are compatible unphased contexts and do not assign an allele to a homolog. (C) Mean distinct potential KIR-protein/HLA-protein pairs per individual under the declared 2019 specificity rules. Filled/open points show lower/upper annotation scenarios, not confidence intervals. Identical repeated KIR or HLA proteins count once. (D) Lower-scenario contributions by receptor/ligand rule. Shared ligands and linked genes contribute repertoire components, not independent mechanistic findings. Functional labels follow published evidence and do not measure NK-cell responses.

### 3.9 HLA C combinations resolved receptor sequence form and binding annotations

Among 1,384 KIR2DL1/C2 co-carriers, allele typing distinguished 906 R245-only profiles, 111 C245-only profiles, 347 mixed profiles and 20 unresolved profiles (Figure 7B; Table 4). Exact examples were *003/*003 (R/R; n = 328), *003/*004 (R/C; n = 161) and *004/*004 (C/C; n = 14). These genotypes share a positive KIR2DL1/C2 gene-level call, and the examples each contain two receptor copies, but differ at a residue with published effects on inhibitory signaling and surface stability (44). C245-only profiles were more frequent among the SAS co-carriers (46/336, 13.7%) and AFR co-carriers (55/537, 10.2%) than among EAS co-carriers (1/119, 0.8%). Mixed profiles accounted for 30.2% in AFR, 28.3% in SAS and 4.2% in EAS. Their distribution across centromeric motif contexts linked these sequence classes to the structural backgrounds described above (Figure 7B), without assigning alleles to individual chromosomes.

KIR2DS4 provided a different example, involving receptor form. Of 112 KIR2DS4/HLA-C*05:01 gene co-carriers, only 55 carried a predicted full-length receptor allele: 20 had full-length-only genotypes and 35 had mixed full-length and truncated genotypes. The remaining 57 carried only truncated alleles (Table S9C; Figure S7C). The latter category included alleles such as *003 and *006, whereas *001 represented a full-length form (45). Truncated-only genotypes accounted for 36/59 co-carriers in EUR (61.0%), 13/21 in AMR (61.9%) and 5/23 in AFR (21.7%); within GBR the count was 14/20 (70.0%). The EAS and SAS comparisons involved only three and six co-carriers, respectively. These counts identify the potential for a full-length receptor, not activation by HLA-C*05:01, which depends on the presented peptide (58). The broader analysis of six specified HLA-C proteins included 1,175 KIR2DS4 co-carriers, of whom 718 had an observed full-length allele and 457 (38.9%) carried only truncated alleles (Figure 6C). Known full-length alleles remained informative when another copy was unresolved. Truncated-only genotypes occurred in 155/246 EUR (63.0%), 100/157 SAS (63.7%) and 106/432 AFR (24.5%) co-carriers.

KIR2DS5 showed the clearest difference in published binding annotation (Figure 6D). Among 513 KIR2DS5/C2 gene co-carriers, 121 carried an allotype with reported C2 binding, 369 carried only allotypes for which no binding was detected in the published assay panel, and 23 remained uncertain. Binding-annotated alleles occurred in 116/186 AFR co-carriers (62.4%), including 34/48 in LWK and 15/20 in GWD, but in only 1/83 EUR co-carriers (1.2%) and none of the 31 EAS co-carriers. All 31 EAS co-carriers carried *002; the single EUR co-carrier with binding annotation carried *005 and was in TSI. Within complete AFR paired genotypes, *007 was carried by 39/183 individuals. These assignments follow experimental differences among KIR2DS5 allotypes (46). The absence of a binding-annotated allele was not equated with universal inactivity, and the 21 uncertain SAS co-carriers were retained rather than assigned to the panel non-binding group.

### 3.10 Allele and motif variation produced different potential repertoires

The individual examples were combined by counting distinct potential KIR-allotype/HLA-allotype pairs under the specificity rules used by Nemat-Gorgani et al. (48). Identical repeated proteins counted once, so the repertoire reflected protein diversity rather than a product of gene copies. Across 2,297 individuals, the mean was 6.10 pairs under the lower annotation scenario and 6.20 under the upper scenario; 144 people had a nonzero annotation-uncertainty span (Figure 7C; Table S9AE–AH). The super-population means were AFR 6.26–6.40, AMR 5.57–5.60, EAS 5.79–5.82, EUR 6.17–6.20 and SAS 6.29–6.45. The ranges describe annotation scenarios, not confidence intervals. The 24 population summaries are supplied in Table S9AE; lower-scenario means ranged from 4.94 in CLM (n = 16) to 6.74 in BEB (n = 104).

Similar total repertoire sizes had different receptor–ligand compositions (Figure 7D; Table S9AF). KIR2DS5/C2 pairs were concentrated in AFR, whereas HLA-A*11 combinations made a larger contribution in EAS and SAS. KIR2DS4/HLA-C combinations also contributed more in AFR than in SAS. Thus, comparable repertoire counts did not imply interchangeable receptor composition. Copy counts and centromeric/telomeric motifs described the linked contexts carrying these proteins, while retaining neighboring receptors as distinct molecular pair types. These genotype-based repertoires do not establish coexpression on individual cells, peptide presentation or measured NK-cell responses.

### 3.11 Between-population receptor and ligand frequencies after genomic calibration

Between-population frequency covariation addressed a separate question. Across 24 populations, C2-binding-annotated KIR2DS5 copies correlated with HLA-C2 before genomic calibration (full-core Spearman rho = 0.693; genome-matched rho = 0.672). High-expression-annotated KIR3DL1/Bw4-80I and KIR2DL1 R245/C2 had raw full-core rho values of 0.353 and −0.551, respectively. In 2,164 public-genome-matched individuals, covariance adjustment and a chromosome-separated, frequency-matched genomic reference provided adequate support for all 23 reserved comparisons. None was significant after genomic calibration and Holm correction (all adjusted p = 1; Figure S8B; Table S9AM–AR). For KIR2DS5/C2, the adjusted frequency r was 0.147 and the unadjusted genomic empirical p was 0.529. The distinct raw and adjusted statistics are reported separately.

### 3.12 Population-stratified tests did not establish consistent KIR and HLA co-carriage

The information gained from allele annotation was distinct from preferential genetic co-carriage. The four known gene-level receptor–ligand comparisons had population-stratified odds ratios from 0.84 to 1.26, and all four Holm-adjusted p values were 1 (Figure S8A; Table S9M). The smallest adjusted p in the six-test targeted family was 0.233 for the KIR2DL1 residue-245 profile/C2 comparison (raw p = 0.039). Exploratory allele-resolution screens yielded no Benjamini–Hochberg (55) or Benjamini– Yekutieli (56) discoveries in their original families. None of the 72 regional tests passed Holm (54) correction after preserving linked KIR profiles and conditioning the resolution refinements on the regional parent genotype; the smallest adjusted p was 0.307 (Figure S7E; Table S9O–Q).

An earlier KIR2DL2/C1 analysis did identify super-population heterogeneity within its original eight-test family (Holm p = 0.0256), with opposite directions in EUR and SAS. The result was sensitive to correction scope (p = 0.0511 across the combined 16 primary and pedigree-sensitivity tests). In the later joint KIR2DL2/2DS2 analysis, the heterogeneity p was 0.0036 before correction and 0.661 after adjustment across 367 supporting tests (Figure S7F). This was a different, linked-marker contrast on overlapping samples, not independent replication. These tests were stratified by population and therefore condition out between-population frequency covariation, which is where published evidence for KIR–HLA coevolution resides (48, 51); they constrain within-population assortment of receptor and ligand only. The between-population question is addressed separately in Section 3.11. The data therefore support retaining allele-based biological annotation and genomic context, while leaving receptor-specific within-population genetic enrichment unresolved.

## 4. Discussion

This study evaluated an integrated system for high-resolution KIR genotyping that combines a targeted short-read sequencing assay, centralized copy-aware allele analysis, and interactive review of genotype assignments. Applied to 2,298 individuals from 24 populations of the 1000 Genomes Project, the system provided allele-level genotypes, gene copy-number assignments, and predictions of KIR structural backgrounds across all 17 loci. Beyond evaluating genotyping performance, the resulting dataset permitted population-scale characterization of KIR allele diversity, relationships between alleles and haplotype motifs, and inherited KIR–HLA receptor–ligand configurations that cannot be distinguished by conventional gene-content typing.

The laboratory workflow also accommodates different sample volumes and sequencing platforms. High-throughput batches of 768 samples were prepared for NextSeq sequencing within three days, whereas smaller batches of 96–192 samples were prepared for MiSeq or MiSeq i100 sequencing within two days. This flexibility allows KIR genotyping to be adapted to both large population studies and smaller laboratory workloads using existing short-read sequencing infrastructure.

Independent validation against complete, phased PacBio KIR sequences demonstrated high concordance between the short-read system and reference genotypes. Automated assignments were concordant for copy number in 98.6% of gene–sample comparisons and for both copy number and allele identity in 92.8%. Following GeMS-KIR12 review, all 768 evaluated assignments were concordant with the reference sequences. Because the corresponding long-read sequences were excluded from the allele reference set and were unavailable during review, this comparison independently assessed the accuracy of the integrated assay-and-analysis workflow. Most provisional discrepancies involved allele assignments rather than copy number, particularly at KIR3DP1, KIR2DP1 and KIR2DL1, illustrating the value of expert review in resolving difficult genotypes.

Comparison with three independently developed computational approaches provided complementary evidence for the performance of the automated analysis. Across 311 shared samples, the Scisco KIR-Allele pipeline showed the lowest overall mismatch with the four-method consensus, although performance varied among genes (34–36). Because these comparisons used provisional automated calls, the results were independent of subsequent desktop editing. The lower overall discordance supports the utility of copy-number-constrained allele interpretation, particularly in a gene family characterized by extensive paralogy and structural variation. The relative contributions of individual computational features to performance were not evaluated separately.

The closest technological precedent is the exon-targeted, copy-aware KIR typing approach described by Wagner et al. (28), which demonstrated allele-level genotyping at registry scale. The present system follows similar principles but integrates targeted amplification, cloud-based analysis, copy-number reconciliation, structural interpretation and interactive genotype review into a unified workflow. Unlike long-read sequencing, which can directly determine complete KIR haplotype sequences, the short-read approach requires structural inference from gene content and copy-number information. Its principal practical advantage is compatibility with established short-read laboratory infrastructure and standardized analytical procedures.

The population analyses demonstrate both the utility and the limitations of KIR structural inference from short-read data. Exact haplotype pairs could be assigned uniquely to fewer than 10% of individuals, whereas resolution improved markedly after collapsing compatible structures into centromeric and telomeric motifs. These findings support the modular interpretation of KIR haplotypes established in earlier studies (5–8). The resulting motif distributions varied among populations, as did the allele compositions associated with individual structural backgrounds. Notably, some genes on tB01 backgrounds were dominated by a single allele, whereas genes on tA01 backgrounds frequently exhibited broader allelic diversity. Differences in leading alleles at genes shared between structural backgrounds further demonstrate that gene content alone does not adequately describe KIR genetic variation. Although these associations were derived from structurally unambiguous motif-homozygous carriers rather than direct chromosome-level phasing, they establish a useful relationship between haplotype structure and allele composition.

The paired KIR and HLA analyses illustrate the biological significance of this additional resolution. Individuals with identical gene-level receptor–ligand classifications could carry KIR alleles encoding proteins with different documented properties. Among KIR3DL1/HLA-B Bw4 co-carriers, allele typing distinguished conventionally surface-expressed allotypes from genotypes lacking an allele annotated for ordinary surface expression. KIR2DL1 alleles distinguished residue-245 variants associated with differences in inhibitory signaling, while KIR2DS4 typing separated full-length from truncated receptor forms. Similarly, KIR2DS5 allele assignments distinguished allotypes with reported HLA-C2 binding from those lacking detectable binding in published experimental panels (42–46). The frequencies of these configurations differed among populations, illustrating how gene-content typing can conceal substantial inherited differences in potential receptor function.

Combining these annotations with high-resolution HLA genotypes further permitted estimation of individual potential receptor–ligand repertoires using previously established specificity rules (48). Although mean repertoire sizes were broadly similar among several population groups, their constituent receptor–ligand combinations differed. Thus, repertoire composition may provide information not captured by either gene-content typing or the total number of potentially interacting pairs. These sequence-derived predictions do not establish receptor expression, peptide-dependent binding, NK-cell education or functional responses, which require direct experimental assessment.

The availability of paired KIR and HLA genotypes also permitted examination of potential genetic associations between receptor and ligand variation. Some receptor–ligand frequencies showed substantial correlations across populations, but none remained significant after calibration against genome-wide population covariance and correction for multiple testing. Population-stratified analyses likewise did not establish consistent enrichment of specific receptor–ligand combinations within populations. These approaches address different aspects of genetic association: between-population frequency covariation may reflect population history or selection, whereas within-population tests evaluate genetic co-carriage after accounting for population structure. The present findings therefore do not establish preferential genetic association beyond the modeled population background, nor do they exclude receptor–ligand coevolution reported in broader population comparisons (48, 51). Importantly, the absence of statistically robust genetic association does not diminish the established functional differences among KIR allotypes or the value of resolving their inherited combinations with HLA.

An important practical feature of the system is that the KIR and HLA assays use the same basic two-stage multiplex PCR workflow, sample processing procedures and Illumina sequencing infrastructure. The principal difference lies in sequence interpretation. Whereas HLA typing generally begins with established gene identities and diploid copy number, KIR analysis must first establish gene presence and copy number, resolve highly homologous paralogs, and infer compatible haplotype structures. Centralized cloud-based analysis allows these more complex computational procedures to be performed without requiring dedicated local bioinformatics infrastructure. This compatibility with existing HLA laboratory operations provides a practical route for incorporating allele-level KIR typing into research and potentially clinical workflows.

Several limitations should be recognized. The four-method comparison evaluated agreement with a consensus that included the present workflow and therefore does not provide an independent measure of absolute accuracy; the separate PacBio reference comparison addresses this limitation for the evaluated samples. Short-read sequence data cannot generally establish complete chromosome-level KIR haplotypes, and 92 individuals (4.0%) had gene-copy profiles incompatible with the structural reference catalog. This limitation concerns haplotype inference rather than necessarily indicating incorrect allele assignments, which use a separate allele reference set. Because the assay targets exonic sequences, intronic and regulatory differences may remain unresolved. In addition, the functional annotations and receptor–ligand repertoires were inferred from published evidence rather than measured directly, and some genetic association analyses were limited by sample size and the available population structure. Genomic calibration was restricted to the 2,164 core individuals with matched public genotypes; the other 133 individuals remained in the full-core analyses. This availability restriction changed the population denominators and may limit the generalizability of the calibrated results to the full typed cohort (Table S9AO).

This study establishes an integrated, scalable approach to high-resolution KIR genotyping that combines accurate copy-aware allele assignment with ambiguity-aware structural interpretation. By using laboratory procedures and sequencing infrastructure shared with established HLA typing systems, the technology provides a practical means of incorporating allele-resolved KIR information into large-scale immunogenetic studies and future investigations of KIR–HLA contributions to human disease.

## Supporting information

all supplementary files

## Data availability statement

The data and code supporting the conclusions of this article are provided as supplementary material. Analysis code, dependencies and instructions are provided in Supplementary Code 1, and sample-level KIR and HLA genotypes, population annotations, haplotype catalogs and supporting data are provided in Supplementary Data 1. Supplementary Tables S1–S11 and Supplementary Figures S1–S8 provide the supporting results. The underlying samples and population annotations are from the 1000 Genomes Project and are publicly available through the International Genome Sample Resource (IGSR). Original sequencing reads are not included. Further enquiries can be directed to the corresponding author.

## Ethics statement

This study analyzed publicly available, de-identified DNA samples and genomic data from the 1000 Genomes Project, distributed by the NHGRI Sample Repository for Human Genetic Research at the Coriell Institute for Medical Research. Informed consent for broad data sharing and secondary research use was obtained from participants by the original project. Ethical approval was not required for this secondary analysis in accordance with local legislation and institutional requirements.

## Author contributions

CWP and DEG conceived the study and designed the analyses. CWP and RW developed and performed the KIR typing assays. CWP, AT and YS performed sample handling, plate design and laboratory quality control. WCN and RW developed the sequence-analysis and allele-assignment pipeline and curated the allele calls. CWP performed the allele-frequency, diversity and differentiation analyses. CWP performed the haplotype estimation and prediction analyses. CWP, CF, YO, AI and RW performed the comparison with earlier 1000 Genomes typing. CWP and DEG prepared the figures and tables. DEG and CWP wrote the original draft. All authors reviewed and edited the manuscript and approved the submitted version. DEG supervised the study and acquired funding.

## Funding

Funding was derived in part from NIH grants R01DK132406 and UM1AI068614 and from the HLA Typing Core Resource operated by the Geraghty Laboratory at the Fred Hutchinson Cancer Center. Scisco Genetics, Inc. supplied KIR typing kits at cost.

## Acknowledgments

We thank the individuals who donated samples to the 1000 Genomes Project; this work would not have been possible without their participation. We acknowledge the 1000 Genomes Project Consortium and the International Genome Sample Resource (IGSR) at EMBL-EBI for curating and providing continued access to the panel and its associated population data (59, 60), and the NHGRI Sample Repository for Human Genetic Research at the Coriell Institute for Medical Research for distribution of the DNA samples. During preparation of this manuscript the authors used Claude (Anthropic) and Codex (OpenAI) to assist with drafting and editing of manuscript text, including figure and table legends, to assist with developing and checking analysis code, to check internal consistency across sections, and to suggest structural and analytical revisions. All laboratory work, data generation, statistical analysis and interpretation of results were performed by the authors. The authors reviewed and verified all content and take full responsibility for the published work.

## Conflict of interest

Authors CWP, WCN, RW, CF, AT, YS, YO, and DEG are affiliated with Scisco Genetics Inc., the manufacturer of the KIR and HLA typing assays used in this study. The remaining author declares that the research was conducted in the absence of any commercial or financial relationships that could be construed as a potential conflict of interest.

## Generative AI statement

During preparation of this manuscript the authors used Claude (Anthropic) and Codex (OpenAI) to assist with drafting and editing of manuscript text, including figure and table legends, to check internal consistency across sections, and to suggest structural and analytical revisions. All laboratory work, data generation, statistical analysis and interpretation of results were performed by the authors. The authors reviewed and verified all content and take full responsibility for the published work.

## Supplementary material

Supplementary Code 1, Supplementary Data 1, Supplementary Tables S1–S11 and Supplementary Figures S1–S8 accompany this article. Legends for the supplementary tables and figures are provided at the end of this manuscript.

## Supplementary table legends

**Table S1. Cohort composition.**

Individuals by super-population and named population, with percentages relative to the 2,298-person KIR cohort.

**Table S2. KIR allele richness.**

Retained richness after requiring at least five observed allele copies, reported overall and by super-population.

**Table S3. Published typing comparisons.**

Original published-call comparison overview, gene summaries and row-level records, retaining the original source definitions.

**Table S4. Structural ambiguity and resolution.**

Overall and super-population counts and proportions of unresolved, uniquely resolved and ambiguous individuals across exact haplotypes, collapsed combinations and regional motifs.

**Table S5. Exact haplotype frequencies.**

Weighted exact-architecture frequencies using compatible primary ambiguity preferences where available.

**Table S6. Population motif frequencies.**

Centromeric, telomeric and collapsed-combination frequencies by population, with ambiguity-aware weights.

**Table S7. Full motif-associated allele distributions.**

Observed allele counts and frequencies overall, by super-population and by named population among structurally unambiguous motif-homozygous carriers.

**Table S8. Super-population motif-allele source data.**

Fixed top-two allele selection, plotting data, and overall and super-population source counts.

**Table S9. Paired KIR and HLA analyses.**

Table S9 retains the original paired functional configurations, exact genotype summaries, allele and motif contexts, annotation lookup, eligibility, association families, linkage, power and calibration. Tabs S9AE–AS add the integrated allotype repertoire and its genomic-background comparison, including raw-call reconciliation. The index maps all 46 tabs and their scopes.

**Table S10. Gene-specific copy-number lookup intervals.**

Reference copy-number classes, normalized-ratio boundaries, informative exons and control-filtering summaries derived from 408 samples with known copy number. Ratios were normalized to KIR3DL3 under the assumption of two copies.

**Table S11. KIR validation in 48 reference samples.**

Locus-level copy-number and combined copy-number/allele concordance for the unedited software calls, provisional discordances requiring review, and the author-confirmed outcome after desktop review in 48 reference samples.

## Supplementary figure legends

**Figure S1.**
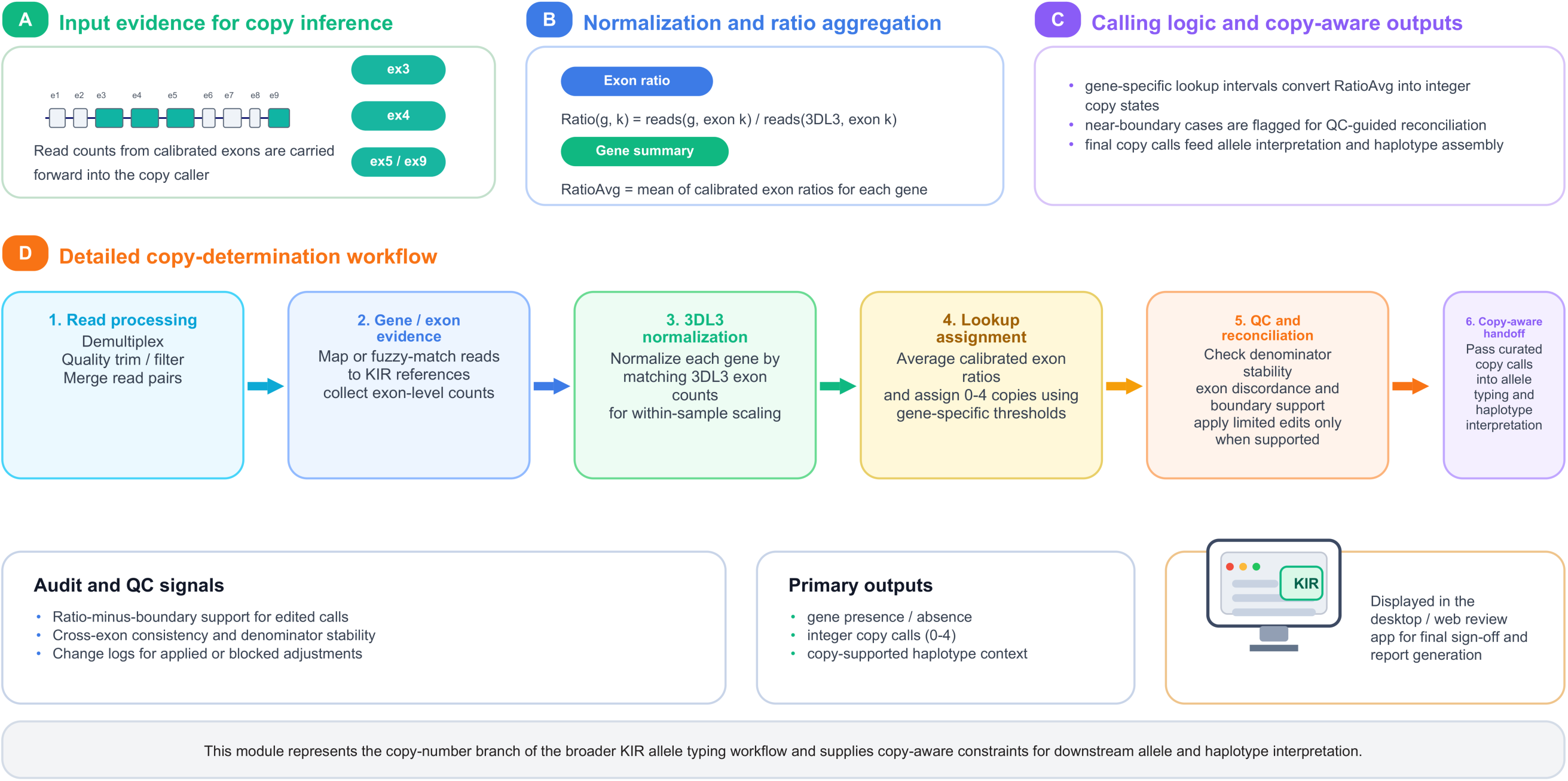
Copy-number workflow. Detailed schematic of the copy-number determination module embedded within the broader KIR allele-typing workflow, including read processing, exon evidence collection, KIR3DL3 normalization, ratio lookup calling, quality-control-guided reconciliation, and copy-aware handoff to allele interpretation.

**Figure S2.**
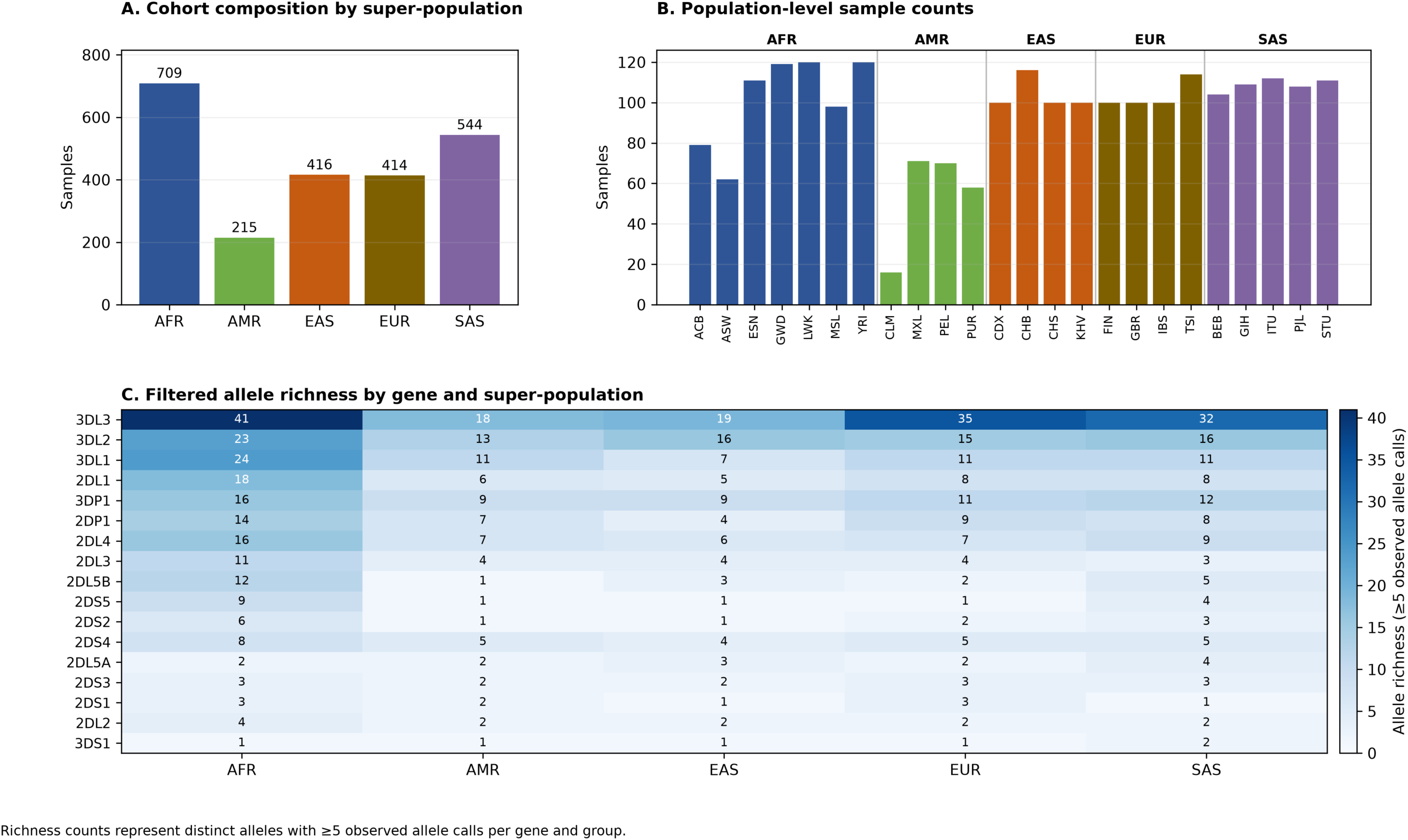
Cohort composition and filtered KIR allele richness. Panel A shows the number of individuals in each super-population. Panel B shows counts for all 24 populations grouped by super-population. Panel C shows filtered allele richness (minimum of five observed allele copies) across KIR genes and super-populations.

**Figure S3.**
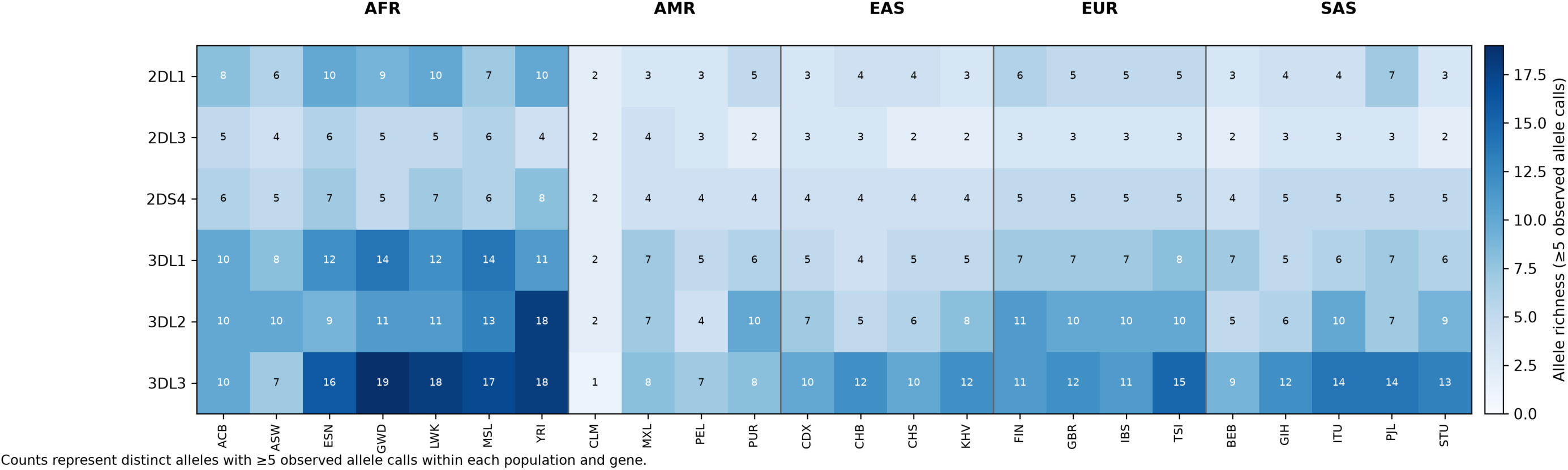
Population-specific KIR allele richness. Retained allele richness across 24 populations at KIR2DL1, KIR2DL3, KIR2DS4, KIR3DL1, KIR3DL2 and KIR3DL3. Richness required at least five observed allele copies within each population.

**Figure S4.**
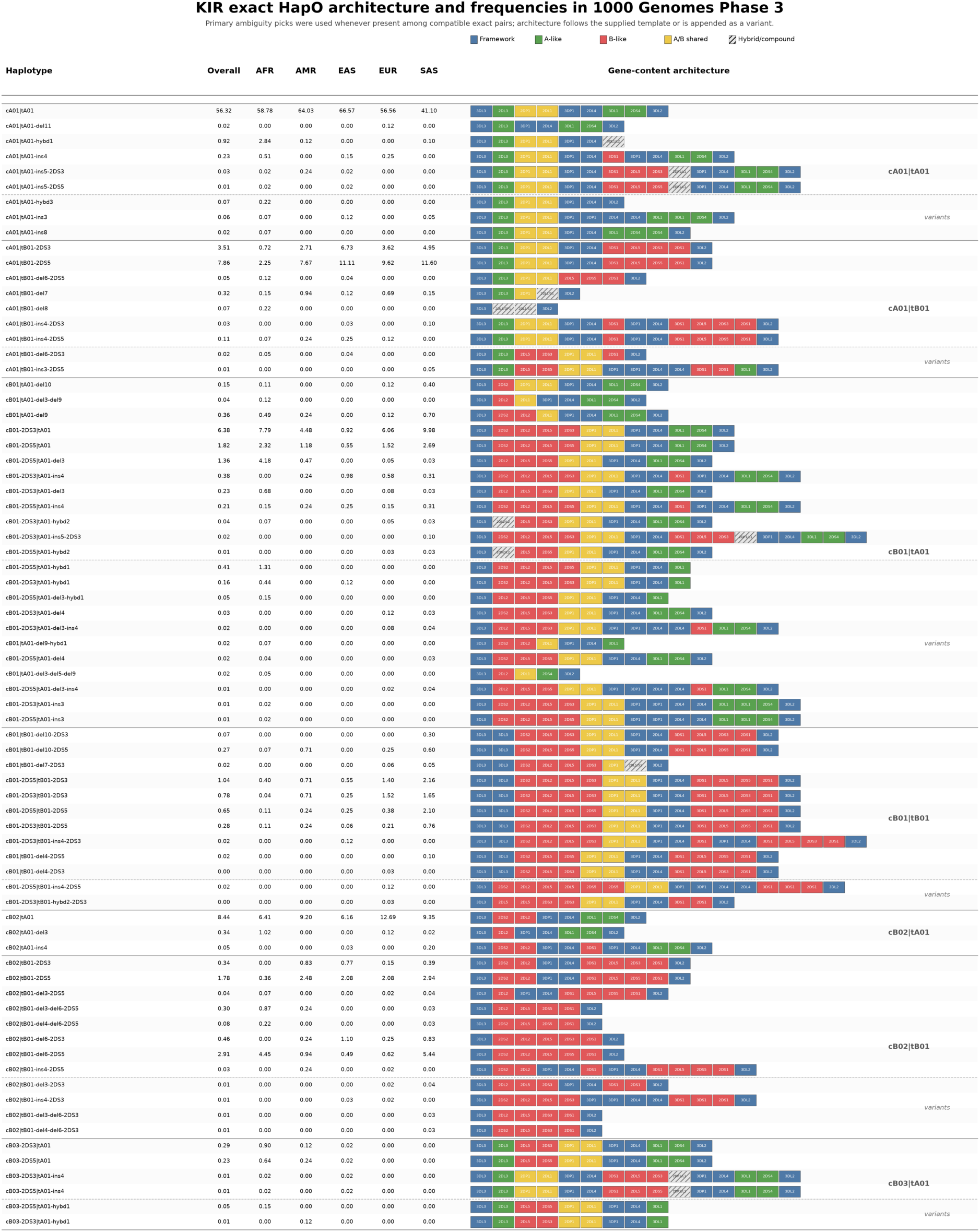
Weighted exact-haplotype architecture frequencies. Descriptive summary of weighted exact-haplotype frequencies using ambiguity-aware compatibility summaries together with a designated primary ambiguity pick when present among compatible exact solutions. The left columns report overall and super-population frequencies, and the right panel shows the corresponding gene-content architecture blocks grouped by major family background and variants. Exact labels are not uniquely resolved for most individuals (see Figure 3).

**Figure S5.**
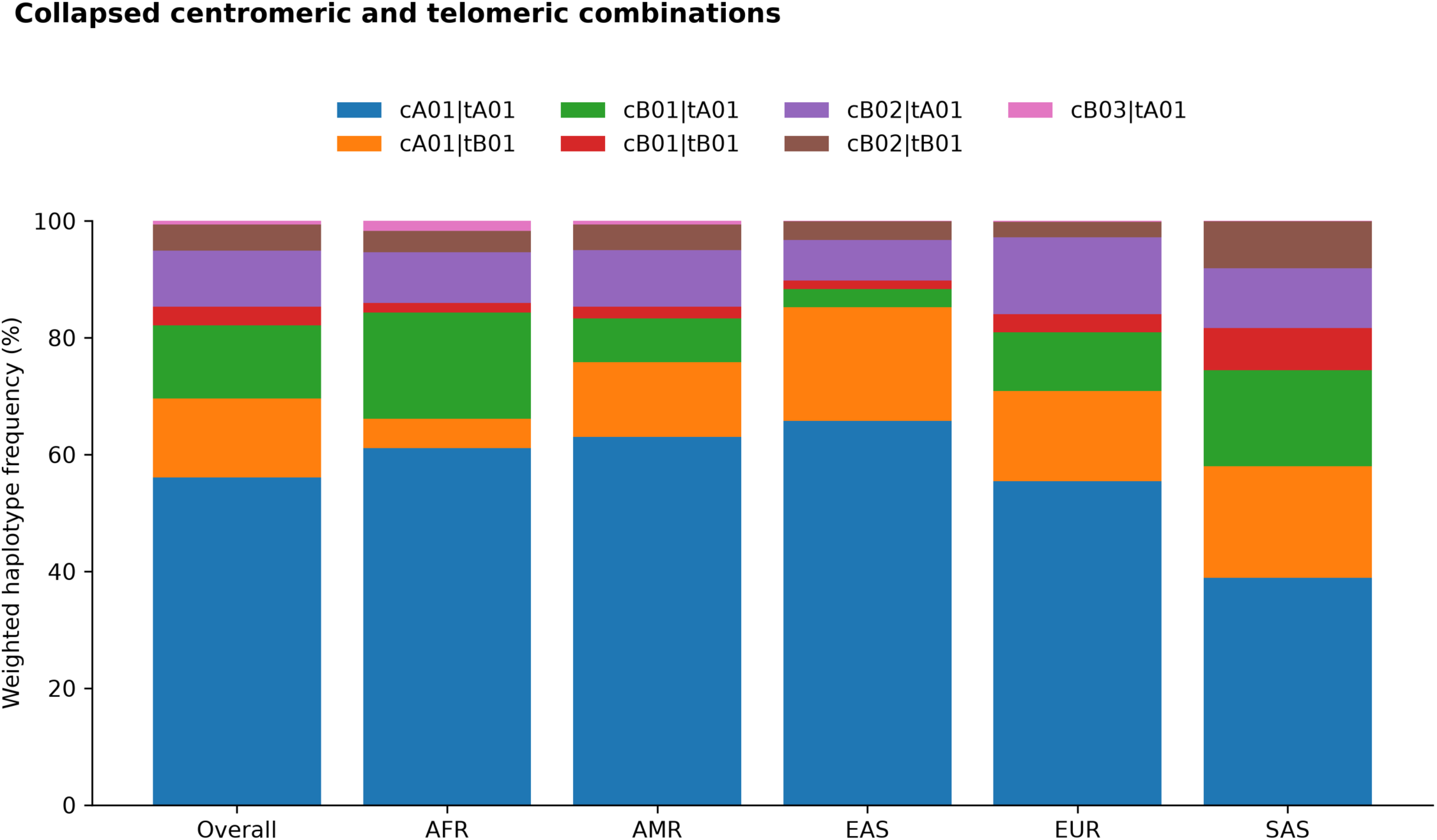
Weighted collapsed haplotype-combination frequencies. Stacked bar plots showing weighted frequencies of collapsed haplotype combinations across the cohort and super-populations after reduction of exact haplotype structures to centromeric|telomeric backgrounds.

**Figure S6.**
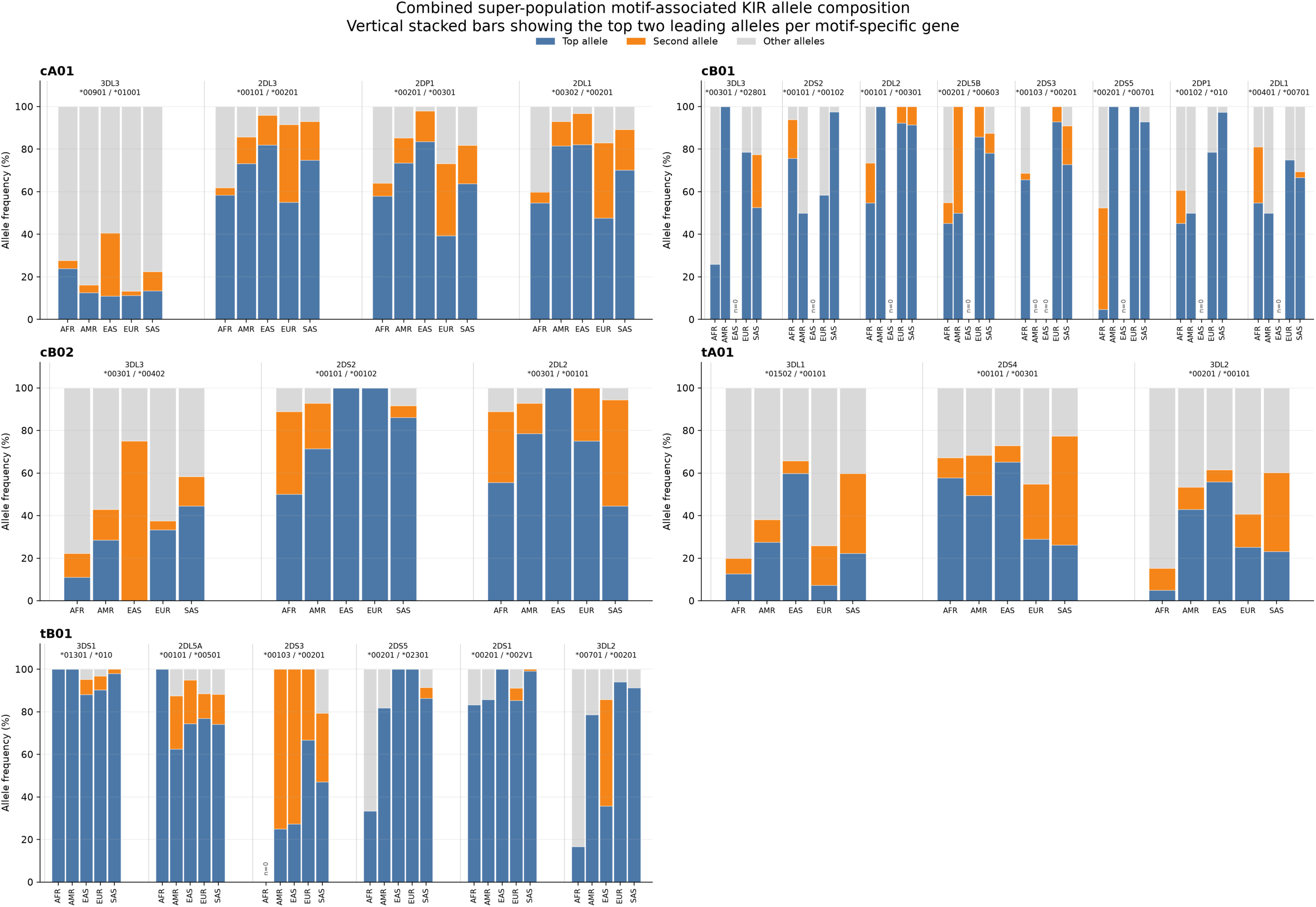
Super-population top-two motif-associated KIR allele composition. Vertical stacked bars show, for each motif-specific gene, the two overall-leading alleles across all motif-homozygous carriers, with the same two alleles then displayed across AFR, AMR, EAS, EUR, and SAS and all remaining alleles combined into an Other category. Blank bars labeled n=0 indicate no observed allele copies, rather than 100% Other.

**Figure S7.**
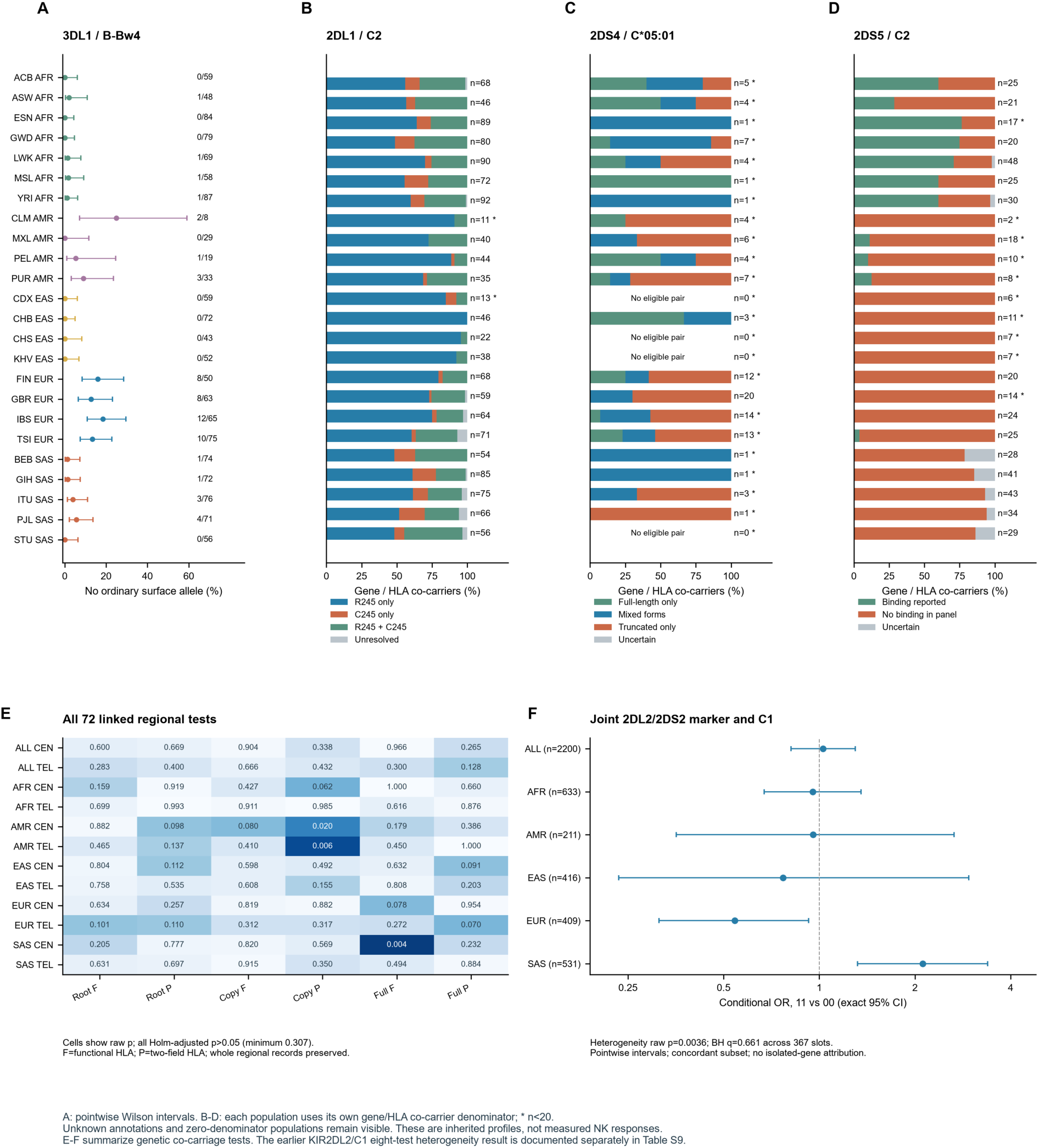
Population profiles and supporting KIR and HLA analyses. (A) Frequency of no ordinary surface-allele annotation among KIR3DL1/HLA-B Bw4 co-carriers across all 24 populations, with pointwise Wilson 95% intervals and explicit n/N counts. (B–D) The same population order is used for KIR2DL1/C2 sequence profiles, KIR2DS4/C*05:01 receptor forms and KIR2DS5/C2 binding annotations. Every denominator is shown; an asterisk indicates fewer than 20 co-carriers and zero denominators are labeled as having no eligible pair. (E) All 72 linked regional tests, with raw p values displayed for the pooled cohort and five super-populations, each divided into centromeric and telomeric arms. F and P denote functional and two-field HLA, respectively; root, copy-to-protein and protein-to-full comparisons remain in one Holm family. (F) Pointwise exact conditional intervals for the joint KIR2DL2/2DS2 11-versus-00 marker and C1 in the concordant subset; heterogeneity p = 0.0036 and BH q = 0.661 across 367 supporting tests. These contrasts do not isolate a KIR2DL2 receptor effect. Functional categories derive from inherited annotation, and intervals do not adjust for sampling design or documented relatedness.

**Figure S8.**
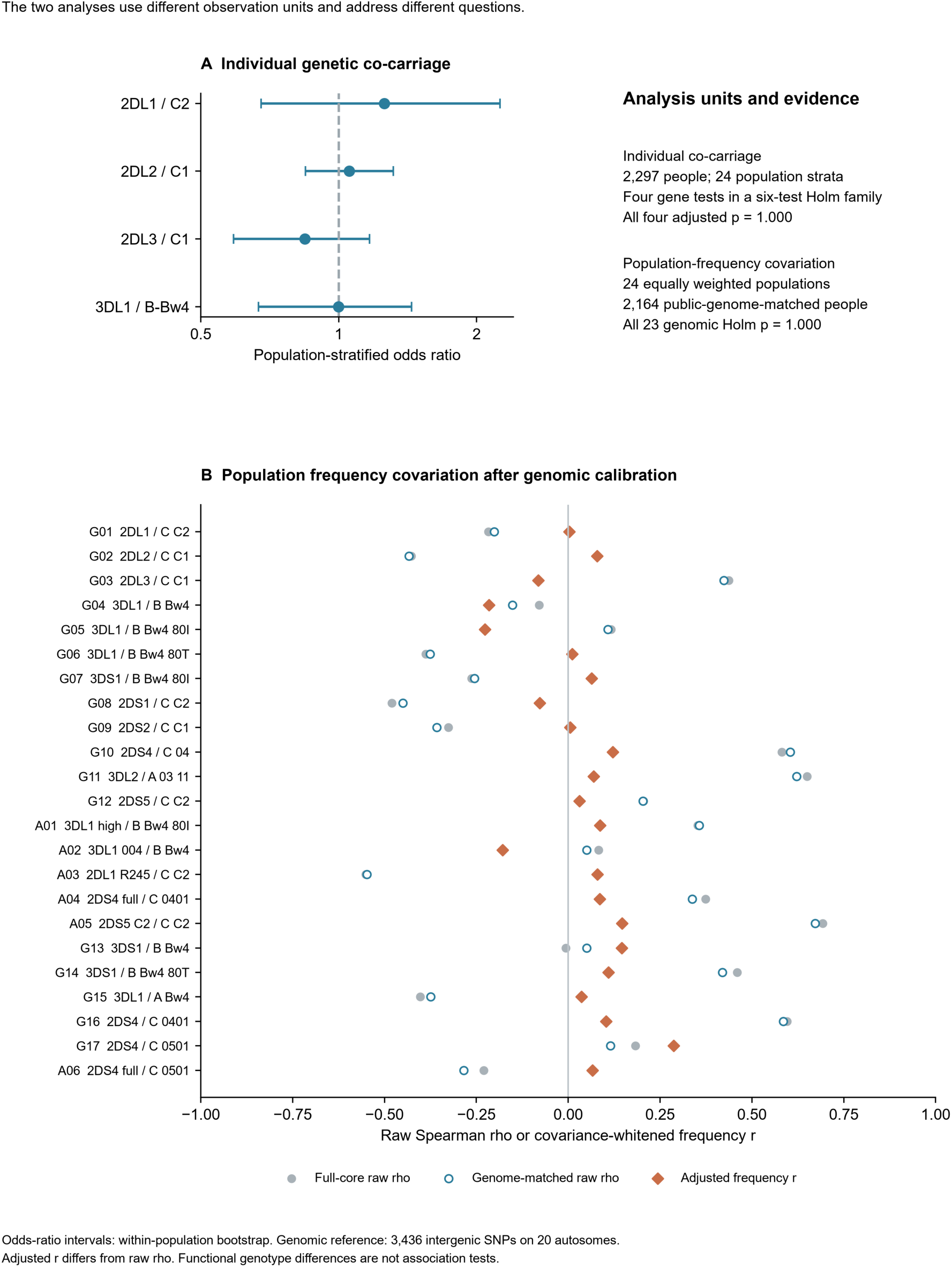
Genetic co-carriage and population frequency covariation. (A) Population-stratified Mantel–Haenszel odds ratios for four canonical gene-level receptor–ligand comparisons in the 2,297-person core. Intervals are pointwise 95% bootstrap intervals from 5,000 within-population resamples; all four p values are 1 after the six-test Holm correction. (B) Full-core raw Spearman correlations, raw correlations in the public-genome-matched subset and covariance-whitened frequency correlations for 23 reserved comparisons across 24 equally weighted populations. The genomic subset contains 2,164 individuals before class-specific annotation filters. Genomic background uses 3,436 intergenic SNPs from 20 autosomes, excluding chromosomes 6 and 19. Each empirical tail is the larger of two chromosome-separated calibration tails, followed by Holm across 23 slots; all adjusted p values are 1. Adjusted frequency r and raw Spearman rho are different statistics. The observation unit in A is an individual within population strata; in B it is a population-frequency pair.

