## Supplementary material for "Technology solutions for immune loci: integrated high-resolution KIR genotyping reveals allelic, structural, and KIR–HLA diversity across 24 human populations": all supplementary files: FigureS1_KIR_copy_number_workflow.pdf

### Figure S1. Copy-number calling within the KIR allele typing pipeline

#### A Input evidence for copy inference

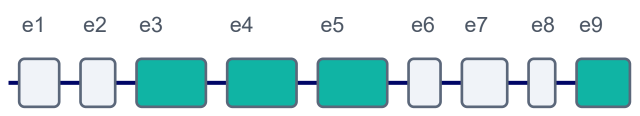


Read counts from calibrated exons are carried forward into the copy caller

ex3

ex4

ex5 / ex9

#### B Normalization and ratio aggregation

**Exon ratio**

Ratio(g, k) = reads(g, exon k) / reads(3DL3, exon k)

**Gene summary**

RatioAvg = mean of calibrated exon ratios for each gene

#### C Calling logic and copy-aware outputs

- gene-specific lookup intervals convert RatioAvg into integer copy states
- near-boundary cases are flagged for QC-guided reconciliation
- final copy calls feed allele interpretation and haplotype assembly

#### D Detailed copy-determination workflow

**1. Read processing**
Demultiplex
Quality trim / filter
Merge read pairs

→

**2. Gene / exon evidence**
Map or fuzzy-match reads
to KIR references
collect exon-level counts

→

**3. 3DL3 normalization**
Normalize each gene by
matching 3DL3 exon
counts
for within-sample scaling

→

**4. Lookup assignment**
Average calibrated exon
ratios
and assign 0-4 copies using
gene-specific thresholds

→

**5. QC and reconciliation**
Check denominator
stability
exon discordance and
boundary support
apply limited edits only
when supported

→

**6. Copy-aware handoff**
Pass curated
copy calls
into allele
typing and
haplotype
interpretation

##### Audit and QC signals

- Ratio-minus-boundary support for edited calls
- Cross-exon consistency and denominator stability
- Change logs for applied or blocked adjustments

##### Primary outputs

- gene presence / absence
- integer copy calls (0-4)
- copy-supported haplotype context

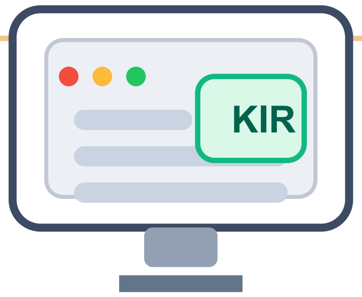


Displayed in the desktop / web review app for final sign-off and report generation

This module represents the copy-number branch of the broader KIR allele typing workflow and supplies copy-aware constraints for downstream allele and haplotype interpretation.
