## Supplementary material for "Technology solutions for immune loci: integrated high-resolution KIR genotyping reveals allelic, structural, and KIR–HLA diversity across 24 human populations": all supplementary files: FigureS2_KIR_cohort_and_richness.pdf

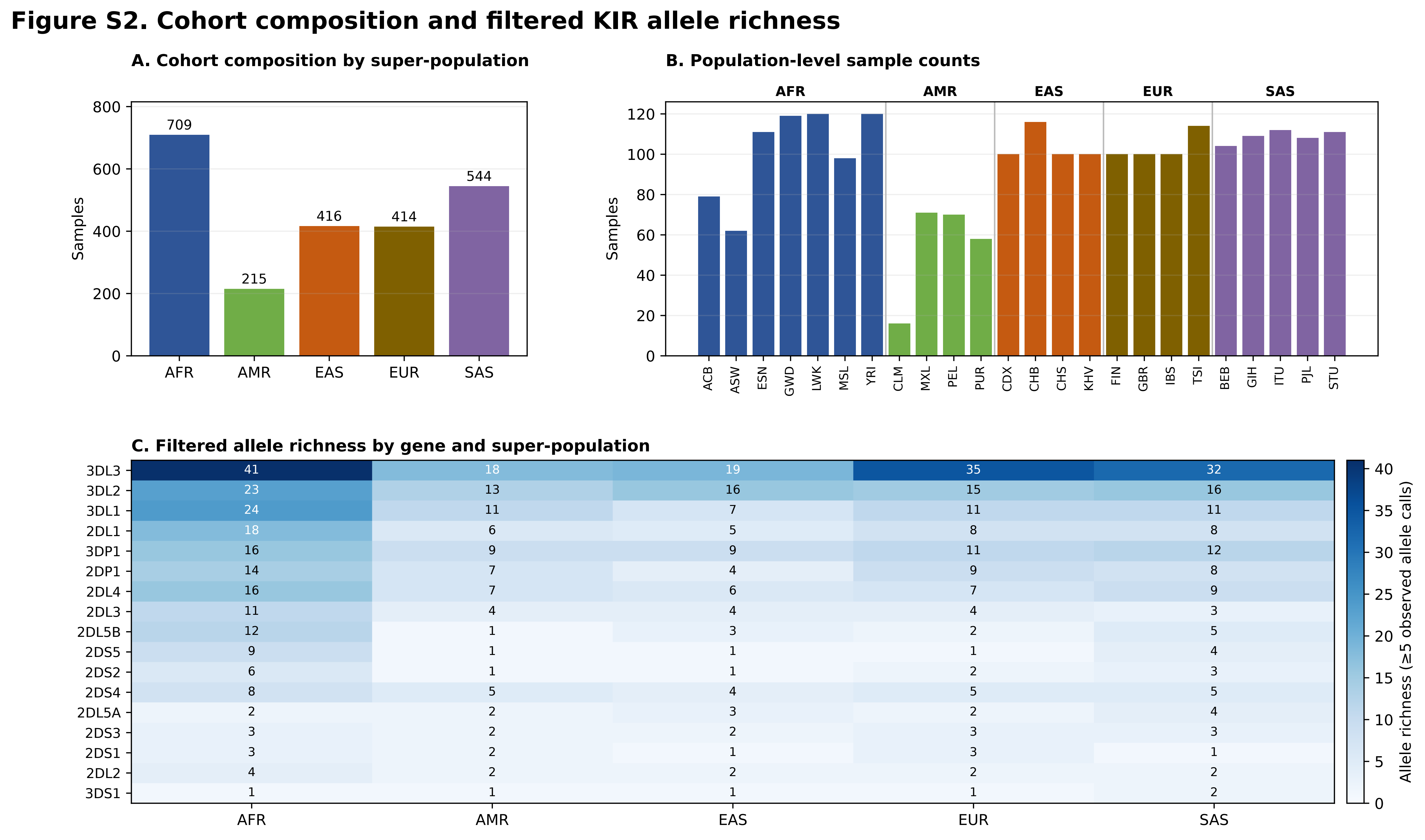


| Gene | AFR | AMR | EAS | EUR | SAS |
|---|---|---|---|---|---|
| 3DL3 | 41 | 18 | 19 | 35 | 32 |
| 3DL2 | 23 | 13 | 16 | 15 | 16 |
| 3DL1 | 24 | 11 | 7 | 11 | 11 |
| 2DL1 | 18 | 6 | 5 | 8 | 8 |
| 3DP1 | 16 | 9 | 9 | 11 | 12 |
| 2DP1 | 14 | 7 | 4 | 9 | 8 |
| 2DL4 | 16 | 7 | 6 | 7 | 9 |
| 2DL3 | 11 | 4 | 4 | 4 | 3 |
| 2DL5B | 12 | 1 | 3 | 2 | 5 |
| 2DS5 | 9 | 1 | 1 | 1 | 4 |
| 2DS2 | 6 | 1 | 1 | 2 | 3 |
| 2DS4 | 8 | 5 | 4 | 5 | 5 |
| 2DL5A | 2 | 2 | 3 | 2 | 4 |
| 2DS3 | 3 | 2 | 2 | 3 | 3 |
| 2DS1 | 3 | 2 | 1 | 3 | 1 |
| 2DL2 | 4 | 2 | 2 | 2 | 2 |
| 3DS1 | 1 | 1 | 1 | 1 | 2 |

Richness counts represent distinct alleles with ≥5 observed allele calls per gene and group.
