## Supplementary material for "Technology solutions for immune loci: integrated high-resolution KIR genotyping reveals allelic, structural, and KIR–HLA diversity across 24 human populations": all supplementary files: FigureS3_KIR_population_richness.pdf

### Figure S3. Population-level filtered allele richness for selected KIR genes

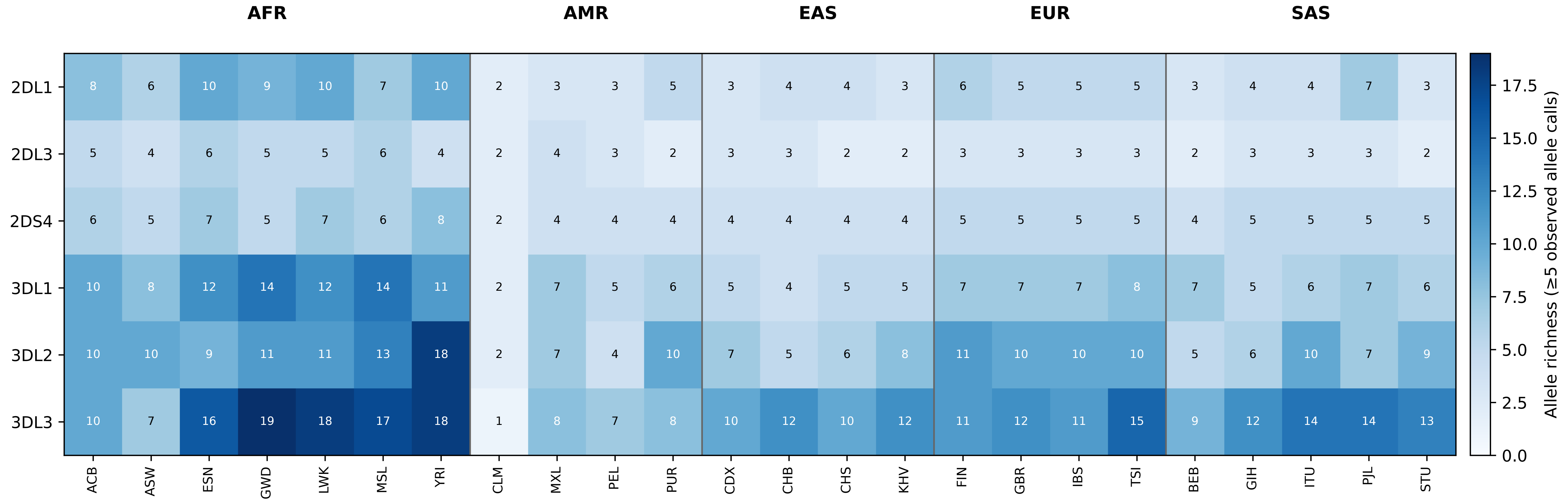


Counts represent distinct alleles with ≥5 observed allele calls within each population and gene.
