## Supplementary material for "Technology solutions for immune loci: integrated high-resolution KIR genotyping reveals allelic, structural, and KIR–HLA diversity across 24 human populations": all supplementary files: FigureS4_KIR_exact_haplotype_architectures.pdf

### KIR exact HapO architecture and frequencies in 1000 Genomes Phase 3

Primary ambiguity picks were used whenever present among compatible exact pairs; architecture follows the supplied template or is appended as a variant.

Legend: Framework · A-like · B-like · A/B shared · Hybrid/compound

| Haplotype | Overall | AFR | AMR | EAS | EUR | SAS | Gene-content architecture | |
|---|---|---|---|---|---|---|---|---|
| cA01\|tA01 | 56.32 | 58.78 | 64.03 | 66.57 | 56.56 | 41.10 | 3DL3 2DL3 2DP1 2DL1 3DP1 2DL4 3DL1 2DS4 3DL2 | |
| cA01\|tA01-del11 | 0.02 | 0.00 | 0.00 | 0.00 | 0.12 | 0.00 | 3DL3 2DL3 3DP1 2DL4 3DL1 2DS4 3DL2 | |
| cA01\|tA01-hybd1 | 0.92 | 2.84 | 0.12 | 0.00 | 0.00 | 0.10 | 3DL3 2DL3 2DP1 2DL1 3DP1 2DL4 3DL1/L2 | |
| cA01\|tA01-ins4 | 0.23 | 0.51 | 0.00 | 0.15 | 0.25 | 0.00 | 3DL3 2DL3 2DP1 2DL1 3DP1 2DL4 3DS1 3DP1 2DL4 3DL1 2DS4 3DL2 | |
| cA01\|tA01-ins5-2DS3 | 0.03 | 0.02 | 0.24 | 0.02 | 0.00 | 0.00 | 3DL3 2DL3 2DP1 2DL1 3DP1 2DL4 3DS1 2DL5 2DS3 2DP1/L1 3DP1 2DL4 3DL1 2DS4 3DL2 | cA01\|tA01 |
| cA01\|tA01-ins5-2DS5 | 0.01 | 0.02 | 0.00 | 0.02 | 0.00 | 0.00 | 3DL3 2DL3 2DP1 2DL1 3DP1 2DL4 3DS1 2DL5 2DS5 2DP1/L1 3DP1 2DL4 3DL1 2DS4 3DL2 | |
| cA01\|tA01-hybd3 | 0.07 | 0.22 | 0.00 | 0.00 | 0.00 | 0.00 | 3DL3 2DL3 2DP1 2DL1 3DP1 2DL4 3DL2 | |
| cA01\|tA01-ins3 | 0.06 | 0.07 | 0.00 | 0.12 | 0.00 | 0.05 | 3DL3 2DL3 2DP1 2DL1 3DP1 3DP1 2DL4 2DL4 3DL1 3DL1 2DS4 3DL2 | *variants* |
| cA01\|tA01-ins8 | 0.02 | 0.07 | 0.00 | 0.00 | 0.00 | 0.00 | 3DL3 2DL3 2DP1 2DL1 3DP1 2DL4 3DL1 2DS4 2DS4 3DL2 | |
| cA01\|tB01-2DS3 | 3.51 | 0.72 | 2.71 | 6.73 | 3.62 | 4.95 | 3DL3 2DL3 2DP1 2DL1 3DP1 2DL4 3DS1 2DL5 2DS3 2DS1 3DL2 | |
| cA01\|tB01-2DS5 | 7.86 | 2.25 | 7.67 | 11.11 | 9.62 | 11.60 | 3DL3 2DL3 2DP1 2DL1 3DP1 2DL4 3DS1 2DL5 2DS5 2DS1 3DL2 | |
| cA01\|tB01-del6-2DS5 | 0.05 | 0.12 | 0.00 | 0.04 | 0.00 | 0.00 | 3DL3 2DL3 2DP1 2DL1 2DL5 2DS5 2DS1 3DL2 | |
| cA01\|tB01-del7 | 0.32 | 0.15 | 0.94 | 0.12 | 0.69 | 0.15 | 3DL3 2DL3 2DP1 2DL1/S1 3DL2 | |
| cA01\|tB01-del8 | 0.07 | 0.22 | 0.00 | 0.00 | 0.00 | 0.00 | 3DL3 2DL3/DP1 2DL1/S1 3DL2 | cA01\|tB01 |
| cA01\|tB01-ins4-2DS3 | 0.03 | 0.00 | 0.00 | 0.03 | 0.00 | 0.10 | 3DL3 2DL3 2DP1 2DL1 3DP1 2DL4 3DS1 3DP1 2DL4 3DS1 2DL5 2DS3 2DS1 3DL2 | |
| cA01\|tB01-ins4-2DS5 | 0.11 | 0.07 | 0.24 | 0.25 | 0.12 | 0.00 | 3DL3 2DL3 2DP1 2DL1 3DP1 2DL4 3DS1 3DP1 2DL4 3DS1 2DL5 2DS5 2DS1 3DL2 | |
| cA01\|tB01-del6-2DS3 | 0.02 | 0.05 | 0.00 | 0.04 | 0.00 | 0.00 | 3DL3 2DL3 2DL5 2DS3 2DP1 2DL1 2DS1 3DL2 | *variants* |
| cA01\|tB01-ins3-2DS5 | 0.01 | 0.00 | 0.00 | 0.00 | 0.00 | 0.05 | 3DL3 2DL3 2DL5 2DS5 2DP1 2DL1 3DP1 3DP1 2DL4 2DL4 3DS1 2DS1 3DL1 3DL2 | |
| cB01\|tA01-del10 | 0.15 | 0.11 | 0.00 | 0.00 | 0.12 | 0.40 | 3DL3 2DS2 2DP1 2DL1 3DP1 2DL4 3DL1 2DS4 3DL2 | |
| cB01\|tA01-del3-del9 | 0.04 | 0.12 | 0.00 | 0.00 | 0.00 | 0.00 | 3DL3 2DL2 2DL1 3DP1 2DL4 3DL1 2DS4 3DL2 | |
| cB01\|tA01-del9 | 0.36 | 0.49 | 0.24 | 0.00 | 0.12 | 0.70 | 3DL3 2DS2 2DL2 2DL1 3DP1 2DL4 3DL1 2DS4 3DL2 | |
| cB01-2DS3\|tA01 | 6.38 | 7.79 | 4.48 | 0.92 | 6.06 | 9.98 | 3DL3 2DS2 2DL2 2DL5 2DS3 2DP1 2DL1 3DP1 2DL4 3DL1 2DS4 3DL2 | |
| cB01-2DS5\|tA01 | 1.82 | 2.32 | 1.18 | 0.55 | 1.52 | 2.69 | 3DL3 2DS2 2DL2 2DL5 2DS5 2DP1 2DL1 3DP1 2DL4 3DL1 2DS4 3DL2 | |
| cB01-2DS5\|tA01-del3 | 1.36 | 4.18 | 0.47 | 0.00 | 0.05 | 0.03 | 3DL3 2DL2 2DL5 2DS5 2DP1 2DL1 3DP1 2DL4 3DL1 2DS4 3DL2 | |
| cB01-2DS3\|tA01-ins4 | 0.38 | 0.00 | 0.24 | 0.98 | 0.58 | 0.31 | 3DL3 2DS2 2DL2 2DL5 2DS3 2DP1 2DL1 3DP1 2DL4 3DS1 3DP1 2DL4 3DL1 2DS4 3DL2 | |
| cB01-2DS3\|tA01-del3 | 0.23 | 0.68 | 0.00 | 0.00 | 0.08 | 0.03 | 3DL3 2DL2 2DL5 2DS3 2DP1 2DL1 3DP1 2DL4 3DL1 2DS4 3DL2 | |
| cB01-2DS5\|tA01-ins4 | 0.21 | 0.15 | 0.24 | 0.25 | 0.15 | 0.31 | 3DL3 2DS2 2DL2 2DL5 2DS5 2DP1 2DL1 3DP1 2DL4 3DS1 3DP1 2DL4 3DL1 2DS4 3DL2 | |
| cB01-2DS3\|tA01-hybd2 | 0.04 | 0.07 | 0.00 | 0.00 | 0.05 | 0.03 | 3DL3 2DS2/L2 2DL5 2DS3 2DP1 2DL1 3DP1 2DL4 3DL1 2DS4 3DL2 | |
| cB01-2DS3\|tA01-ins5-2DS3 | 0.02 | 0.00 | 0.00 | 0.00 | 0.00 | 0.10 | 3DL3 2DS2 2DL2 2DL5 2DS3 2DP1 2DL1 3DP1 2DL4 3DS1 2DL5 2DS3 2DP1/L1 3DP1 2DL4 3DL1 2DS4 3DL2 | |
| cB01-2DS5\|tA01-hybd2 | 0.01 | 0.00 | 0.00 | 0.00 | 0.03 | 0.03 | 3DL3 2DS2/L2 2DL5 2DS5 2DP1 2DL1 3DP1 2DL4 3DL1 2DS4 3DL2 | cB01\|tA01 |
| cB01-2DS5\|tA01-hybd1 | 0.41 | 1.31 | 0.00 | 0.00 | 0.00 | 0.00 | 3DL3 2DS2 2DL2 2DL5 2DS5 2DP1 2DL1 3DP1 2DL4 3DL1 | |
| cB01-2DS3\|tA01-hybd1 | 0.16 | 0.44 | 0.00 | 0.12 | 0.00 | 0.00 | 3DL3 2DS2 2DL2 2DL5 2DS3 2DP1 2DL1 3DP1 2DL4 3DL1 | |
| cB01-2DS5\|tA01-del3-hybd1 | 0.05 | 0.15 | 0.00 | 0.00 | 0.00 | 0.00 | 3DL3 2DL2 2DL5 2DS5 2DP1 2DL1 3DP1 2DL4 3DL1 | |
| cB01-2DS3\|tA01-del4 | 0.03 | 0.00 | 0.00 | 0.00 | 0.12 | 0.03 | 3DL3 2DS2 2DL5 2DS3 2DP1 2DL1 3DP1 2DL4 3DL1 2DS4 3DL2 | |
| cB01-2DS3\|tA01-del3-ins4 | 0.02 | 0.00 | 0.00 | 0.00 | 0.08 | 0.04 | 3DL3 2DL2 2DL5 2DS3 2DP1 2DL1 3DP1 3DP1 2DL4 2DL4 3DS1 3DL1 2DS4 3DL2 | |
| cB01\|tA01-del9-hybd1 | 0.02 | 0.07 | 0.00 | 0.00 | 0.00 | 0.00 | 3DL3 2DS2 2DL2 2DL1 3DP1 2DL4 3DL1 | *variants* |
| cB01-2DS5\|tA01-del4 | 0.02 | 0.04 | 0.00 | 0.00 | 0.00 | 0.03 | 3DL3 2DS2 2DL5 2DS5 2DP1 2DL1 3DP1 2DL4 3DL1 2DS4 3DL2 | |
| cB01\|tA01-del3-del5-del9 | 0.02 | 0.05 | 0.00 | 0.00 | 0.00 | 0.00 | 3DL3 2DL2 2DL1 2DS4 3DL2 | |
| cB01-2DS5\|tA01-del3-ins4 | 0.01 | 0.00 | 0.00 | 0.00 | 0.02 | 0.04 | 3DL3 2DL2 2DL5 2DS5 2DP1 2DL1 3DP1 3DP1 2DL4 2DL4 3DS1 3DL1 2DS4 3DL2 | |
| cB01-2DS3\|tA01-ins3 | 0.01 | 0.02 | 0.00 | 0.00 | 0.00 | 0.00 | 3DL3 2DS2 2DL2 2DL5 2DS3 2DP1 2DL1 3DP1 3DP1 2DL4 2DL4 3DL1 3DL1 2DS4 3DL2 | |
| cB01-2DS5\|tA01-ins3 | 0.01 | 0.02 | 0.00 | 0.00 | 0.00 | 0.00 | 3DL3 2DS2 2DL2 2DL5 2DS5 2DP1 2DL1 3DP1 3DP1 2DL4 2DL4 3DL1 3DL1 2DS4 3DL2 | |
| cB01\|tB01-del10-2DS3 | 0.07 | 0.00 | 0.00 | 0.00 | 0.00 | 0.30 | 3DL3 3DL3 2DS2 2DL5 2DS3 2DP1 2DL1 3DP1 2DL4 3DS1 2DL5 2DS3 2DS1 3DL2 | |
| cB01\|tB01-del10-2DS5 | 0.27 | 0.07 | 0.71 | 0.00 | 0.25 | 0.60 | 3DL3 3DL3 2DS2 2DL5 2DS5 2DP1 2DL1 3DP1 2DL4 3DS1 2DL5 2DS5 2DS1 3DL2 | |
| cB01\|tB01-del7-2DS3 | 0.02 | 0.00 | 0.00 | 0.00 | 0.06 | 0.05 | 3DL3 3DL3 2DS2 2DL2 2DL5 2DS3 2DP1 2DL1/S1 3DL2 | |
| cB01-2DS5\|tB01-2DS3 | 1.04 | 0.40 | 0.71 | 0.55 | 1.40 | 2.16 | 3DL3 3DL3 2DS2 2DL2 2DL5 2DS3 2DP1 2DL1 3DP1 2DL4 3DS1 2DL5 2DS5 2DS1 3DL2 | |
| cB01-2DS3\|tB01-2DS3 | 0.78 | 0.04 | 0.71 | 0.25 | 1.52 | 1.65 | 3DL3 3DL3 2DS2 2DL2 2DL5 2DS3 2DP1 2DL1 3DP1 2DL4 3DS1 2DL5 2DS3 2DS1 3DL2 | |
| cB01-2DS5\|tB01-2DS5 | 0.65 | 0.11 | 0.24 | 0.25 | 0.38 | 2.10 | 3DL3 3DL3 2DS2 2DL2 2DL5 2DS5 2DP1 2DL1 3DP1 2DL4 3DS1 2DL5 2DS5 2DS1 3DL2 | |
| cB01-2DS3\|tB01-2DS5 | 0.28 | 0.11 | 0.24 | 0.06 | 0.21 | 0.76 | 3DL3 3DL3 2DS2 2DL2 2DL5 2DS3 2DP1 2DL1 3DP1 2DL4 3DS1 2DL5 2DS5 2DS1 3DL2 | cB01\|tB01 |
| cB01-2DS3\|tB01-ins4-2DS3 | 0.02 | 0.00 | 0.00 | 0.12 | 0.00 | 0.00 | 3DL3 3DL3 2DS2 2DL2 2DL5 2DS3 2DP1 2DL1 3DP1 2DL4 3DS1 3DP1 2DL4 3DS1 2DL5 2DS3 2DS1 3DL2 | |
| cB01\|tB01-del4-2DS5 | 0.02 | 0.00 | 0.00 | 0.00 | 0.00 | 0.10 | 3DL3 3DL3 2DS2 2DL5 2DS5 2DP1 2DL1 3DP1 2DL4 3DS1 2DL5 2DS5 2DS1 3DL2 | |
| cB01\|tB01-del4-2DS3 | 0.00 | 0.00 | 0.00 | 0.00 | 0.03 | 0.00 | 3DL3 3DL3 2DS2 2DL5 2DS3 2DP1 2DL1 3DP1 2DL4 3DS1 2DL5 2DS3 2DS1 3DL2 | |
| cB01-2DS5\|tB01-ins4-2DS5 | 0.02 | 0.00 | 0.00 | 0.00 | 0.12 | 0.00 | 3DL3 2DS2 2DL2 2DL5 2DL5 2DS5 2DS5 2DP1 2DL1 3DP1 3DP1 2DL4 2DL4 3DS1 3DS1 2DS1 3DL2 | *variants* |
| cB01-2DS3\|tB01-hybd2-2DS3 | 0.00 | 0.00 | 0.00 | 0.00 | 0.03 | 0.00 | 3DL3 2DL5 2DL5 2DS3 2DS3 2DP1 2DL1 3DP1 2DL4 3DS1 2DS1 3DL2 | |
| cB02\|tA01 | 8.44 | 6.41 | 9.20 | 6.16 | 12.69 | 9.35 | 3DL3 2DS2 2DL2 3DP1 2DL4 3DL1 2DS4 3DL2 | |
| cB02\|tA01-del3 | 0.34 | 1.02 | 0.00 | 0.00 | 0.12 | 0.02 | 3DL3 2DL2 3DP1 2DL4 3DL1 2DS4 3DL2 | cB02\|tA01 |
| cB02\|tA01-ins4 | 0.05 | 0.00 | 0.00 | 0.03 | 0.00 | 0.20 | 3DL3 2DS2 2DL2 3DP1 2DL4 3DS1 3DP1 2DL4 3DL1 2DS4 3DL2 | |
| cB02\|tB01-2DS3 | 0.34 | 0.00 | 0.83 | 0.77 | 0.15 | 0.39 | 3DL3 2DS2 2DL2 3DP1 2DL4 3DS1 2DL5 2DS3 2DS1 3DL2 | |
| cB02\|tB01-2DS5 | 1.78 | 0.36 | 2.48 | 2.08 | 2.08 | 2.94 | 3DL3 2DS2 2DL2 3DP1 2DL4 3DS1 2DL5 2DS5 2DS1 3DL2 | |
| cB02\|tB01-del3-2DS5 | 0.04 | 0.07 | 0.00 | 0.00 | 0.02 | 0.04 | 3DL3 2DL2 3DP1 2DL4 3DS1 2DL5 2DS5 2DS1 3DL2 | |
| cB02\|tB01-del3-del6-2DS5 | 0.30 | 0.87 | 0.24 | 0.00 | 0.00 | 0.03 | 3DL3 2DL2 2DL5 2DS5 2DS1 3DL2 | |
| cB02\|tB01-del4-del6-2DS5 | 0.08 | 0.22 | 0.00 | 0.00 | 0.00 | 0.03 | 3DL3 2DS2 2DL5 2DS5 2DS1 3DL2 | |
| cB02\|tB01-del6-2DS3 | 0.46 | 0.00 | 0.24 | 1.10 | 0.25 | 0.83 | 3DL3 2DS2 2DL2 2DL5 2DS3 2DS1 3DL2 | cB02\|tB01 |
| cB02\|tB01-del6-2DS5 | 2.91 | 4.45 | 0.94 | 0.49 | 0.62 | 5.44 | 3DL3 2DS2 2DL2 2DL5 2DS5 2DS1 3DL2 | |
| cB02\|tB01-ins4-2DS5 | 0.03 | 0.00 | 0.24 | 0.00 | 0.02 | 0.00 | 3DL3 2DS2 2DL2 3DP1 2DL4 3DS1 3DP1 2DL4 3DS1 2DL5 2DS5 2DS1 3DL2 | |
| cB02\|tB01-del3-2DS3 | 0.01 | 0.00 | 0.00 | 0.00 | 0.02 | 0.04 | 3DL3 2DL2 2DL5 2DS3 3DP1 2DL4 3DS1 2DS1 3DL2 | |
| cB02\|tB01-ins4-2DS3 | 0.01 | 0.00 | 0.00 | 0.03 | 0.02 | 0.00 | 3DL3 2DS2 2DL2 2DL5 2DS3 3DP1 3DP1 2DL4 2DL4 3DS1 3DS1 2DS1 3DL2 | *variants* |
| cB02\|tB01-del3-del6-2DS3 | 0.01 | 0.00 | 0.00 | 0.00 | 0.00 | 0.03 | 3DL3 2DL2 2DL5 2DS3 2DS1 3DL2 | |
| cB02\|tB01-del4-del6-2DS3 | 0.01 | 0.00 | 0.00 | 0.00 | 0.00 | 0.03 | 3DL3 2DS2 2DL5 2DS3 2DS1 3DL2 | |
| cB03-2DS3\|tA01 | 0.29 | 0.90 | 0.12 | 0.02 | 0.00 | 0.00 | 3DL3 2DL3 2DL5 2DS3 2DP1 2DL1 3DP1 2DL4 3DL1 2DS4 3DL2 | |
| cB03-2DS5\|tA01 | 0.23 | 0.64 | 0.24 | 0.02 | 0.00 | 0.00 | 3DL3 2DL3 2DL5 2DS5 2DP1 2DL1 3DP1 2DL4 3DL1 2DS4 3DL2 | |
| cB03-2DS3\|tA01-ins4 | 0.01 | 0.02 | 0.00 | 0.02 | 0.00 | 0.00 | 3DL3 2DL3 2DP1 2DL1 3DP1 2DL4 3DS1 2DL5 2DS3 2DP1/L1 3DP1 2DL4 3DL1 2DS4 3DL2 | cB03\|tA01 |
| cB03-2DS5\|tA01-ins4 | 0.01 | 0.02 | 0.00 | 0.02 | 0.00 | 0.00 | 3DL3 2DL3 2DP1 2DL1 3DP1 2DL4 3DS1 2DL5 2DS5 2DP1/L1 3DP1 2DL4 3DL1 2DS4 3DL2 | |
| cB03-2DS5\|tA01-hybd1 | 0.05 | 0.15 | 0.00 | 0.00 | 0.00 | 0.00 | 3DL3 2DL3 2DL5 2DS5 2DP1 2DL1 3DP1 2DL4 3DL1 | *variants* |
| cB03-2DS3\|tA01-hybd1 | 0.01 | 0.00 | 0.12 | 0.00 | 0.00 | 0.00 | 3DL3 2DL3 2DL5 2DS3 2DP1 2DL1 3DP1 2DL4 3DL1 | |
