## Supplementary material for "Technology solutions for immune loci: integrated high-resolution KIR genotyping reveals allelic, structural, and KIR–HLA diversity across 24 human populations": all supplementary files: FigureS7_KIR_HLA_population_support.pdf

**Figure S7. Population profiles and supporting KIR and HLA analyses**

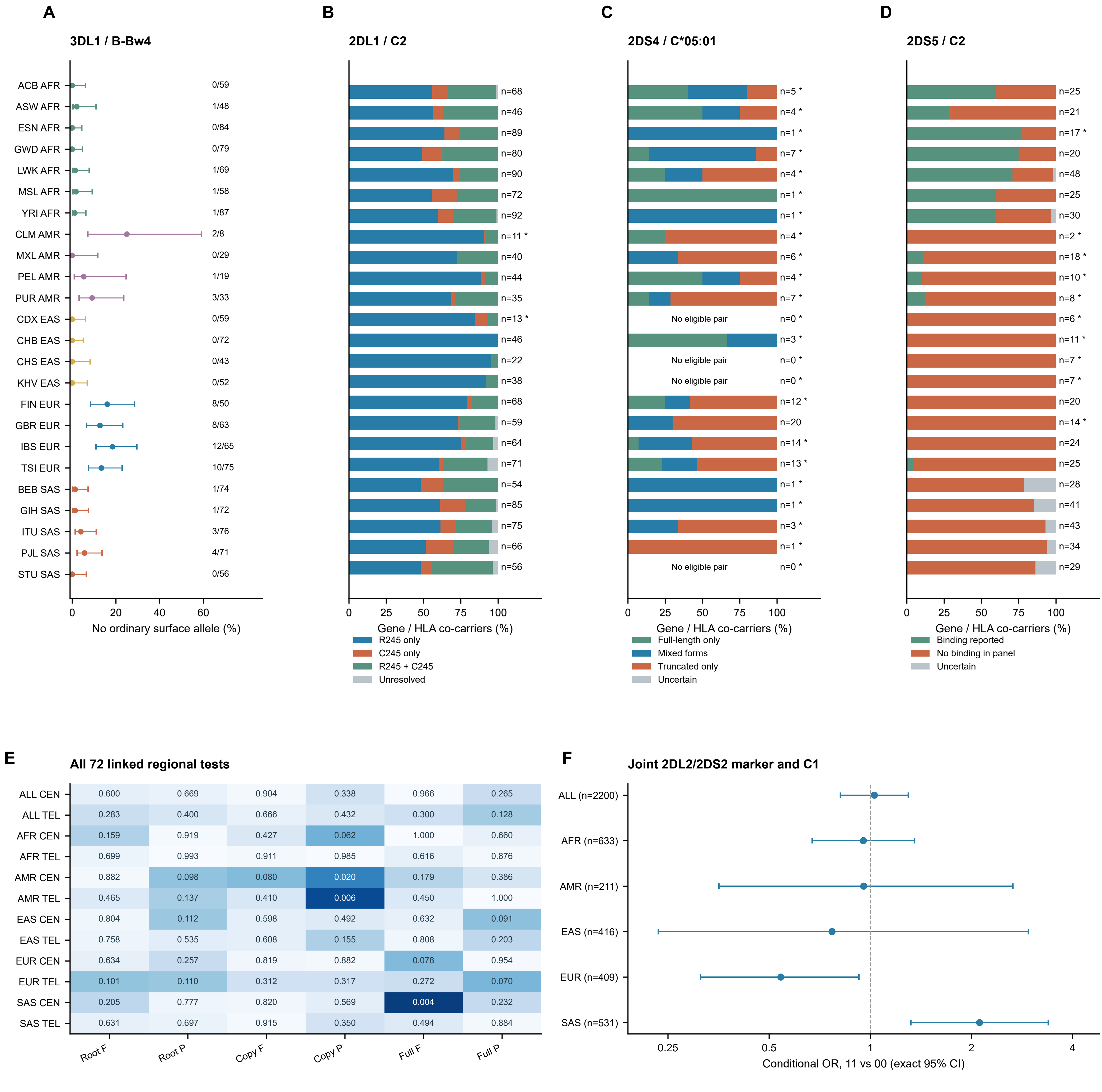


Cells show raw p; all Holm-adjusted p>0.05 (minimum 0.307).
F=functional HLA; P=two-field HLA; whole regional records preserved.

Heterogeneity raw p=0.0036; BH q=0.661 across 367 slots.
Pointwise intervals; concordant subset; no isolated-gene attribution.

A: pointwise Wilson intervals. B-D: each population uses its own gene/HLA co-carrier denominator; * n<20.
Unknown annotations and zero-denominator populations remain visible. These are inherited profiles, not measured NK responses.
E-F summarize genetic co-carriage tests. The earlier KIR2DL2/C1 eight-test heterogeneity result is documented separately in Table S9.
