## Supplementary material for "Technology solutions for immune loci: integrated high-resolution KIR genotyping reveals allelic, structural, and KIR–HLA diversity across 24 human populations": all supplementary files: FigureS8_KIR_HLA_association_calibration.pdf

### Supplementary Figure S8 Genetic co-carriage and population covariation

The two analyses use different observation units and address different questions.

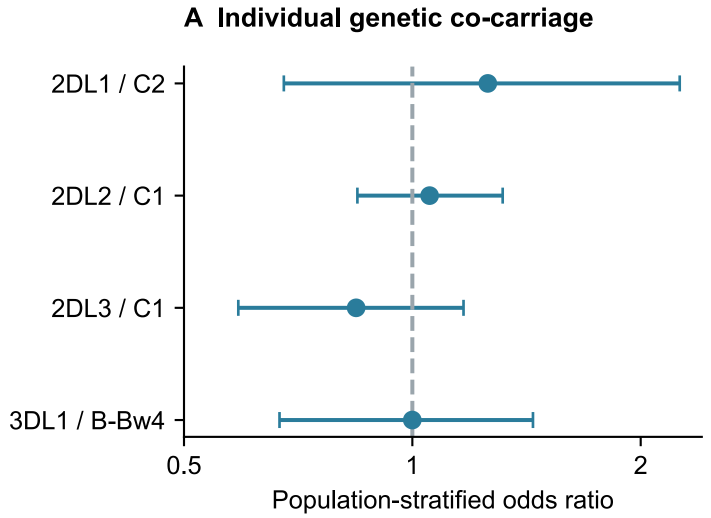


#### Analysis units and evidence

Individual co-carriage
2,297 people; 24 population strata
Four gene tests in a six-test Holm family
All four adjusted p = 1.000

Population-frequency covariation
24 equally weighted populations
2,164 public-genome-matched people
All 23 genomic Holm p = 1.000

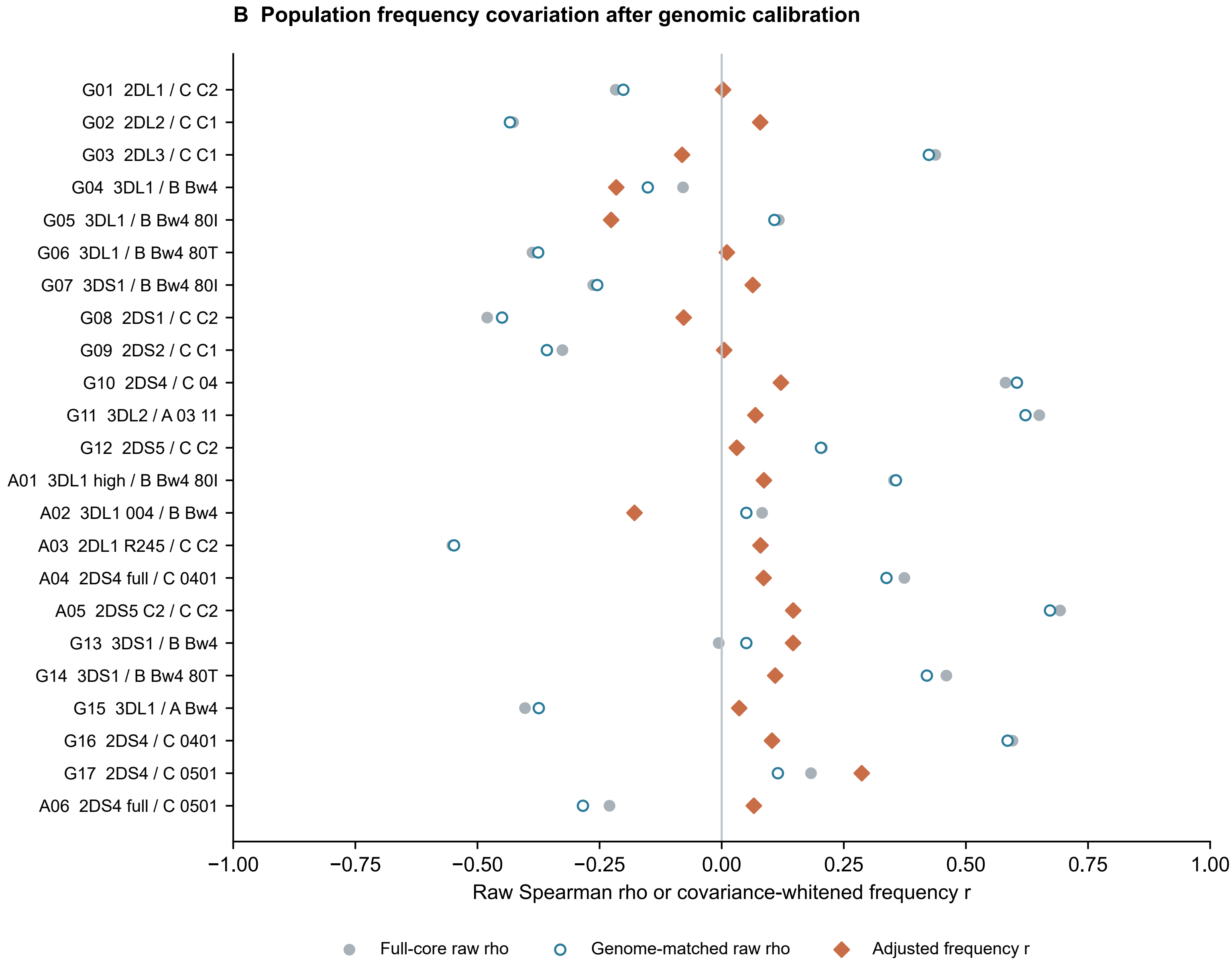


Odds-ratio intervals: within-population bootstrap. Genomic reference: 3,436 intergenic SNPs on 20 autosomes.
Adjusted r differs from raw rho. Functional genotype differences are not association tests.
