## Supplementary figures and images for "Technology solutions for immune loci: integrated high-resolution KIR genotyping reveals allelic, structural, and KIR–HLA diversity across 24 human populations"

### FigureS1_KIR_copy_number_workflow.png

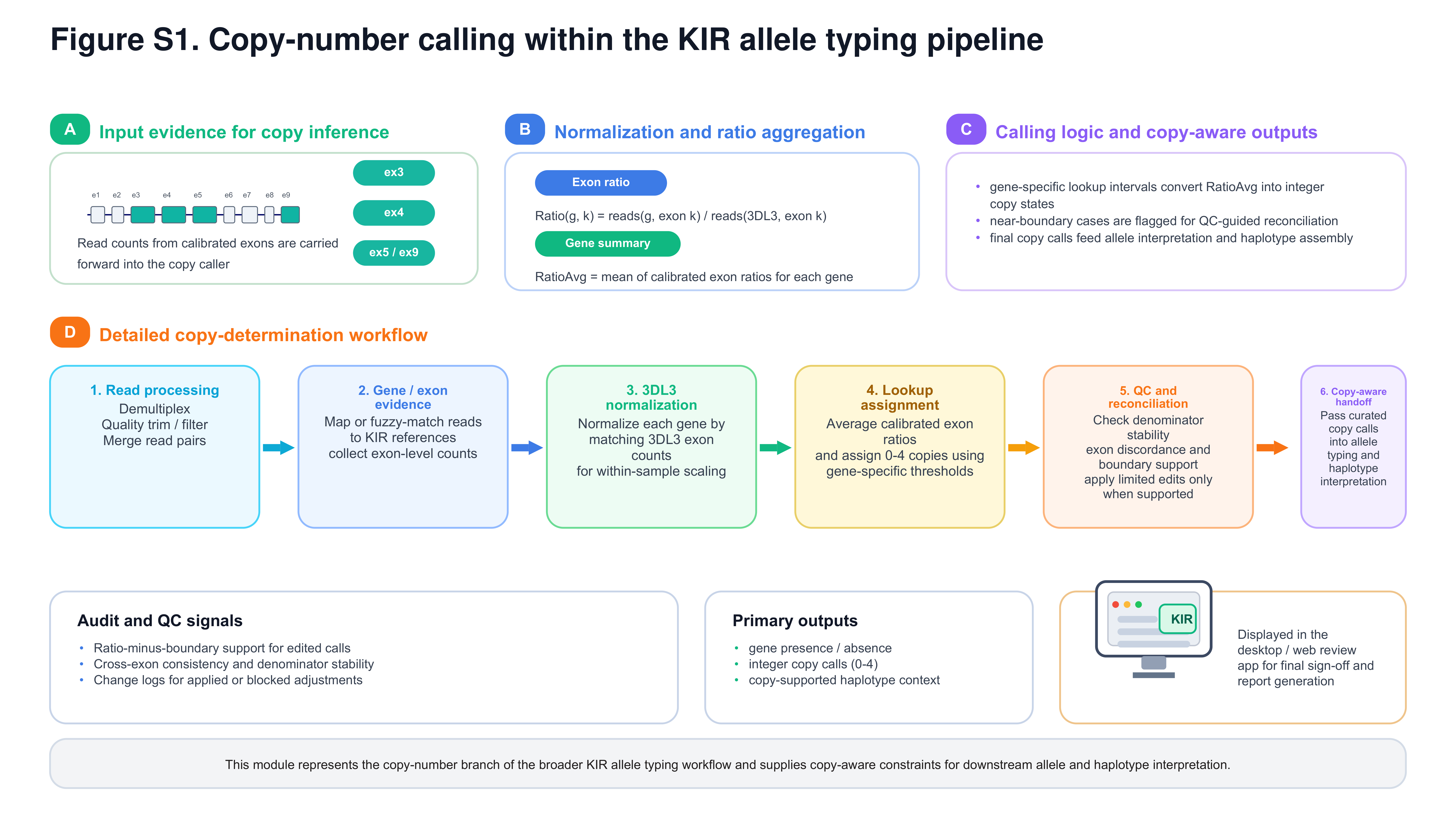

### FigureS2_KIR_cohort_and_richness.png

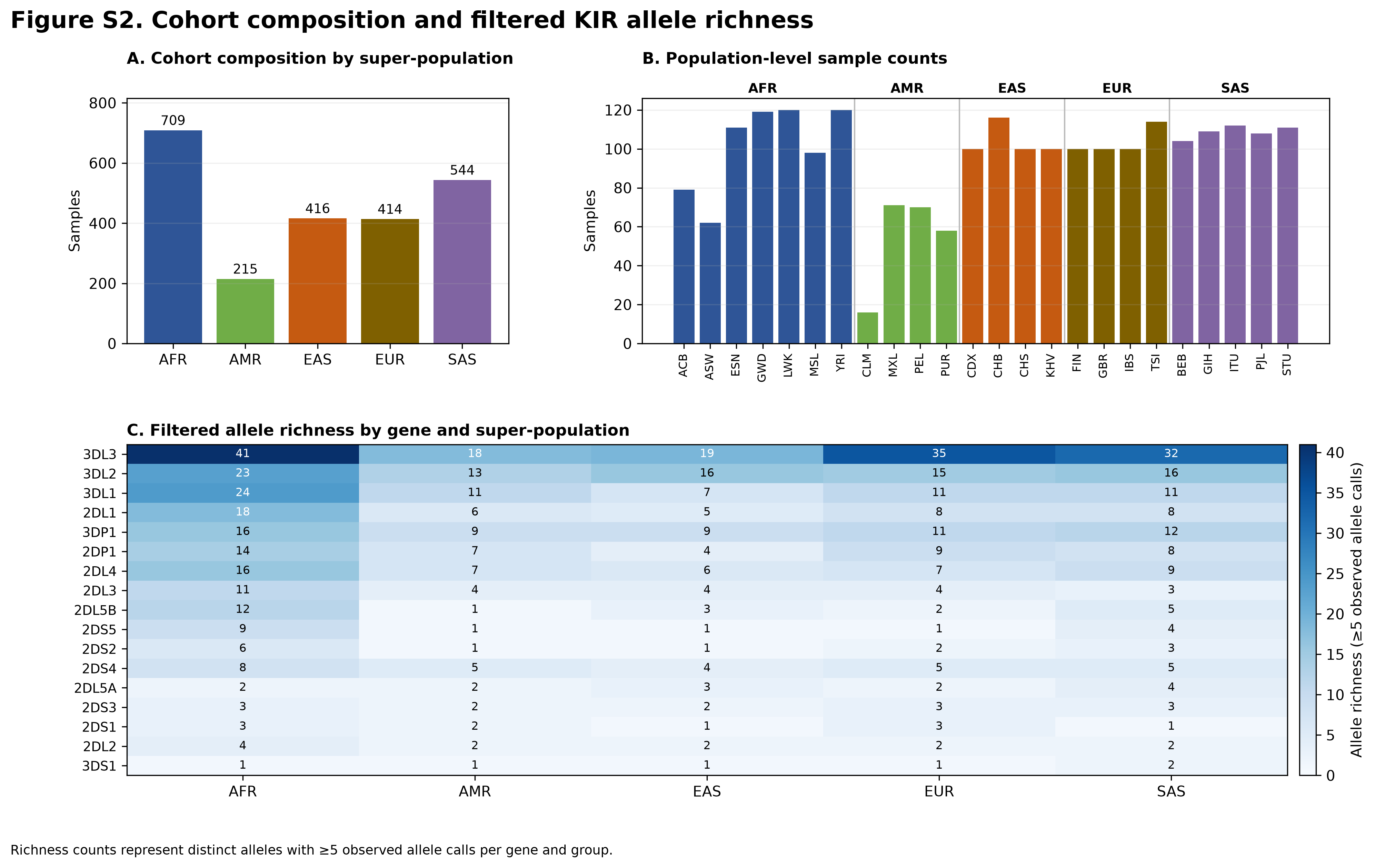

### FigureS3_KIR_population_richness.png

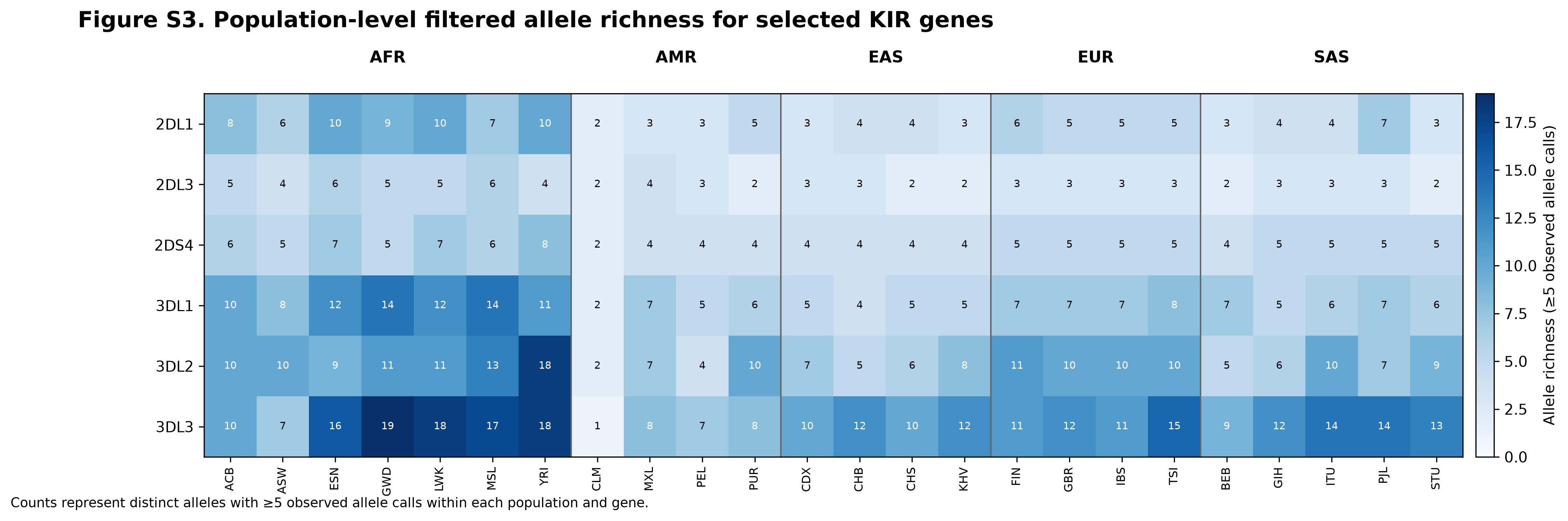

### FigureS4_KIR_exact_haplotype_architectures.png

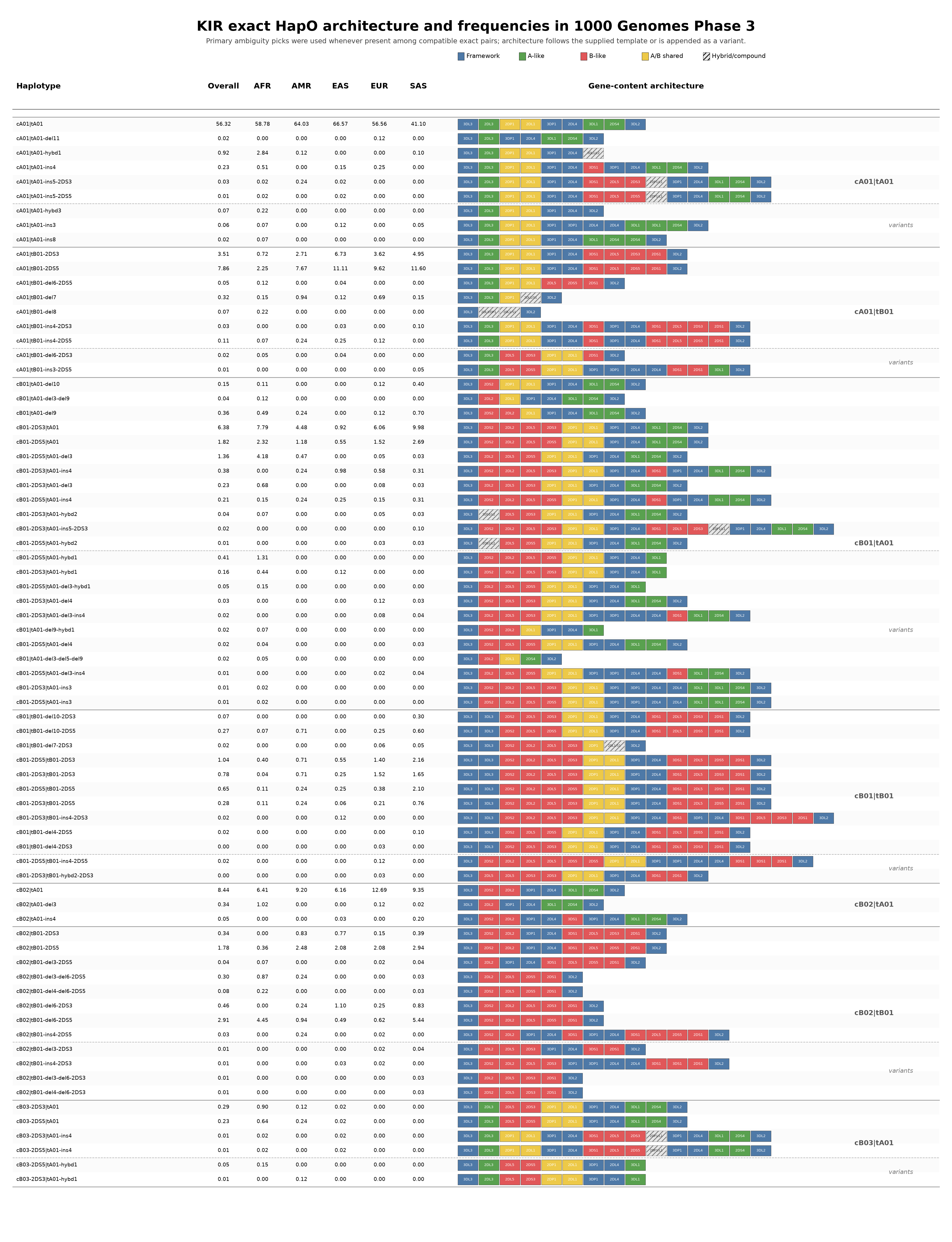

### FigureS5_KIR_collapsed_haplotype_frequencies.pdf

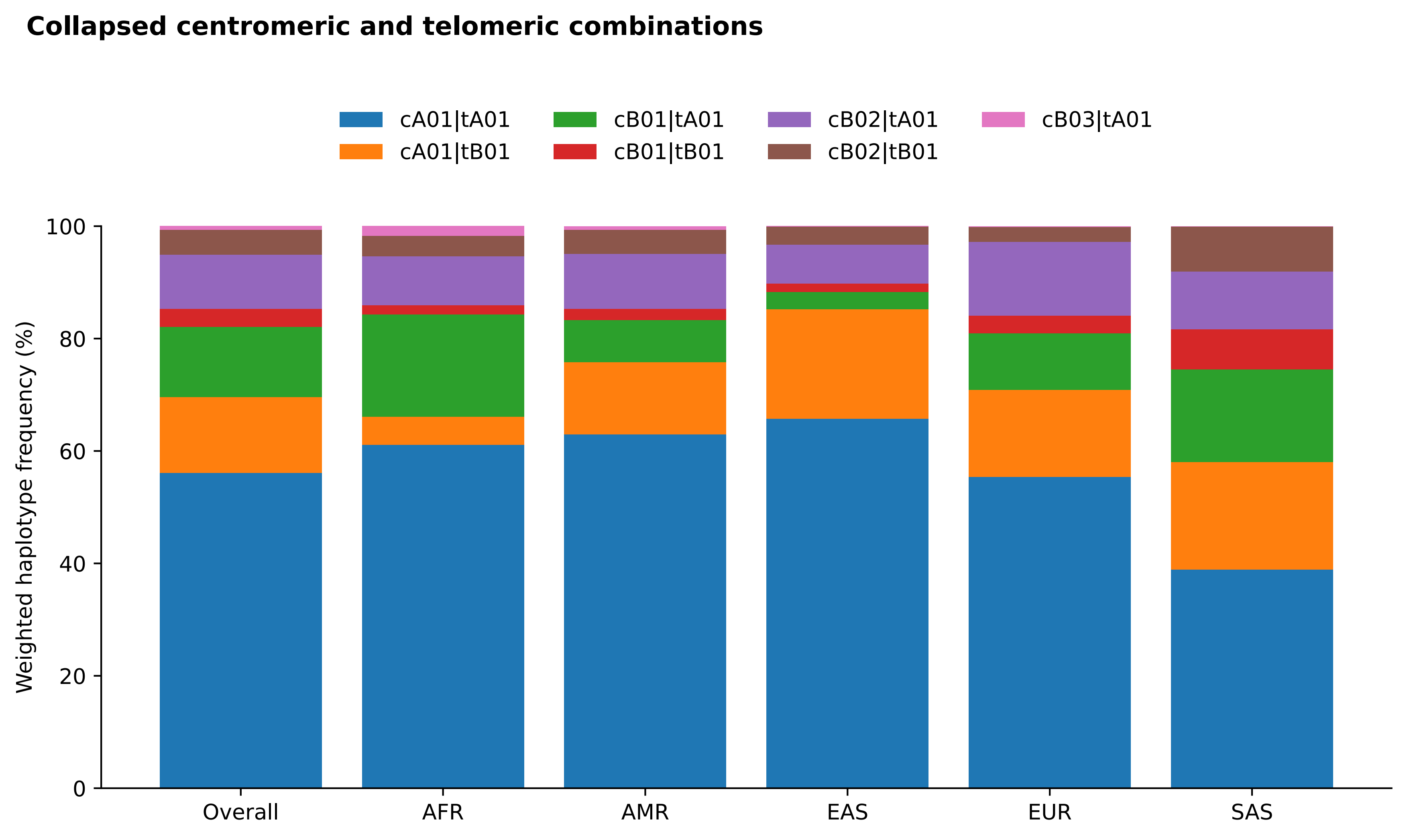

### FigureS5_KIR_collapsed_haplotype_frequencies.png

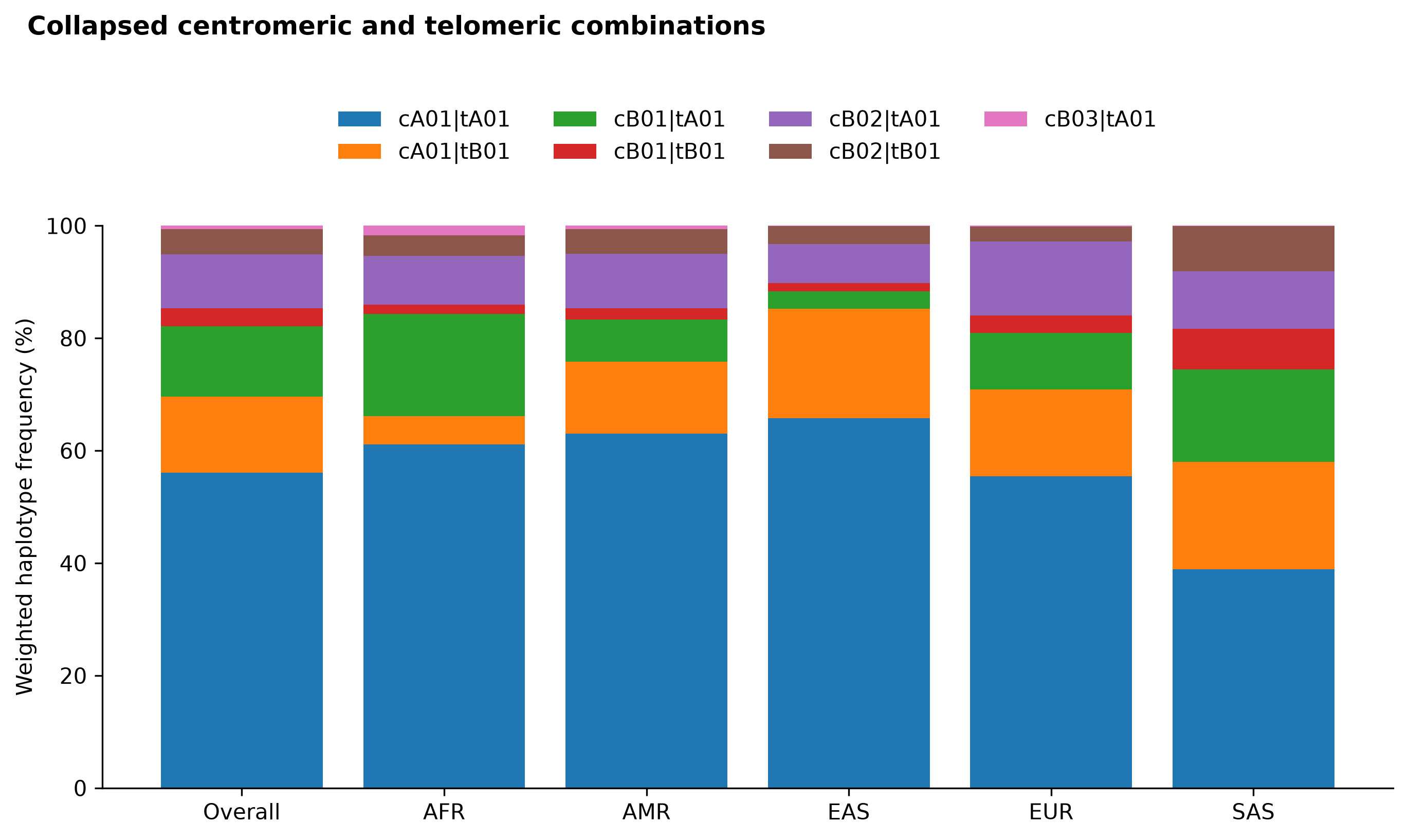

### FigureS6_KIR_motif_alleles_by_superpopulation.pdf

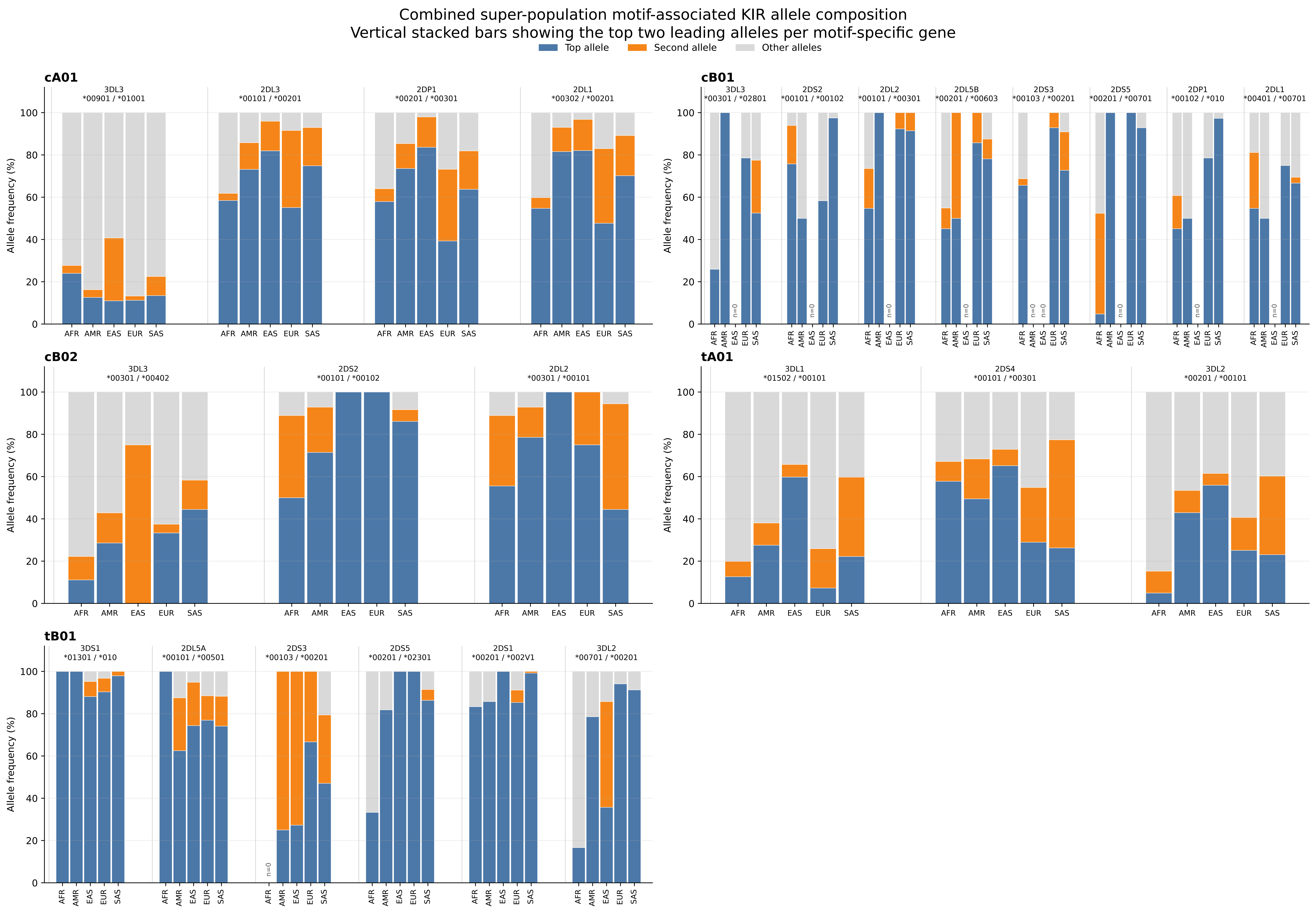

### FigureS6_KIR_motif_alleles_by_superpopulation.png

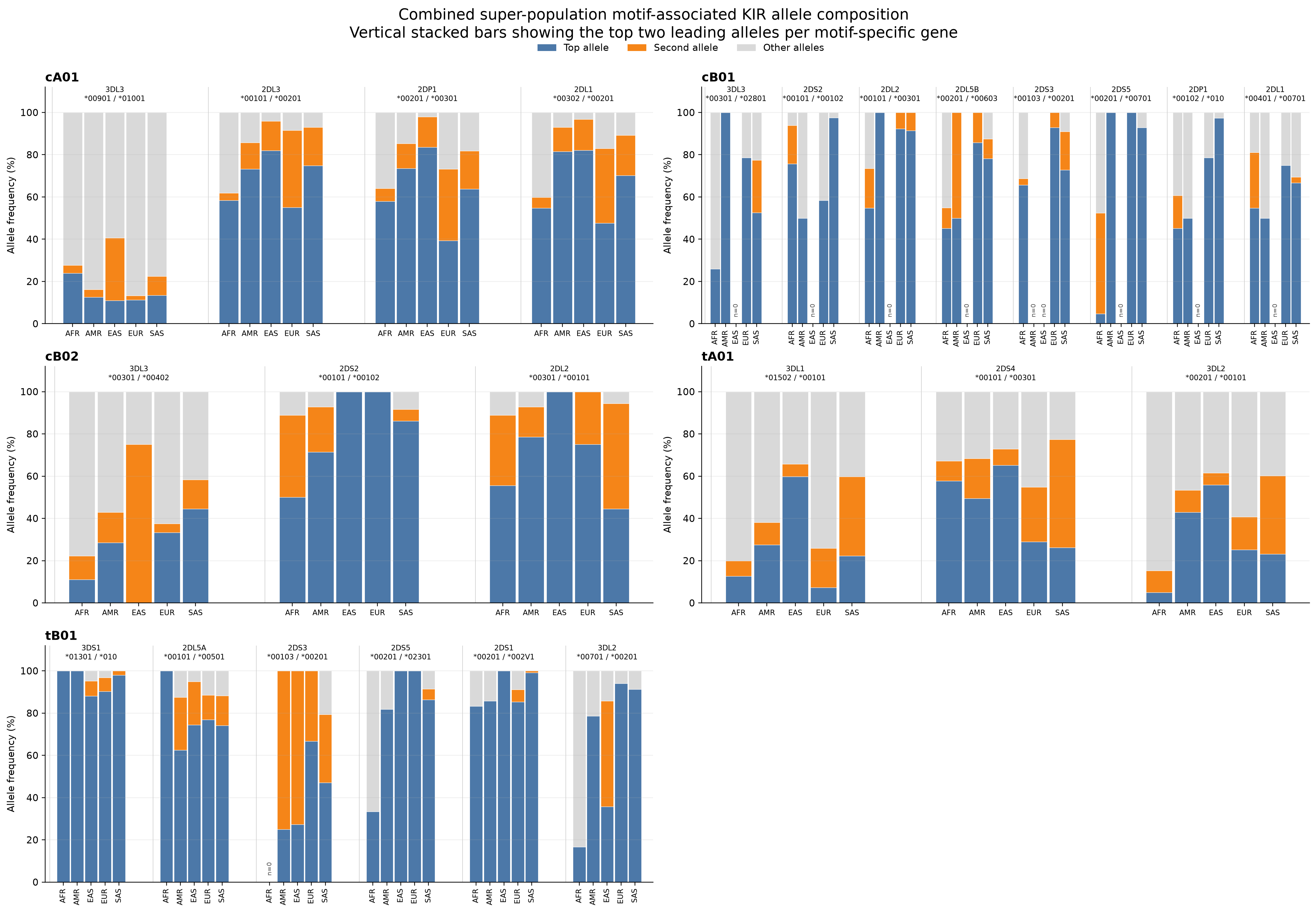

### FigureS7_KIR_HLA_population_support.png

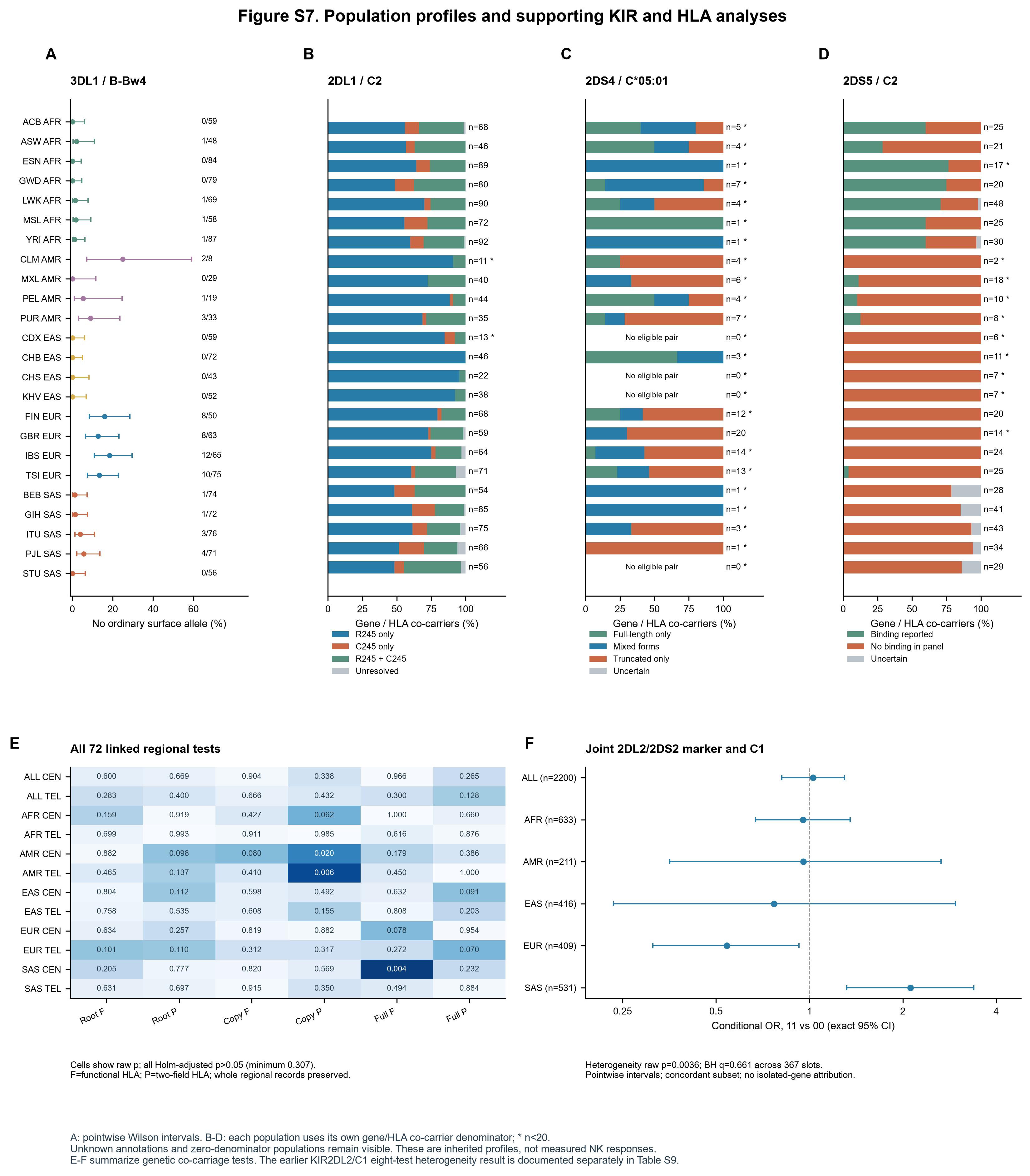

### FigureS8_KIR_HLA_association_calibration.png

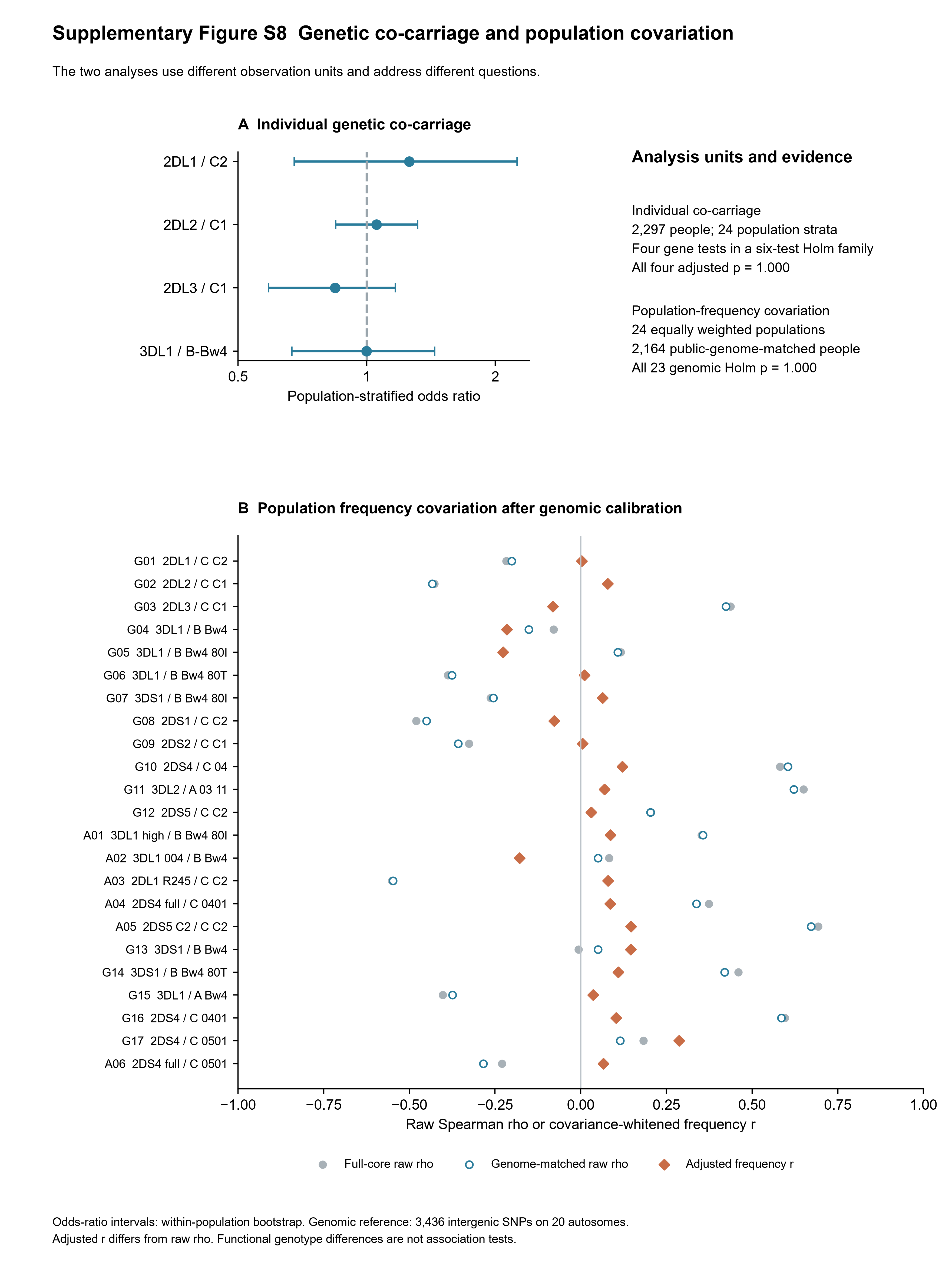
